# Shared and distinct sequence-function signatures define different modes of human TpoR activation

**DOI:** 10.1101/2025.09.09.675271

**Authors:** Xinyu Wu, Harry McLeod, Anna Malinovitch, Sally Hunter, Samyuktha Ramesh, Margareta Go, Julie V. Nguyen, Alan F. Rubin, Wessel A.C. Burger, Piers Blombery, Matthew E. Call, Melissa J. Call

## Abstract

The human thrombopoietin receptor (hTpoR) exists primarily as JAK2-associated monomers that become activated when converted to dimeric forms that support JAK trans-phosphorylation. This can be achieved by several different modes of stimuli, including the natural ligand Tpo, biologic agonists that bind the same site as Tpo, small-molecule drugs that bind the transmembrane (TM) domain, oncogenic mutations in and near the TM domain, and by association with constitutively active JAK V617F or a mutant form of the chaperone protein calreticulin. It is unclear how the dimeric structures induced by synthetic agonists and mutations relate to one another, and whether any of these induce the same active structure as the native ligand Tpo, yet this has important implications both for fundamental cytokine receptor biology and for development of targeted interventions for hTpoR-driven myeloproliferative diseases. Here we used deep mutational scanning (DMS) across the TM and juxtamembrane (JM) regions of hTpoR to extract feature-rich sequence-function signatures across a variety of different activating contexts. While each displayed some unique features, synthetic agonists and activating mutations all exhibited strong dependence on a common TM interface that is consistent with previous models of a left-handed, near-parallel helix dimer with H499 facing lipid. In contrast, Tpo-mediated activation was broadly insensitive to TM-JM substitutions, indicating that it does not rely on the same interface. Modeling with AlphaFold 3 (AF3) consistently yielded a right-handed, “splayed” helix dimer that is close at the extracellular face, contains H499 in the interface and diverges toward the cytosolic face, resting on an intracellular amphipathic JM helix that lies parallel to the membrane, which is also observed in a DMS/AF3 analysis of human erythropoietin receptor. This splayed Tpo-bound dimer could be stably inserted into a lipid bilayer with associated JAK2 using molecular dynamics and is supported by experiments showing that most or all of the TM domain can be replaced by poly-valine, with little effect on Tpo-driven activation but catastrophic effects on responses to synthetic ligands. Our data support at least two different structural modes of hTpoR activation that reconcile prior biochemical models, rationalize patient variants, and inform mechanism-based agonist and antagonist design.

## INTRODUCTION

Mutations that alter activity of the human thrombopoietin receptor (hTpoR/Mpl) can cause myeloproliferative disease, and the TpoR signaling axis is therapeutically modulated to counteract signaling defects in several important clinical settings (**Figure 1A**). During normal physiological activation, two TpoR chains come together around a single Tpo ligand to form an asymmetric dimer^1,2^ that signals through JAK2^3^, which docks via two conserved motifs (Box 1 and Box 2) in the TpoR cytoplasmic tail^4^. A simple model of triggering posits that Tpo binding dimerizes TpoR^5^, leading to close proximity of JAK2 monomers that cross-phosphorylate to activate one another and initiate signaling. However, dimerization of TpoR is not sufficient for JAK2 activation, because engineered coiled-coil fusion receptors and cysteine crosslinking studies have identified both active and inactive dimeric orientations of their transmembrane (TM) domains^6–8^.

**Figure 1.**
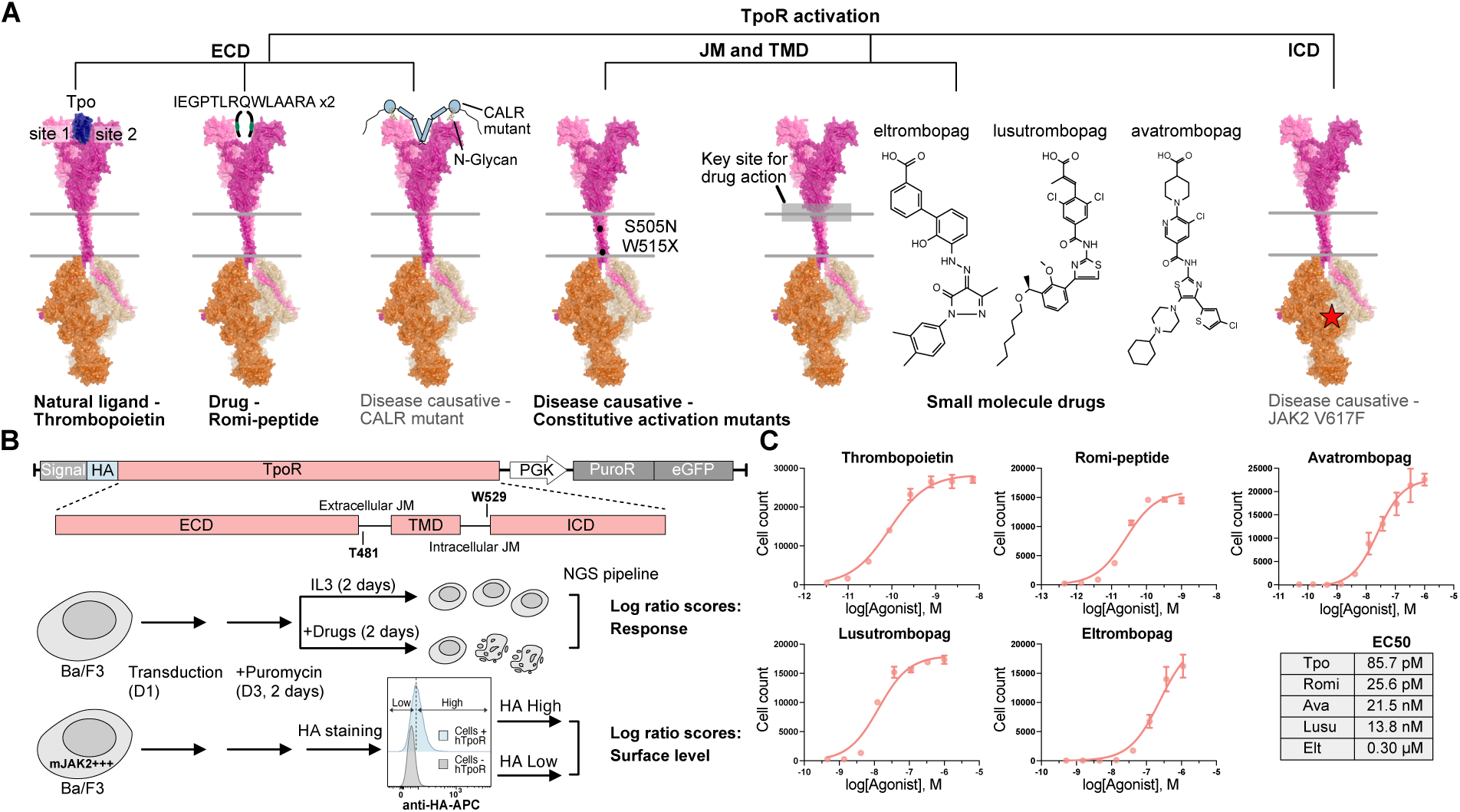
Different modes of hTpoR activation and outline of DMS workflow. **(A)** Modes of human TpoR activation and the site of action of each type of stimulus on the receptor. The hTpoR is shown in pink, JAK2 is shown in orange. Stimuli described in grey text below the images were not examined in this study. **(B)** Retroviral vector structure and DMS workflow. hTpoR (red) includes an N-terminal HA tag and is followed by a puromycin resistance-eGFP fusion. The segment T481-W529 spans the juxtamembrane (JM) and transmembrane (TM) domains and represents the sequence covered by DMS libraries. Below the vector design is a schematic overview of the DMS workflow, including two selection arms for activity on agonists (top) and sorting for surface expression levels (bottom). **(C)** A representative dose response on WT hTpoR for each type of agonist used in this study. The x-axis displays the agonist concentration on a logarithmic scale and the y-axis shows raw cell count. The EC50 values in the table represent the mean of two biological replicates. Error bar: mean ± SD of three technical replicates.

The structure of the TM and adjoining juxtamembrane (JM) regions during receptor activation are of high interest because of the many ways the TM domain can be manipulated to activate the receptor (**Figure 1A**). There are more than a dozen mutations in the TM domain that induce constitutive activation of hTpoR^9^, with S505N^10–12^ and a variety of mutations at W515^13–15^ regularly found in patients with essential thrombocythemia. Additionally, several small-molecule drugs activate hTpoR by directly engaging the TM domain (**Figure 1A**), and these are used to boost platelet levels in various clinical settings. The first of this class of molecules, eltrombopag^16^, was developed from a screen for hTpoR agonists in Ba/F3 cells^17^, a widely employed model system for JAK2-dependent cytokine receptor studies. Eltrombopag requires H499, a TM feature unique to hominid TpoRs, and NMR studies with a precursor compound indicate that this is the drug binding site^18^. Development of additional small-molecule agonists avatrombopag^19^ and lusutrombopag^19^ followed, and these also activate the receptor by engaging H499. Finally, a dimeric fusion protein of a Tpo peptide mimic, romiplostim, activates TpoR by binding to the same extracellular site as the natural ligand^20^. It is believed that all of these mutations and therapeutic agonists activate the receptor by inducing dimerization, but experimentally determined structures showing precisely how this translates to intracellular JAK2 transphosphorylation are not yet available.

Structural changes in both TM and JM regions have been implicated in various TpoR activation scenarios based on biochemical and biophysical studies of receptor fragments^6,18,21,22^, cysteine crosslinking of intact receptors^6,7^ and a great deal of exploratory engineering and mutagenesis. These studies point to specific dimeric interfaces within the TM domain as well as potential transitions between helical and non-helical structure in the extracellular and intracellular JM regions. TM-binding drugs and activating mutations may have common structural underpinnings^23^, while the oncogenic V617F mutant of JAK2 induces an active TM conformation that may be distinct from these^7^. How any of these relate to the native Tpo-induced structure is yet unclear.

In this study, we took a comprehensive mutagenesis approach to map the signatures of sequence dependence in the TM-JM region, as indicators of their mechanisms of action, for four different classes of activating stimuli: the native ligand Tpo, three small-molecule agonists that act on the TM domains, a peptide based on the biologic agonist romiplostim (here called romi-peptide), and the constitutively active clinical variants S505N and W515K. We employed deep mutational scanning (DMS) to interrogate the requirements for hTpoR surface expression and activation spanning a region that begins in the last structured module of the extracellular domain, extends through the TM domain, and ends at the beginning of the Box 1 motif that docks to the JAK2 FERM domain (**Figure 1B**). Surprisingly, we found that Tpo can effectively stimulate nearly every variant that supported surface expression. In stark contrast, the TM-acting drugs, activating mutation S505N and romi-peptide had signatures indicating strong dependence on extracellular JM and TM domain interfaces, while the activating mutation W515K had a largely unique signature at the intracellular JM region. Viewed in the context of existing and new receptor dimer models, these shared and distinct signatures illuminate the different ways that TpoR can be activated and suggest that native and non-native stimuli can activate the receptor via distinct structural mechanisms.

## RESULTS

### Assay design and library construction

As in previous work^9^, we used the cytokine dependence of Ba/F3 cells to discriminate active from inactive hTpoR variants. Three different DMS libraries were constructed covering amino acid positions T481-W529 (**Methods; Supplemental Figure 1**) on the hTpoR wild-type (WT), S505N or W515K backgrounds, and these were installed into a retroviral vector with a GFP-puromycin resistance cassette (**Figure 1B**). Libraries contained around 1000 protein variants, each with one single-amino-acid substitution. Dose response assays were performed with all agonists on Ba/F3 cells expressing WT hTpoR (**Figure 1C**) and yielded EC50 values similar to those previously reported^1,16,19,20,24^. From these curves, we determined target concentrations to perform DMS screens using high (EC95) and low (EC30) agonist concentrations for selection.

### Characterization of Tpo response and hTpoR surface expression

We screened our WT-background hTpoR library for Tpo response by treatment at high and low concentrations for two days (**Figure 1B**), with maintenance on IL-3 as the non-selective reference. Under these conditions, variants responding to Tpo are enriched and non-responsive variants are depleted. The heatmaps in **Figure 2A-B** clearly show that all nonsense variants (top row) were impaired, as expected. The TM domain is predicted to fall between W491 and W515 (UniProt entry P40238), and except for H499 we saw a clear requirement for amino acids that confer stability in the membrane at a subset of these positions (T496-R514). Unexpectedly, hTpoR responsiveness at EC95 was unaffected by substitution of residues W491-V495 with polar or charged residues, suggesting that this segment may not need to be fully embedded in the membrane to be receptive to activation by the native ligand.

**Figure 2.**
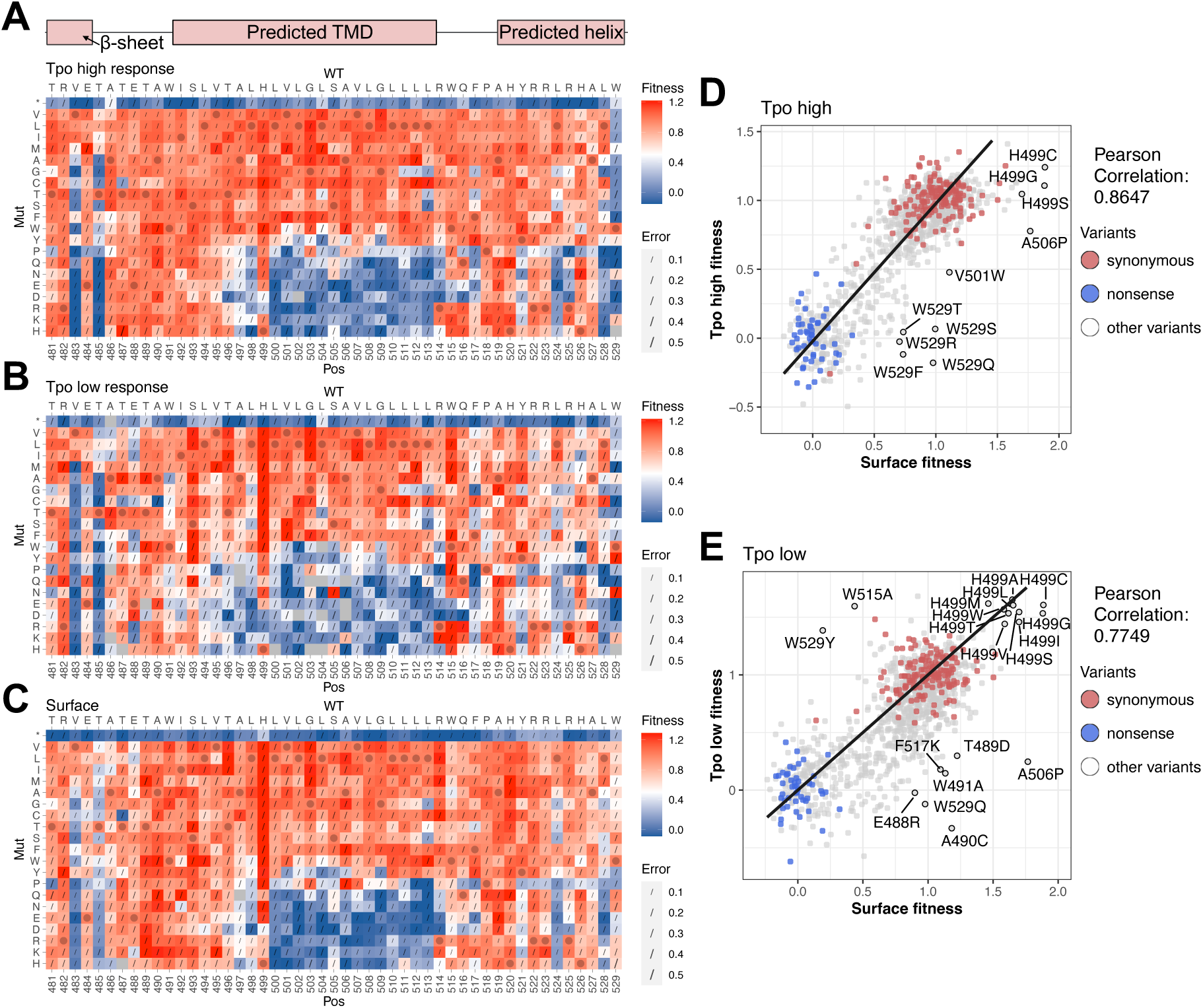
DMS analysis of Tpo response and hTpoR surface expression. **(A)** DMS sequence-function heatmap showing the response of hTpoR variants to high Tpo concentration (EC95). Predicted secondary structure is illustrated above the heatmap. Columns show consecutive amino acid residues of the hTpoR protein sequence from the N-terminal (left) to C-terminal (right) direction for the region scanned in this study, with native sequence at the top and position number at the bottom. Rows indicate the substituting amino acids with “*” referring to premature stop codons. Each box depicts the normalized fitness score in a three-color gradient: blue indicates impaired response, red indicates WT-like response, white indicates intermediate response. WT amino acids at each position are indicated with a dot. Errors (derived from “Sigma” values reported in DiMSum) are indicated with slashes with the length proportional to magnitude. Missing variants are shown in grey. **(B)** DMS heatmap showing the response of hTpoR variants to low Tpo concentration (EC30). Details as in (A). **(C)** DMS heatmap showing surface levels of TpoR variants. Blue indicates variants with impaired surface levels, red indicates variants present at WT-like levels, white indicates intermediate levels. All other details as in (A-B). **(D-E)** Scatter plots of Tpo response scores (y-axes) at high (EC95, panel D) and low (EC30, panel E) concentrations against fitness scores reflecting surface expression levels (x-axes), with each variant in the library represented by a dot. A straight line connects the clusters corresponding to synonymous-WT (red; centered at 1,1) and nonsense variants (blue; centered at 0,0), while other variants are shown in gray. The Pearson correlation score is displayed on the right side of the plot. Visually identified examples of variants where activity does not reflect surface expression and some H499 variants are labelled with an open circle and variant ID.

Many variants will exhibit defective responses due to poor surface expression, so to focus our attention on variants whose altered activity could not be explained by altered expression, we screened again by surface staining and flow-sorting into hTpoR surface-high and hTpoR surface-low groups (**Figure 1B**). This surface abundance data (**Figure 2C**) exhibited a striking concordance with activity data (**Figure 2D-E**): almost every variant that was impaired in activity was also impaired in surface abundance, most due to polar substitutions in the hydrophobic TM domain. Also consistent with activity data, polar-substituted variants at W491-V495 expressed well, indicating that this part of the predicted TM domain does not need to be embedded in the plasma membrane for stable receptor expression. Variants at W529, at the start of the cytoplasmic Box1 motif, generally had good surface expression but poor activity. These variants likely bind JAK2 but are unable to activate it. Variants with hydrophobic substitutions at H499 were more abundant at the cell surface compared to WT TpoR, but this only translated into an increased growth/survival benefit at low Tpo concentration (**Figure 2E**). We note the especially strong surface expression defects associated with mutations at V483 and T485, which form part of the final β-sheet of the membrane proximal fibronectin domain, and with charged/polar substitutions at Y521, L524 and L528 in the cytoplasmic tail (**Figure 2C**). The former likely causes folding defects, while the helical periodicity of the latter indicates the existence of a short helical segment in the intracellular JM region between the RWQFP motif^21^ and the beginning of the Box 1 motif. Overall, given the high concordance of Tpo response and surface expression, these data do not provide evidence of an extensive TM interface in the receptor that is absolutely required for Tpo-induced signaling.

### Features of the JM/TM that affect activation with synthetic agonists

In contrast to our results with Tpo, DMS screens in the presence of synthetic agonists revealed strong signatures of sequence dependence in and near the membrane that could not be accounted for by surface level changes, at both high and low concentrations (compare heatmaps in **Supplemental Figure 2** to surface expression in **Figure 2C**). As expected, all small-molecule drug treatment groups were severely impacted by mutations at H499 (**Supplemental Figure 2C-H**), and functional enhancement was observed for the strong constitutively active mutants S505N and W515X, particularly at low agonist concentrations (**Supplemental Figure 2B, 2D, 2F, 2H**). In addition to these expected outcomes, we saw complex patterns in each dataset and sought to reduce the high-dimensional data to classify mutations based on shared features. UMAP clustering^25^ (parameters in **Methods**) on fitness scores from the TpoR surface screen and from high and low concentrations of all agonists (11 data sets in total) organized variants into five distinct clusters (**Figure 3A**). For ease of visual interpretation, the locations of variants in each cluster were separately mapped back onto sequence-function grids (**Figure 3B-F**) and color-coded to the relevant UMAP cluster in panel A.

**Figure 3.**
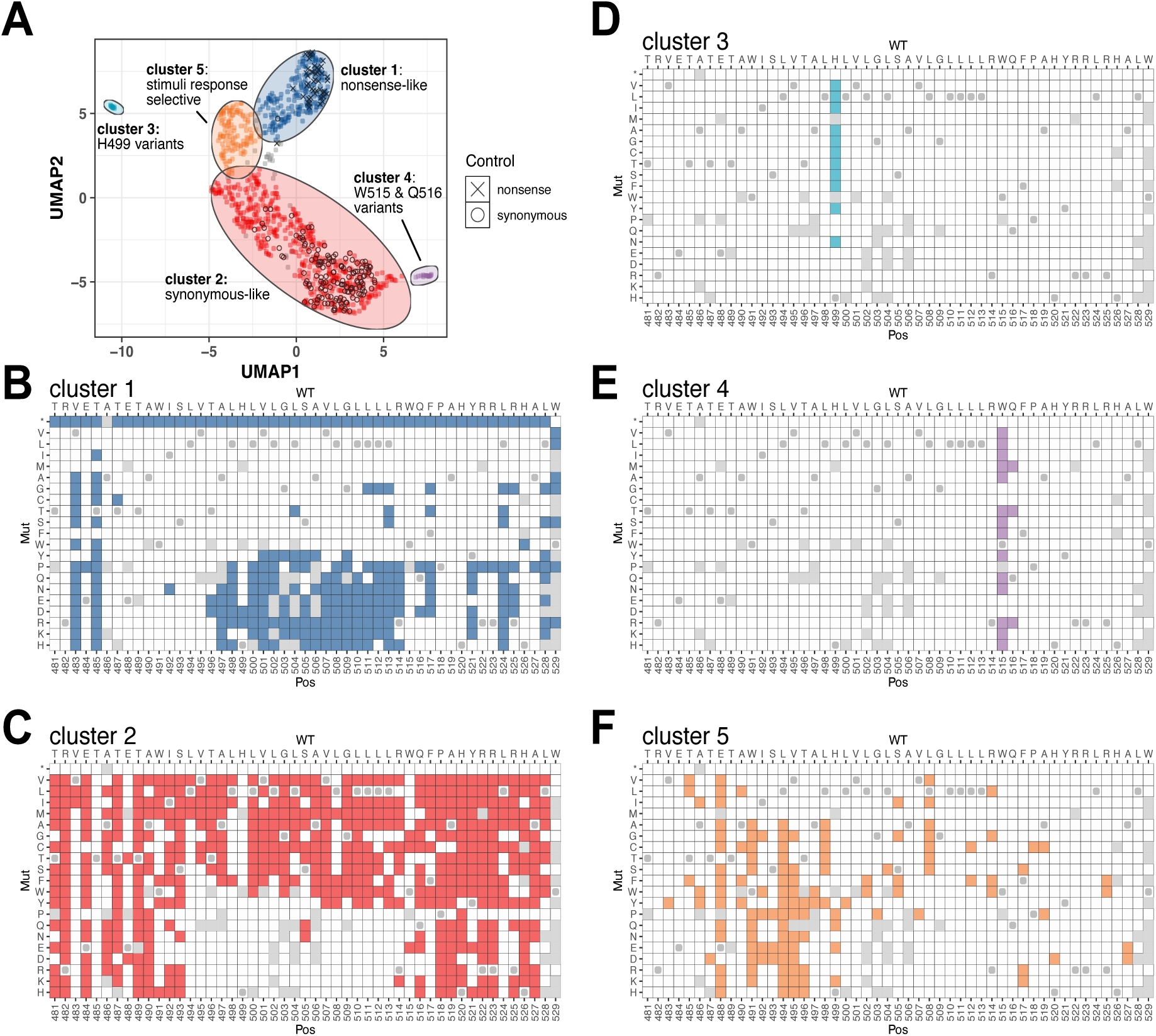
UMAP cluster analysis of DMS datasets. **(A)** UMAP analysis of the TpoR ligand response datasets (10 treatment groups) and the surface levels dataset (11 in total). Nonsense variants are displayed as crosses, synonymous-WT as open circles, and all other variants as colored dots. **(B-F)** Variants populating each cluster are mapped back onto the heatmap grid structure to illustrate their locations in the scanned region. Solid colors are matched to the UMAP in panel (A) but color/shade is not representative of any quantitative score. Missing variants are shown in grey. Variants not part of the cluster are shown in white. The WT amino acid at each position is marked with a dot. Grey variants from the center of the UMAP in (A) were not part of any cluster and are mapped back onto the grid structure for reference in **Supplemental Figure 3**.

Cluster 1 (blue) comprises variants that were inactive in all treatment groups, including both nonsense and missense variants that prevented surface expression (**Figure 3B**). There were some additional mutants in this cluster that were partially impaired in both activity and surface levels, such as helix-disrupting proline substitutions throughout the TM domain and in the intracellular JM helix. Cluster 2 (red) contained variants that were active in all conditions, behaving like WT TpoR or constitutively active mutants (**Figure 3C**). Variants in remaining clusters had differential effects across stimuli: cluster 3 (cyan) contained only H499 substitutions that prevented drug activity and increased receptor expression (**Figure 3D**), while cluster 4 (purple) contained only variants at positions W515 and Q516 (**Figure 3E**). Variants at W515 endow the receptor with constitutive activity^9,13^, while variants at Q516 do not^9^, but the latter appeared to uniquely enhance TpoR small-molecule drug responsiveness. Cluster 5 (orange) contained variants that were deficient in responsiveness to all three small-molecule drugs but maintained Tpo, and sometimes romi-peptide, responsiveness (**Figure 3F**). These were most concentrated in the extracellular JM-TM region (E488-V496), but variants at multiple positions throughout the TM domain were also impaired. The small group of variants that were not assigned to a cluster (grey) were scattered throughout the variant space and did not reveal any clear pattern (**Supplemental Figure 3**).

### Comparison of prominent features from each treatment group with cluster analysis

Visual inspection of the sequence-function maps suggested that each cluster likely contained multiple overlaid sequence-dependency “signatures” with features that are more prominent in some treatment conditions and less so in others. To identify the most prominent features of each stimulus for comparison to the cluster maps, we plotted variant activity fitness scores against TpoR surface fitness scores and isolated variants that had significantly higher or lower activity than can be explained by surface expression level alone (**Figure 4A-E**, EC95 data; **Supplemental Figure 4A-E,** EC30 data). These were again mapped back to the variant grid for ease of interpretation, this time with color shading indicating the degree of deviation from the trendline in the direction of more (orange) or less (green) activity than expected. This analysis again revealed the overall lack of effects on the Tpo response for most variants (**Figure 4A**). This also allowed us to systematically compare data from separate screens for constitutive activity using the variant libraries we made on the S505N and W515K backgrounds (**Figure 4F-G, Supplemental Figure 2I-L**), which were not compatible with our cluster analysis owing to their unique base sequences. As expected, H499 variant effects in cluster 3 were attributable to eltrombopag, avatrombopag and lusutrombopag datasets, while romi-peptide, S505N and W515K treatment groups were less prominently affected. Cluster 4, containing W515 and Q516 variants, was dominated by enhancing effects in eltrombopag, avatrombopag and lusutrombopag treatment groups, and this feature was shared with S505N but not W515K.

**Figure 4.**
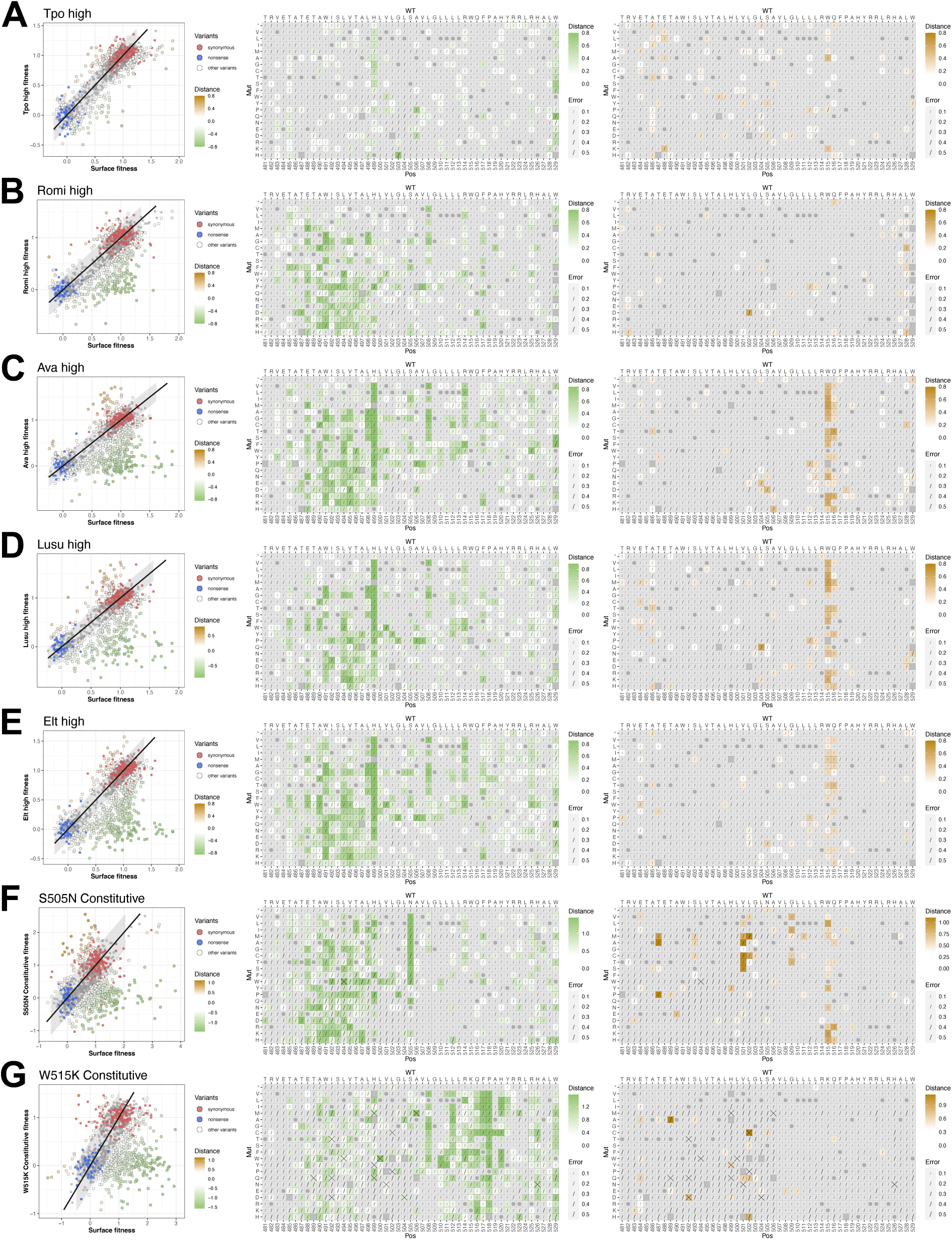
Identification of sequence-function signatures for agonists and activating mutations. Each panel depicts the departure from the correlation between activity and surface scores, as described in Figure 2D, for agonists at EC95 **(A-E)** or disease-associated activating mutations **(F-G)**. The shaded ribbon on the scatter plots was calculated from the spread of synonymous wildtype and nonsense variants and encompasses 2 SD. Outlier variants below the ribbon are functionally impaired and are displayed on the sequence-function maps (middle) in a gradient of white to green indicating their distance from the straight line. Variants above are highlighted in gold indicating the degree of enhanced activity and are displayed on sequence-function maps (right). Errors indicated by slashes were calculated as the square root of the sum of squared surface expression and activity errors, and variant scores with an error over 0.5 are marked with a cross. The corresponding residual plots and maps for EC30 treatments are provided in **Supplemental Figure 4** for comparison.

Signatures of all non-native stimuli had significant overlap with variant cluster 5, which revealed particular sensitivity to polar substitutions in the extracellular JM-TM region and a mostly helical pattern involving W491, L494, V495, L498, S505, L508, L512 and R514 dependency in the TM domain. All non-Tpo agonists, including romi-peptide, were strongly impacted by mutations that decreased hydrophobicity above L498, and all were dependent on L508. Small-molecule drugs and W515K were negatively impacted by R514 mutations, but the effects of these mutations on romi-peptide and S505N were weak. W515K displayed a strong dependence on L512 and was uniquely sensitive to mutations at Q516, F517, and P518, some of which also enhanced responses to small-molecule drugs. That such signatures are weak or absent under Tpo stimulation at both low and high concentrations (**Figure 2; Supplemental Figure 4A-B**) indicates that most of these features are uniquely required for responses to non-native agonists and constitutive activity driven by oncogenic mutations.

### Analysis of single variants displaying stimulus-selective effects

From the cluster 5 variants that negatively impacted responses to all agonists except Tpo, we chose two positions that were intolerant to almost every alteration and remade constructs as traditional single variants for more extensive analysis. In the TM domain, we compared L508I and L508V variants with WT TpoR in dose-response tests against all agonists (**Figure 5A-B**). For both variants, we saw significant impairment on romi-peptide and near-complete loss of response to small molecules, but only very modest effects on Tpo. This result confirms the DMS data reflecting a stringent requirement for leucine at position 508, and it suggests a highly specific contact that is likely to be far from the agonist binding sites^18,20^. In the extracellular JM region, we selected E488N and also confirmed its highly specific impairment on synthetic agonists but not Tpo (**Figure 5C**). Combined with the observation from our DMS screens that E488D is well tolerated in all treatment groups (**Supplemental Figure 2**), this latter result shows that negative charge is particularly important at this position and that the effect is not unique to the small-molecule drugs that likely bind in its vicinity^18^.

**Figure 5.**
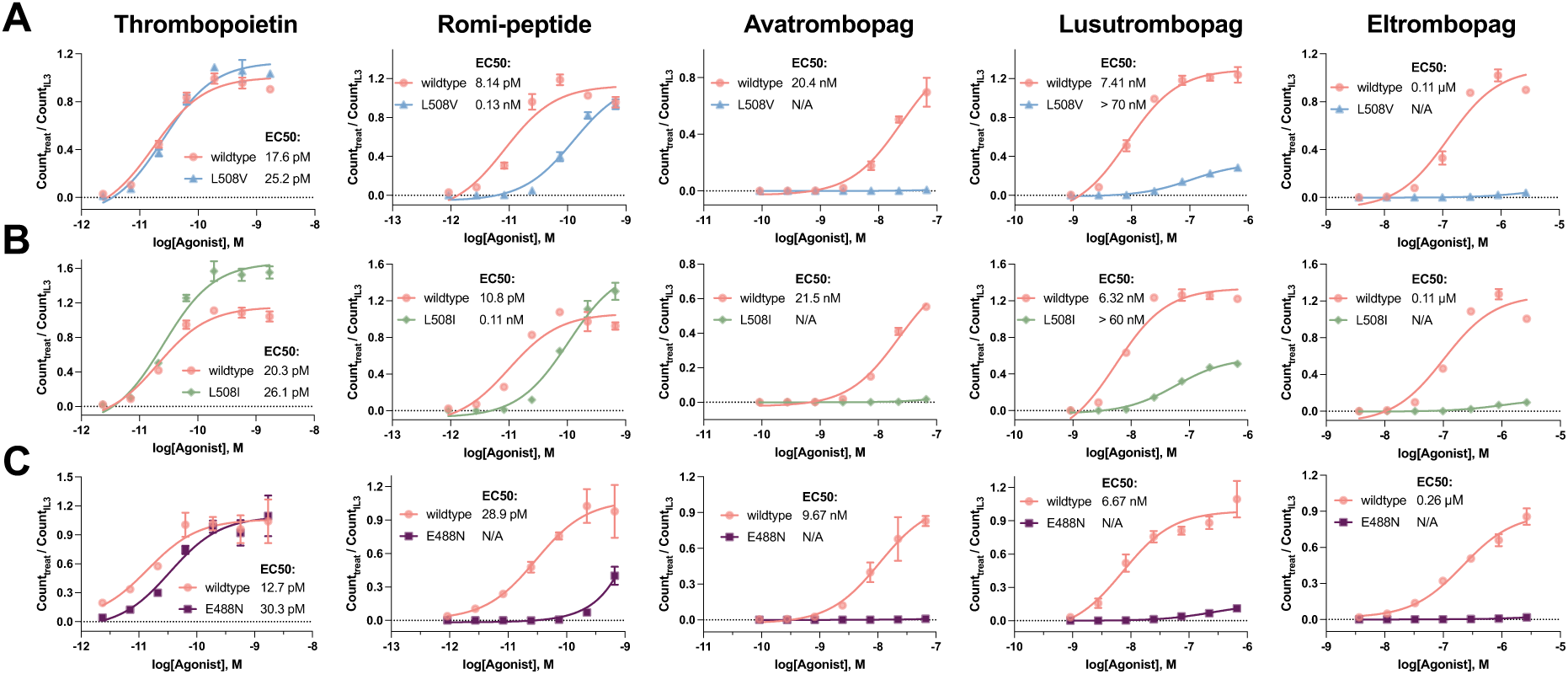
Validation of example stimulus-specific single variant effects. Single-variant responses to Tpo or the indicated synthetic agonists for variants L508V **(A)**, L508I **(B)** and E488N **(C)**. Cell counts were normalized by dividing the cell counts from each well by those from IL3 treated cells. Mean ± SD of three technical replicates is shown for one representative experiment and EC50 values were calculated as the mean of two biological replicates.

### Alignment of DMS data with models of activated receptors

The available cryo-EM structures of mouse and human Tpo-bound TpoR^1,2^ do not contain the membrane-associated portions. However, a recent experimentally guided modeling study provides an instructive template for contextualizing our deep mutagenesis data in the TM domain. The Tpo-and JAK2-bound hTpoR model generated by Pogozheva et al^26^ using AlphaFold Multimer aligns well with experimentally determined ligand-bound structures of the mouse and human TpoR ectodomain dimers^1,2^ and the active JAK1 dimer^27^. This model (**Figure 6A-B**) features an extensive left-handed TM helix dimer interface that has contacts at nearly all the positions identified as functionally important for synthetic agonist and oncogenic mutation activity in this study and others. Importantly, it leaves H499 in an outward-facing position where it can be bound by small-molecule agonists^18^ and features a short amphipathic helix in the intracellular JM region, as also predicted in earlier studies^28,29^. The close packing at the extracellular (top) end of the TM domain is evident in our DMS data showing high sensitivity to large aromatic substitutions at A490 and small substitutions at W491 (see **Figure 4B-F**, green variants), with the latter also identified in another recent study^23^. The branched aliphatic packing below this is supported by detrimental effects of large aromatic and small substitutions at L494, V495 and L498 (again see **Figure 4B-F**). Additionally, L498, V501, L502, S505 and L508 (**Figure 6B**, orange) are all sites where specific polar substitutions can drive or enhance constitutive activity associated with myeloproliferative diseases^9,11,14,21,23,30^, likely via inter-helical hydrogen bonds^31,32^. L508 and L512 show two different signatures of close complementary packing in the DMS data, with L508 tolerating very few TM-compatible substitutions and L512 particularly sensitive to large aromatic substitutions (see *e.g*., **Figure 4C**). These are also the sites of unique crosslinks recently identified in JAK V617F-activated TpoR dimers by cysteine scanning mutagenesis^7^. Interestingly, synthetic agonists, but not activating mutations, exhibit a peak of sensitivity to proline substitutions at positions 506 and 507 (compare **Figure 4C, D, E** with **4F**), indicating that maintenance of ideal helical geometry is especially important for translating binding of these ligands through to the intracellular end of the TM dimer. Finally, the placement of W515 and Q516 in this model (**Figure 6B**) suggests that the enhancing effects of changes at Q516 we observed on small-molecule agonist and S505N-induced activity (see **Figure 4C-F**) could be due to loss of a trans-interaction that weakens the gating function of W515^21^, but Q516 mutations by themselves are not sufficient to drive constitutive activation.

**Figure 6.**
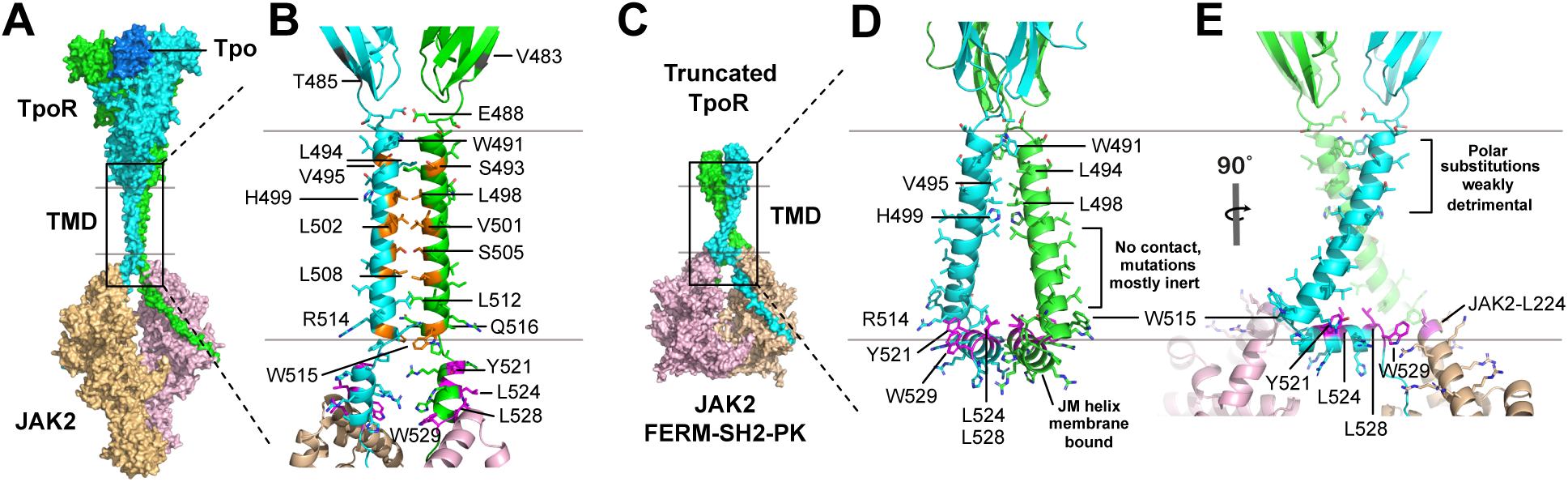
Alignment of DMS data with models of activated hTpoR. **(A)** AlphaFold Multimer-generated model of full-length activated Tpo-hTpoR-JAK2 from Pogozheva et al^26^. Gray lines indicate approximate membrane boundaries. Boxed region is enlarged in **(B)** with selected amino acids labeled. Positions of activating mutations are highlighted in orange and hydrophobic residues in the JM amphipathic helix are highlighted in magenta **(C)** AF3-generated model of hTpoR membrane-proximal fibronectin (D4) domain with TM and intracellular tail bound to JAK2 FERM-SH2-PK. Boxed region is enlarged in **(D-E)** with JAK2 removed in (D) for clarity. Hydrophobic residues in the JM amphipathic helix are highlighted in magenta and in (E) JAK2 L224 is highlighted in light purple. Basic residues in the JAK2 FERM domain that would be near the membrane surface are shown in stick representation.

While this model is largely consistent with the DMS data for synthetic agonist and constitutive mutant hTpoR activity, our screens on the native ligand Tpo revealed little evidence of strong dependence on any specific TM-TM interactions (**Figure 2**, **Figure 4A**) and thus do not support this model as the predominant Tpo-bound active structure. However, in the low-concentration (EC30) Tpo treatment condition (**Figure 2B and E**; **Supplemental Figure 4A**) there was a faint signal in the extracellular JM-TM region that bore some similarity to the pattern of mutations that reduced synthetic agonist and S505N activity. AlphaFold-based tools have continued to rapidly evolve, and so we used the latest iteration AlphaFold 3 (AF3)^33^ to explore whether there are other plausible models that are more consistent with the Tpo DMS data. Our attempts to model the entire Tpo-TpoR-JAK2 complex failed to give reasonable structures, because ectodomains invariably formed interfaces with JAK2. We therefore trimmed the ECD to the membrane-proximal fibronectin domain, removed the JAK2 kinase domain and repeated the predictions, reasoning that the membrane-proximal fibronectin domains would provide bulk to restrict the conformational landscape accessible for the TM-JM sequences, while the pseudokinase domain in JAK2 would enforce dimerization. Four AF3 runs were performed with different seeds, each outputting 5 structures (**Supplemental Figure 5**). Inspection of all structures showed good agreement among JAK2 dimers and well-folded fibronectin domains. The TM domain interface was poorly defined, with small variations in distance and crossing angle between TM helices as well as PAE and pLDDT scores that urged cautious interpretation. However, all were united in that they adopted a triangular shape formed by diverging TM helices with H499 pointing in towards the interface and the JM helix forming the base of the triangle, running parallel to the membrane. We could align known structures of human Tpo-bound TpoR^2^ (8G04) and mouse JAK1-IFNGR^34^ (8EWY) back onto the resulting models with RMSD of 3.04 Å across 134 Cα atoms of the membrane-proximal hTpoR FN domain (D4) and 5.75 Å across 1326 Cα atoms of the modelled JAK2 FERM-SH2-PK dimer (this improves to 3.93 Å across 394 Cα atoms when aligning only the PK domains). In this model (**Figure 6C**), the receptor TM domains adopt a wide, right-handed crossing angle such that they interact only at the extracellular end (W491 to H499) in a conformation that moves H499 closer to the interface. This model is attractive in that it also features close contacts at W491, L494, V495 and L498 where there is clear, albeit weak, sensitivity to multiple substitutions under Tpo EC30 treatment (**Supplemental Figure 4A**), and it accounts for the lack of sensitivity to most mutations below this.

We were surprised that none of the AF3 models recapitulated the left-handed, near-parallel TpoR model selected from AF2 multimer runs in the Pogozheva et al study^26^, where Tpo-bound TpoR dimers were first modeled in the absence of JAK2. We therefore modeled TpoR ectodomain, TM domain and JM helix as a dimer with (**Supplemental Figure 6**) and without (**Supplemental Figure 7**) Tpo and inspected the resulting structures. When Tpo was present, 18 of the 20 resulting structures (4 seeds, 5 models each) again showed a similar triangular arrangement as described above. The two distinct structures instead resembled the arrangement selected in the prior study^26^. Interestingly, when we modeled the same TpoR sequence without Tpo, 11/20 structures resembled the Pogozheva et al model^26^ and only 9/20 resembled the triangular arrangement, suggesting the slight asymmetry enforced by Tpo binding favors the triangular arrangement in the AF3 calculation. To determine if the triangular TpoR TM-JM arrangement was stable, we performed relaxation with molecular dynamics in an explicit bilayer. In preparing the model (**Figure 7A**), we used a representative AF3 dimer with Tpo, encompassing TpoR 26-515, and fused this with a model of from our initial calculation of the TpoR dimer 516-571 and the JAK2 FERM-SH2-PK dimer 36-841. Because the distances separating the TpoR 515 CA atoms in the two models were almost identical, this fusion required minimal editing. The resulting model (**Figure 7B**) was placed in an explicit bilayer populated with POPC and cholesterol, surrounded by solvent and progressively relaxed with reducing harmonic restraints. When restraints were removed entirely, the system rapidly reached equilibrium with minimal deviations from the starting model (**Figure 7C-D**), supporting the plausibility of this arrangement within a membrane environment.

**Figure 7.**
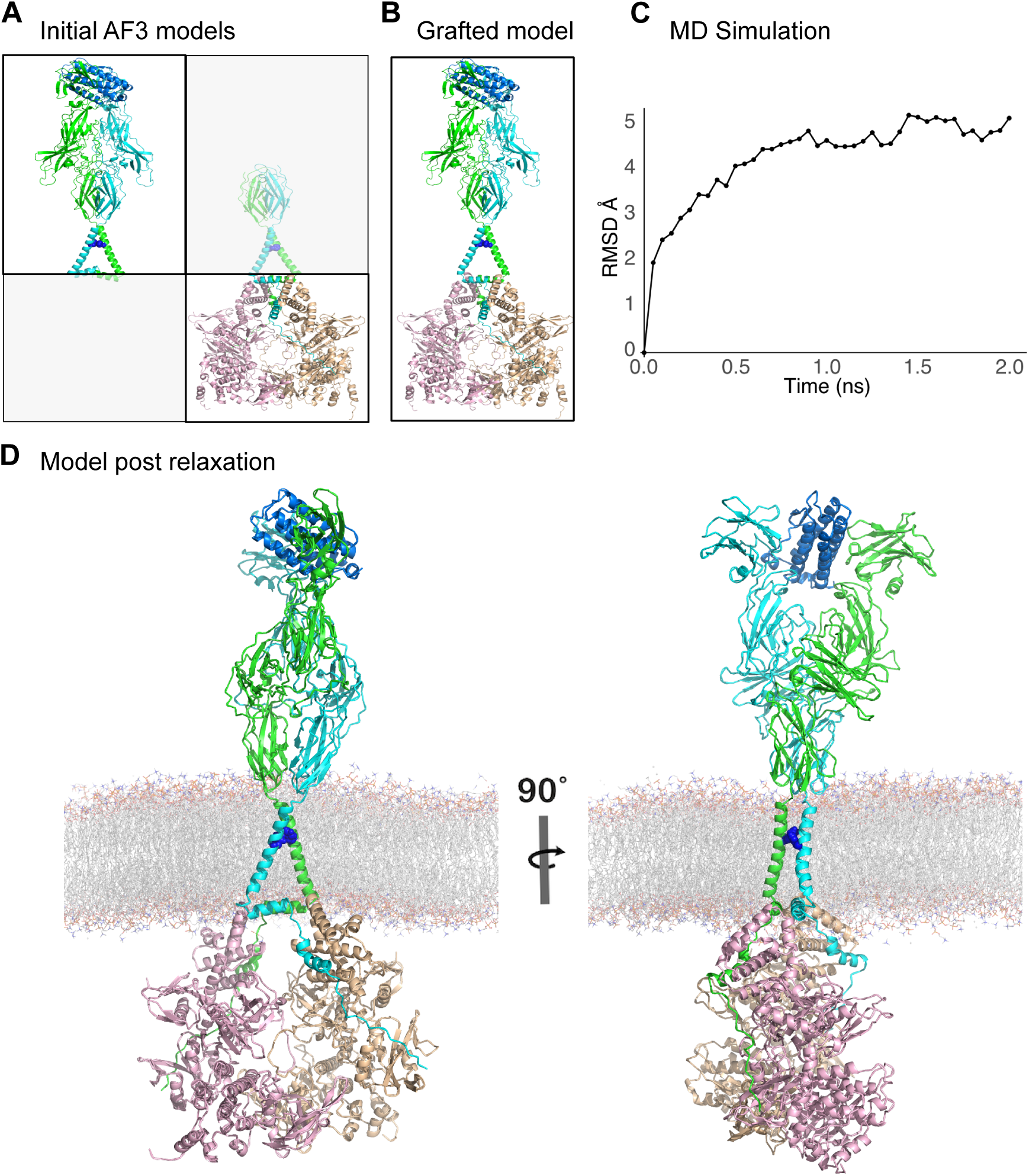
Molecular dynamics relaxation of Tpo-TpoR-JAK2 in a lipid bilayer. **(A)** Approach taken to graft two AF3 models into a near full-length model of hTpoR:hJAK2:hTpo in 2:2:1 stoichiometry. The greyed-out areas were removed from two AF3 models. **(B)** The non-greyed out areas of the models in (A) were grafted together into a single PDB file. **(C)** The model was relaxed into a model lipid bilayer. RMSD distances over the final relaxation run are plotted showing convergence. **(D)** Two views of the final model after MD relaxation.

To experimentally test whether any contacts among the TM domains are required for Tpo-mediated signaling, we made two TpoR constructs that simultaneously replace all or most TM positions with valine (**Figure 8A**): “long polyV” replaced the entire predicted TM domain between W491 and W515 with valine (leaving these tryptophans intact), while “short polyV” left the native sequence intact up to T496 and replaced A497-R514 with valine. Both expressed well in Ba/F3 cells (**Figure 8B**), but neither could be activated by the TM-binding drugs (**Supplemental Figure 8**), as expected due to the loss of H499. However, WT and short polyV receptors were similarly sensitive to Tpo (**Figure 8C**), with only a small shift in EC50 (∼2-fold higher) on the long version. In stark contrast, romi-peptide (**Figure 8D**) had significantly (∼50-fold) reduced activity on the short poly-valine construct and much lower (∼1000-fold) activity on the long version, compared to the WT receptor. The bell-shaped dose response curves for romi-peptide were much more pronounced than for Tpo, indicating that the polyV TpoRs were not only less responsive to this agonist but also more easily saturated by monomeric binding (see Discussion). We conclude from these experiments and the DMS data presented earlier that the AF3-derived model of diverging TM domains represents a plausible conformation of the Tpo-activated receptor. This is also consistent with prior studies showing that there is more than one active TM helix orientation^6,8,28^.

**Figure 8.**
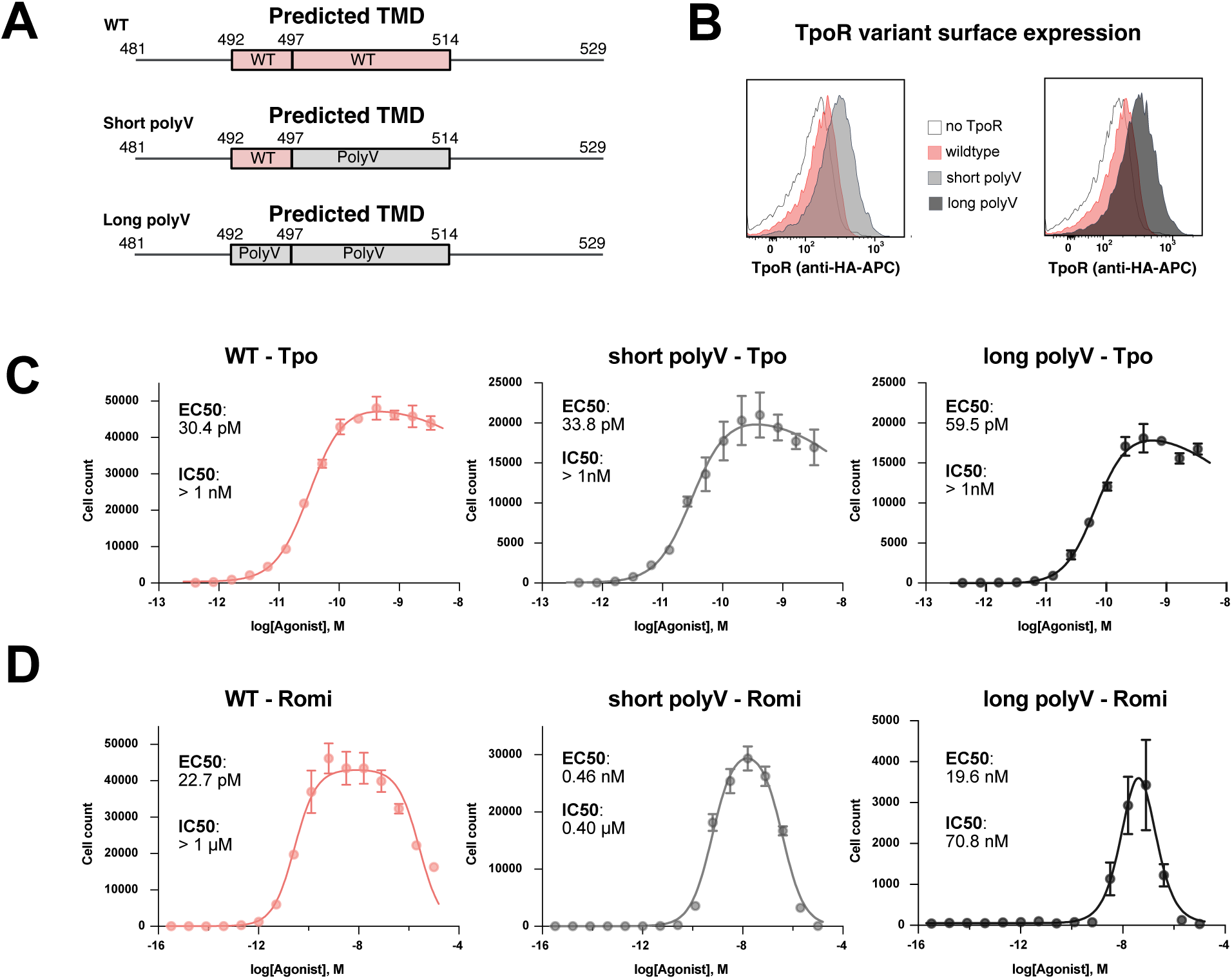
Activity of hTpoR variants with poly-valine TM replacements. **(A)** Design of the short and long poly-valine hTpoR constructs to test the AF3 model. **(B)** Surface expression of HA-tagged WT and poly-valine versions of hTpoR in Ba/F3 cells. **(C-D)** WT and poly-valine hTpoR response to Tpo (C) or romi-peptide (D) treatment. Mean ± SD of three technical replicates is shown for one representative experiment. EC50 values were calculated as the mean of two biological replicates. For biphasic response curves, EC50 denotes the midpoint of the activation phase and IC50 indicates the midpoint of the inhibition phase.

### Role of the intracellular JM amphipathic helix

Both models predict a short (three-turn) amphipathic helix from A519 to L528, similar to previous results obtained from AF2 predictions on W515K receptor fragments^28^. Our DMS data clearly show that this feature is required for stable cell surface expression, because proline substitutions in this region cause significant expression defects (see **Figure 2C**). The AF3-derived MD model also suggests an explanation for the incompatibility of polar substitutions at Y521, L524 and L528, which display perfect α-helical periodicity, with surface expression. While AlphaFold does not incorporate information about the cell membrane, in the partial receptor AF3 models (**Figure 6D-E**) and in our MD-relaxed, near-full-length model (**Figure 7D**), these three amino acids all point up into the inner leaflet of the lipid bilayer, such that they insert into the hydrophobic core and are stabilized by surrounding basic residues that could bind negative lipid headgroups. Poor surface expression of variants likely to have defective membrane association suggests that the amphipathic helix plays a role in JAK2 binding, receptor stabilization at the cell surface, or both.

Other class I cytokine receptors have similar intracellular JM sequences that are required for function, featuring two hydrophobic residues with helical spacing that are located immediately preceding the Box 1 motif^29^, and NMR studies of EpoR TM-JM monomeric peptides^35,36^ indicated helical structure extending well beyond the membrane-embedded portion. We generated an AF3 model of Epo-and JAK2-bound human EpoR (**Supplemental Figure 9, Supplemental Figure 10** and **Supplemental Discussion**) that agrees very well with other models of the TM helix dimer interface^26^ and is compatible with ligand-bound ECD and active JAK2 structures. As with hTpoR, this model predicts a break in secondary structure between the TM and JM helices (at H250/R251) and orients the amphipathic helix laterally to a hypothetical membrane such that hEpoR L254/I258 and JAK2 L224 could insert into the hydrophobic interior. We also performed a DMS screen of the hEpoR TM-JM region similar to the hTpoR one described above (see **Supplemental Figure 11-12** and **Supplemental Discussion**) and observed that hEpoR surface expression was much less sensitive to intracellular JM mutations (**Supplemental Figure 12A**). However, responses to both high (EC95; **Supplemental Figure 12B**) and low (EC30; **Supplemental Figure 12C**) concentrations of Epo were significantly impaired by mutations at L254 and I258 (surface-corrected residual plots in **Supplemental Figure 12F-G**), consistent with early studies on mouse EpoR^29^. Again, these positions could be aliphatic or (sometimes) aromatic, but not polar or small. Interestingly, cysteine at positions 253-259 actually improved surface expression (**Supplemental Figure 12A**) and maintained or enhanced sensitivity to Epo (**Supplemental Figure 12B-C**), which could reflect creation of an acylation site that reinforces membrane association. These results indicate that JM helix membrane association may be a common feature of cytokine receptor signaling that has not been accounted for in mechanistic models of activation (see Discussion).

### Identification of new constitutively active, disease-associated variants and confirmation in a clinical cohort

In a previous DMS screen focused solely on constitutive activity^9^, we found that S493L and S493C variants of hTpoR had weak constitutive activity, and many hydrophobic residues at this position enhanced S505N-driven cell growth. We also noted in the present study that hTpoR S493 variants often had elevated fitness scores, particularly in low-concentration agonist conditions (see **Supplemental Figure 2B, D, F, H, J, L**). Because our previous screen was performed as a pooled assay containing very strongly activating W515 variants that may have masked weaker constitutive activity, we rescreened a subset of positions from T481 to L511 for ligand-free activity. Indeed, we identified (**Figure 9A**) and validated (**Figure 9B-C**) variants with significant constitutive activity that were not observed in our prior studies: S493V, S493I, S493M, S493F and S493A as well as T487A, T487W and L502T. We assessed next-generation sequencing (NGS) data^37^ from patients referred for myeloproliferative neoplasm testing to Peter MacCallum Cancer Centre from Nov 2021 to Oct 2024 (**Figure 9D**) and identified three patients with an S493A (one with a type 1 *CALR* mutation and two in *cis* with *MPL* W515L), one with an S493C (with a type 1 *CALR* mutation), one with an S493F (in cis with a *MPL* W515L) and one with a T487A (with a type 1 *CALR* mutation). The T487A has also been reported by others in the context of hereditary thrombocytosis^38^ and acute megakaryoblastic leukemia^39^. The mutations S493V, S493I, S493M, T487W, L502T were not observed in our cohort. Importantly, none of these latter substitutions are possible via single-base changes. In all of the models depicted in this study, S493 points towards the membrane interior, and the hydrophobic nature of activating mutations observed in this screen suggest that stabilization of membrane insertion may explain their enhancing properties. Thus, in addition to the new insights into sequence-function signatures that define different modes of hTpoR activation, our large DMS study has identified still more activating mutations that are present in patients with myeloproliferative disease and may contribute to disease evolution or severity.

**Figure 9.**
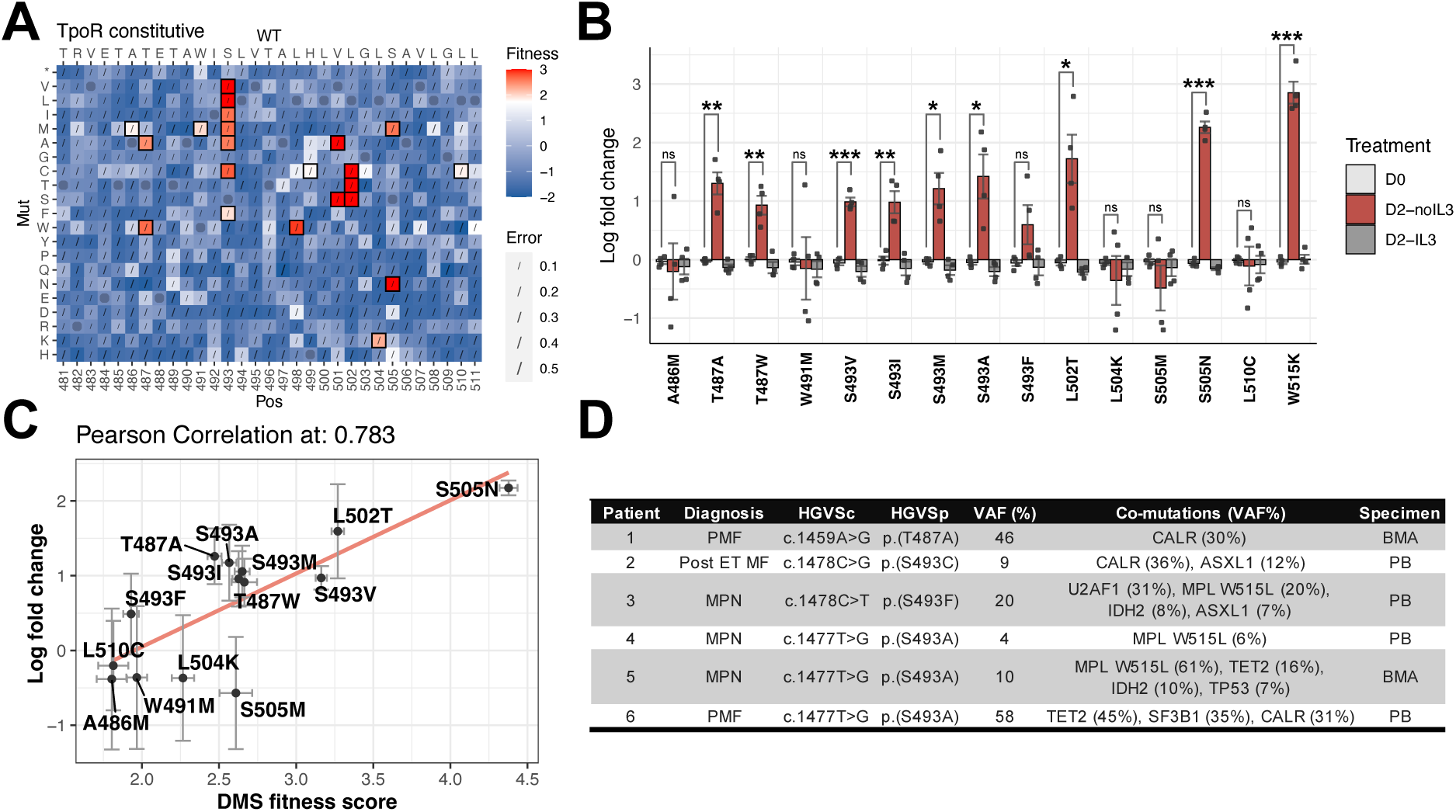
hTpoR S493 is a hotspot for constitutively active or enhancing mutations. **(A)** Heatmap of the constitutive activation DMS screen. Variants with fitness scores over the maximum fitness score of synonymous wildtype variants are shown in red and boxed; others are shown in blue. **(B)** Plot of cell ratio changes of individually tested variants relative to wildtype over four biological replicates (error bar: mean ± SEM). A paired t-test was performed to determine significance levels. p-value < 0.001, ***; 0.001 ≤ p-value < 0.01, **; 0.01 ≤ p-value < 0.05, *; p-value ≥ 0.05, ns. Ratio at day 0 is shown in grey; ratio at day 2 without IL3 treatment is shown in red; ratio at day 2 with IL3 treatment is shown in dark grey. **(C)** Correlation of DMS constitutive activity fitness scores (x-axis) and ratio changes of variants relative to wildtype without IL3 treatment on a logarithm scale (y-axis). The log ratio is the average score of four biological replicates, with vertical error bar indicating standard deviation of these four replicates. The Pearson correlation is 0.783, with the brown line representing the correlation trend. Horizontal error bar shows errors from the DMS dataset. **(D)** Diagnosis of patients. PMF: primary myelofibrosis; ET: essential thrombocythemia; Post-ET MF: post-essential thrombocythemia myelofibrosis; MPN: myeloproliferative neoplasm; PB: peripheral blood; BMA: bone marrow aspirate. The *mpl* gene is identified from MPL NM_005373.2.

## DISCUSSION

The identification of mutations that result in ligand-free hTpoR signaling^10–12,14,21^ and the discovery of small-molecule agonists^16,17,19^ have long focused attention on the TM-JM regions of hTpoR as essential signal transducers. Here we report a comprehensive sequence-function study of these regions in the context of hTpoR activation by four different classes of stimuli, where coupled systematic analysis of ligand responses and surface expression allowed us to focus our attention on TM-JM variants whose functional defects cannot be explained by altered expression, and parallel interrogation of treatment groups allowed identification of patterns not previously appreciated. For example, while S505N and W515K likely induce hTpoR dimers with similar TM rotations, their requirements for specific contacts to form the active structure are dramatically different and this is reflected in their divergent sequence-function signatures. This likely reflects the strength of the asparagine contact^31,32^ in S505N, such that very little in the bottom half of the TM helix can counteract it, and the fact that the structural changes induced by W515 mutations at the membrane-cytosol transition^28,40^ largely circumvent the need for structural reinforcement in the top half.

Synthetic ligands showed very strong signatures of sequence specificity and significant shared requirements in and around the TM domains. Dimension reduction analysis revealed a TM-dependent signature that is shared by all three small-molecule drugs and, to a large extent, a peptide mimic of the biologic agonist romiplostim. This similarity to romi-peptide surprised us because we expected this agonist to phenocopy Tpo^1,20^, but aside from a requirement for H499 that is unique to the small-molecule drugs, the sequence dependencies of romi-peptide are largely indistinguishable from those of small-molecule agonists. Our experiments with poly-valine TpoR constructs confirmed that TM domain interactions cooperate to support the response to romi-peptide, showing a large loss of sensitivity on the short polyV and larger yet for the long polyV. We suggest that in contrast to the asymmetrical binding of Tpo, which enforces a very specific ECD dimer geometry^1,2^, the symmetrical binding mode of romi-peptide effectively brings two TpoR molecules together but relies heavily on TM-JM interfaces to enforce the active conformation and transmit it through the membrane to JAK2. We interpret the progressively narrowing bell-shaped dose response curves on polyV receptors as indicating that, in the absence of this cooperative conformational coupling, the receptor is very easily saturated with non-productive monomeric binding in a 1:1 stoichiometry.

The shared aspects of the synthetic agonist signature are all consistent with a single TM interface similar to the AlphaFold Multimer model from Pogozheva et al^26^ and with a large body of mutagenesis and engineering studies on TpoR, including the locations of most known constitutively activating mutations^9^. This signature includes, among other features, a very specific requirement for L508 and a sensitivity to small substitutions at L498 and hydrophobic substitutions at R514. The specificity at L508 is striking, and it implicates this position in specific structural contacts consistent with the cysteine crosslinks referenced above^7^. Our experiments with L508V and L508I variants revealed that not even similar branched aliphatic amino acids can support full activity, reflecting a degree of complementary packing that must be close enough to preclude β-branched side chains. We^9^ and others^23^ have previously identified L498 as a site where constitutive activity can arise uniquely via mutation to tryptophan, and the particular sensitivity to small amino acid substitutions here is consistent with a need for complementary packing. The role of R514 is likely in stabilizing the TM-JM transition at the intracellular membrane limit, as is common in membrane proteins^41^, because it points directly away from the AlphaFold Multimer model interface.

Perhaps the most striking feature of the shared synthetic agonist signature is the strong dependence on hydrophobicity at positions W491-T496 for activation, despite polar substitutions in this region being very well tolerated for surface expression. We believe this reflects two closely related aspects of hTpoR biology: while there is strong evidence for the specific helical interface at W491, L494, V495 and L498 in the non-native active conformation, the more general negative effect of loss of hydrophobicity at surrounding positions on activation, but not surface expression, also indicates that stable insertion in the membrane is crucial for the active but not the inactive structure. Our data support a model in which the inactive dimeric or monomeric TpoR has this region outside the membrane^23^ and possibly non-helical, and eltrombopag or related drugs help it insert to form the active TM interface. Our finding that S493 variants with hydrophobic substitutions are both weakly activating and enhancing (this study and reference^9^) under many treatment conditions, despite the location of S493 outside of the dimer interface in all models, suggests that their functional effect is also to encourage membrane insertion. Their identification in patients only (thus far) with additional oncogenic lesions would also seem to indicate that they are pathological primarily as second-site mutations. Altogether, these data are consistent with a structural transition in the extracellular JM-TM region proposed by others^18,23^, and they imply that a shift in register of the whole TM domain with respect to the membrane could play a role in transmitting structural changes to the intracellular end by inducing concomitant structural changes at the RWQFP motif^21,28^.

The observation that even drastic changes to sequences in and near the membrane-embedded portion of the receptor have only small effects on signaling from the native ligand Tpo indicates that its binding mode very effectively enforces an active state with little reliance on cooperative interactions through the JM and TM regions. Although weak, most detrimental effects on Tpo signaling were associated with changes at the extracellular JM-TM boundary discussed above. Our AF3 modeling identified a series of closely related alternative active structures to the one identified for synthetic ligands and activating mutations, all of which had a similar triangular arrangement of the TM domains, with the JM helix running parallel to the membrane. This global conformation, which may be formed from an ensemble of similar arrangements rather than a single unique active structure, is consistent with the lack of mutational effects to Tpo-mediated receptor triggering in the rest of the TM domain. In this conformation, W491-H499 still mediates dimeric TM interactions, but the TM helices diverge below this such that there are no further contacts until the intracellular amphipathic helices come back together at the intracellular membrane surface. Experiments with poly-valine receptors further supported this model, demonstrating that the entire bottom half can be simultaneously substituted with valine and only the additional replacement of I492-T496 causes appreciable activation defects.

Our DMS results and structural modeling in the intracellular JM region suggest a previously unappreciated role of an amphipathic helix at Y521-L528, just beyond the well-studied RWQFP motif, in mediating membrane binding. The amphipathic helical nature of this sequence is particularly important for hTpoR surface expression, making assessment of a further functional role difficult. However, two other recent studies point to membrane association of hTpoR W529^28^ and of the JAK2 FERM domain^42^ in the activated state, lending further weight to its potential functional significance, and we note that our AF3 models place JAK2 L224 in an ideal position to insert into the membrane as reported in Wilmes et al^42^, aided by electrostatic interactions between a large basic patch and negatively charged inner membrane lipids. Structural studies of EpoR and leptin receptor (LEPR) tails bound to JAK2 FERM-SH2 dimers^43^ suggest that the last aliphatic residue in the corresponding sequences (part of the “switch” region) could bridge JAK2 molecules by directly binding the FERM domain, which they do not do in the models shown here. However, since these crystal structures were determined in the absence of TM domains and lipids, it is difficult to determine whether this reflects a functionally relevant interaction or merely an alternative way to satisfy the need to bury these residues in a hydrophobic environment during crystallization. Our DMS screen and AF3 modeling of human EpoR further supports membrane association of an intracellular JM amphipathic helix as a general feature of active-state class I cytokine receptors. While early studies of mouse EpoR proposed that this helical segment is a rigid extension of the TM domain^29^, and solution NMR structural studies have modeled it as such^35,36^, we note that the continuity of backbone amide nuclear Overhauser effects defining helical structure in both NMR studies was interrupted precisely at the point where our AF3 model predicts a break (H250/R251 in hEpoR). The flexibility that this implies could therefore be consistent with lateral membrane binding of this helix as pictured here. Ultimately, the precise function of this region and its potential structural role in positioning JAK dimers for transactivation at the inner membrane surface will require experimentally determined receptor complex structures in lipid bilayers.

In summary, our systematic mutagenesis of the hTpoR membrane-associated domains, with parallel analysis of surface expression and many different stimulating conditions, provides a uniquely comprehensive view of sequence-function relationships that confirms and unites many prior studies and provides novel insights into the relative contributions of key structural features to hTpoR function. Our central finding, that Tpo binding induces an active TM conformation that is distinct from the structures induced by synthetic agonists and activating mutations, raises the enticing possibility that pathological signaling could be specifically inhibited using membrane-targeted strategies that would leave normal physiological Tpo signaling intact.

## METHODS

### Plasmids

pDONR221-HA-hTpoR was a gift from Andrew Brooks and used as a cloning intermediate before receptor sequences were installed into pMX-Gateway vectors. The expression vectors pMX-GW-PGK-Puro-GFP and pMX-GW-PGK-Puro-mCherry^9^ were used to express hTpoR and hEpoR variants in Ba/F3 cells. The sequences of TpoR wildtype, S505N and W515K mutants, EpoR wildtype, and associated oligos used in this study are available in the Supplementary Material. Individual TpoR mutations were introduced via a two-step overlap extension PCR. The poly-Valine constructs were created by HiFi assembly of oligos (IDT) with the PCR-amplified pDONR221-HA-hTpoR backbone (see Supplemental Materials below).

### Cell lines

Verified HEK293T cells were sourced from Cellbank Australia (#12022001) and maintained in DMEM (GIBCO, #10313039) supplemented with L-glutamine and 10% FBS (GIBCO, #2526728RP). WEHI-3 cells^44^ were cultured in RPMI-1640 (GIBCO, #11875093) supplemented with L-glutamine (GIBCO, #25030081) and 10% FBS. When WEHI-3 cells reached confluence, the medium was harvested and sterile-filtered as a source of interleukin 3 (IL3). IL3-dependent murine pro-B Ba/F3 cells^45^ were maintained in RPMI-1640 (GIBCO, #11875093) supplemented with L-glutamine, 10% FBS and 10% IL3 medium, harvested from WEHI-3 cells. To make the counting cell line, we transduced pLKO.1-TRC (a gift from Timothy Ryan; Addgene plasmid #191566) into Ba/F3 cells and sorted mTagBFP2 positive cells twice. The cells were maintained in 2 ng/μl puromycin (Thermo, #A1113803) for one week to eliminate non-transduced cells, then aliquoted in cryoprotective medium containing 90% FBS + 10% DMSO and stored in liquid nitrogen. Ba/F3 cells overexpressing murine JAK2 were prepared by transducing Ba/F3 cells with pMSCV-IRES-mCherry-mJAK2 (a gift from Jeff Babon). Cells with high mCherry expression were sorted twice to establish a cell line with JAK2 over-expression. The cell line was aliquoted in cryoprotective medium containing 90% FBS + 10% DMSO and stored in liquid nitrogen. A fresh batch of cells was thawed for new experiments.

### DMS library construction

We mutated every residue in the JM and TM domains using overlap extension PCR^9,46^ (**Supplemental Figure 1)**. Briefly, an oligonucleotide containing a degenerate codon (NNN) was paired with an invariant oligonucleotide to amplify the 5’ fragment of the TpoR open reading frame. The 3’ fragment was amplified using invariant primers and overlapped with the mutated 5’ fragment. These two fragments were used in another round of PCR to generate the full TpoR open reading frame flanked by attL sites, allowing direct insertion of the fragment into pMX-GW-PGK-Puro-GFP using LR clonase (Invitrogen, #11791020). Each position was transformed into electro-competent DH5α (NEB, #C2989), ensuring at least 200 colonies were obtained for each variant codon. These colonies were harvested and miniprepped (Qiagen, #27104). The same steps were performed on every position of interest, and colonies were pooled to generate different libraries: L1 encompasses mutations at residues 488 to 516; L2, residues 481 to 511; L3, residues 507 to 529; and L4, residues 481 to 529. S505N and W515K DMS libraries were made via assembling pooled degenerate oligonucleotides into PCR-amplified destination vectors via HiFi-assembly (NEB, #E2621L), and use S505N and W515K as base sequences. EpoR DMS library was synthesized by Twist Bioscience and initially cloned into the pDONR vector. We then transferred the genes into the expression vector pMX-GW-PGK-Puro-GFP using LR clonase. EpoR variants were also pooled into different libraries: L1 encompasses residues 215 to 243; L2, residues 238 to 266; L3, residues 215 to 266.

### Retroviral and lentiviral production and transduction

To make retrovirus, HEK293T cells were transfected with a transfer vector (DMS libraries or individual TpoR variants) and retroviral packaging vectors MMLV-gag-pol and Env MLVgp2 (gifts from Marco Herold) using calcium phosphate. Parental or JAK2 overexpressing Ba/F3 cells were transduced in the presence of 4 μg/ml polybrene and 10% WEHI-3 medium, at 1,000g, 30°C for 1 hour. Media was replaced the following day, and the transduction efficiency was checked 2 days post-transduction. Cells were subjected to 2 ng/μl puromycin for 2 days to eliminate non-transduced cells.

### Surface staining with anti-HA antibody

Cells were stained with biotinylated anti-HA (Roche, #12158167001) at 250 ng/ml diluted in FACS buffer (PBS (Thermo Fisher, #BP6651) + 100 μg/ml BSA (Sigma-Aldrich, #A7030) + 2 mM EDTA (Invitrogen, #15575020)) on ice for 30 min. Cells were washed 3 times with FACS buffer before staining with streptavidin-allophycocyanin (Biolegend, #405207) diluted 1:800 in FACS buffer. Cells were analyzed using flow cytometry.

### Agonist dose response assay

We purchased the following agonists: Eltrombopag (MCE, #HY-15306), Avatrombopag (MCE, #HY-13463), Lusutrombopag (MCE, #HY-19883), Thrombopoietin (BioLegend, #763702), and Erythropoietin (PeproTech, #100-64). The Romi-peptide was a gift from Jeff Babon and Nadia Kershaw (WEHI)^1^. Agonists were either diluted in DMSO (Sigma-Aldrich, #D4540) or in the case of Tpo, RPMI to create 3-fold dilution series. 1.5 μl of diluted agonists were added to wells in a 96-well U-bottom plate (Corning, #3799) in triplicate. We also added 1.5 μl of either DMSO or RPMI as a no-treatment control. Ba/F3 cells stably expressing TpoR (or EpoR) were harvested at around 80% confluence and washed with 1x DPBS (GIBCO, #14190144) three times to remove residual IL-3. The cells were resuspended in RPMI, and 150 μl of the cell suspension, containing around 10,000 cells per well, were seeded into wells containing agonists. Cells were incubated at 37°C, 5% CO2 for 2 days. Cells were pelleted and resuspended in 100 μl RPMI containing 10,000 mTagBFP2 positive Ba/F3 counting cells. We performed the flow cytometry on FACSymphony™ A3 Cell Analyzer (BD Biosciences). Live cell counts for each well were determined by gating live single cells and determining the percentage of BFP+ counting cells compared to the percentage of GFP+ hTpoR-transduced Ba/F3 cells in the total population:

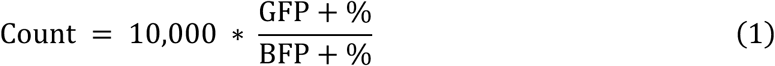

The cell count data was used for dose-response curve fitting in Prism10, with drug concentration transformed into logarithmic scale, to calculate the EC50:

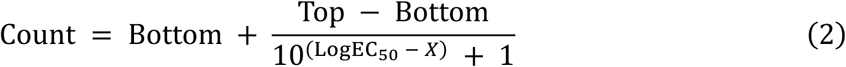

Bell-shaped dose response curves observed with high dose Romi-peptide treatments were fit with the “nls” function from Rstudio v4.3.1 using the following formula:

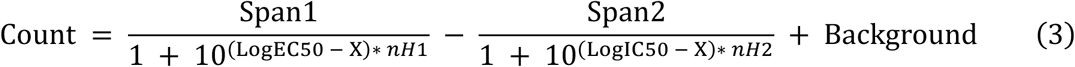

Where Background was fixed using the average cell count from the 3 lowest concentrations of romi-peptide. Span1 and Span2 are mandated to be the same, while nH1 and nH2 are fixed at 1. When fitting saturated TpoR response to Tpo, we only fixed background at the average of the 3 lowest concentrations of Tpo and only report EC50.

*Competition assay:*

For dose response assays in main **Figure 5**, 10,000 Ba/F3 cells expressing wildtype hTpoR (with IRES-mCherry) were mixed with 10,000 Ba/F3 cells expressing hTpoR variants (with IRES-GFP) and cultured in the presence of TpoR agonists, IL3 or no agonist for 2 days. Cell counting was performed using mTagBFP2 positive counting cells by flow cytometry.

For constitutive activity measurements in **Figure 9B**, Ba/F3 cells expressing wildtype hTpoR (with IRES-mCherry) were mixed with Ba/F3 cells expressing hTpoR variants (with IRES-GFP) and the cell ratio measured immediately on day 0. 10,000 mixed Ba/F3 cells were added to triplicate wells +/-IL3 and incubated for 2 days. Cells were harvested and the ratio of GFP to mCherry positive cells was measured again.

### Agonist and Constitutive Activity DMS

We transduced our receptor libraries into 1.2 million BaF3 cells, with a multiplicity of infection of 0.1 to favor single integration events, aiming for at least 40 cells per variant. 2 days after transduction, we selected cells in 2 ng/μl Puromycin for two days. Cells were subsequently harvested and washed three times with PBS and selected. The cells were treated with agonists at high (EC95) and low (EC30) concentrations, maintained in IL-3, or maintained without growth factors (constitutive activity) for 2 days. Dead cells were removed after 2 days with Ficoll-PaqueTM PLUS (Cytiva, #GE17-1440-02) by layering cultures onto 0.5 ml of Ficoll and spinning at 800g for 25 minutes at room temperature. The supernatant containing viable cells was decanted into a fresh tube, leaving dead cells behind in the pellet and mixing the Ficoll such that viable cells could then be pelleted and washed with PBS. RNA was extracted using a RNeasy Mini Kit (Qiagen, #74104).

### Receptor Surface Level DMS

To identify surface presentation-deficient variants, we used BaF3 cells overexpressing JAK2 as the recipients of our receptor libraries and transduced 1.2 million cells at a multiplicity of infection of 0.1. Two days after transduction, we selected cells in 2 ng/μl puromycin for 2 days. Cells were then stained with anti-HA-biotin (3F10) and SA-APC and sorted into surface high and low populations based on APC mean fluorescence intensity, with the top 50% of cells allocated to “surface high” and the bottom 50% to “surface low”. After sorting, cells were directly lysed and their total RNA was extracted using the RNeasy Mini Kit (Qiagen, #74104) following the manufacturer’s instructions.

For all selections, we ensured at least two independent transductions were performed per dataset (**Supplemental Table 1**).

### Next-Generation Sequencing sample preparation

RNA was reverse-transcribed using the LunaScript® RT SuperMix Kit (NEB, #E3025L) and a primer annealing 3’ to the region of interest and featuring a 5’ overhang containing a 16 base pair unique molecular identifier (UMI) and an invariant sequence for 3’ index primer annealing (see supplementary material). The reverse-transcribed product was then amplified by a forward primer that bound 5’ to the region of interest and contained an invariant sequence for 5’ index primer annealing and a reverse primer that bound to the 3’ index overhang. The resulting PCR product was indexed in triplicate using the flanking overhangs to allow multiplexed sequencing. The indexed samples were sent for NextSeq 1000 or NextSeq 2000 (Illumina), paired-end sequencing at the WEHI Advanced Genomic Facility. The average reads per barcode for each sample were typically around 100.

### Illumina data processing and DMS scoring

We demultiplexed and trimmed sequence to the region of interest with cutadapt v3.4^47^. Reads were deduplicated based on their UMI to remove PCR artefacts by moving 16bp UMIs from the 3’ end of the amplicon with UMI-tools^48^ to the address line in the fastq file. The amplicons were then aligned to the reference gene (wild type sequence) with bowtie2^49^, sorted and indexed with SAMtools^50^. The indexed bam file was deduplicated using the cluster method of UMI-tools with an edit-distance-threshold at 1, before being returned to the fastq file format with SAMtools.

The deduplicated fastq files were used for fitness score calculation with DiMSum v1.3^51^. At least 5 input reads in any replicate were required for each variant to be retained (except for RomiLow dataset whose minimum input read was set at 0). Read quality scores over 15 in all replicates were also required. DiMSum calculates fitness scores by comparing variant frequency before (input) and after (output) selection. For agonist treatments, IL3 treated cells were used as input; for constitutive activity screens, the plasmid library itself was used as input and cells surviving after growth factors were removed were specified as output; for surface level screens, the low gate was input and the high gate was output.

In some experiments, selection was too strong, such that we encountered noisy data from near-complete loss of some variants. To account for missing variants without changing the distribution pattern of variants post-selection, we randomly sampled a subset of reads from input samples equal to 10% of the total reads from the output sample, using seqkit^52^. These reads were combined with the output sample so that variants that were at low frequency in both input and output samples no longer returned volatile scores. This strategy was applied to AvaLow and wildtype TpoR constitutive datasets.

### Normalization of DiMSum Fitness scores

DiMSum outputs scores where the wildtype variant is normalized to 0. Except for the data in **Figure 9A**, we re-normalized data to set the mean of nonsense variants to 0 and the mean of synonymous wildtype variants to 1. After normalization scores from libraries L1, L2, L3 and L4 (see supplementary table 1 for which libraries contributed to each dataset) were combined so that continuous sequence-function maps could be plotted.

### Residual analysis

We graphed surface expression fitness to activity fitness scores to determine the correlation between expression and activity scores, and determined which variants were outliers based on this correlation. Scatter plots showed two major populations: synonymous-like variants centered at (1,1); and nonsense-like variants centered at (0,0). We drew a straight line between the centers of both populations and calculated the distance of every variant to the line. Errors for this score were determined by propagating surface expression and activity errors 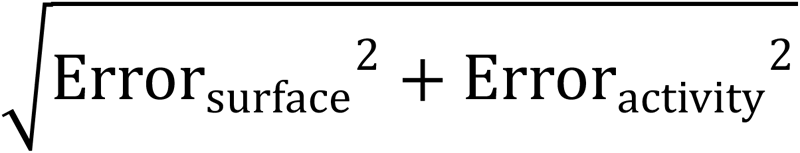. We masked variants that were close to the line by considering the standard deviation of synonymous wildtype and nonsense variant distance scores using the following criteria:

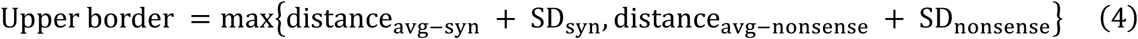

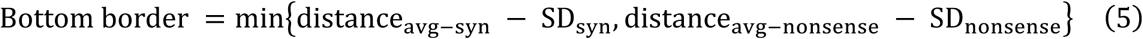

Where distance_@A$(BCD_and distance_@A$(D#DBEDBE_represent mean distance of the synonymous and nonsense variants. SD_BCD_and SD_D#DBEDBE_ represents the standard deviation of the distance of the synonymous and nonsense variants.

Variants outside of the border were colored in gold or green gradients based on their distance above or below the line using “ggplot2” in R studio v4.3.3.

### Clustering TpoR variants

We performed uniform manifold approximation and projection analysis (UMAP)^25^, using “umap” function in Rstudio v4.3.3; and coupled with the Density-Based Spatial Clustering of Applications with Noise algorithm (DBSCAN), using the “dbscan” function, to cluster variants while handling the noisy nature of DMS data. Normalized data from the following datasets were clustered: hTpoR surface, TpoHigh, TpoLow, AvaHigh, AvaLow, EltHigh, EltLow, LusuHigh, LusuLow, RomiHigh and RomiLow. We down-weighted scores from low agonist treatments by dividing their fitness scores by 3, to account for the lower signal to noise at EC30. After the dimension reduction, we performed DBSCAN clustering. The optimization of parameters was guided by the aggregation of synonymous wildtype and nonsense variants. The random seed was set at 123, and we finalized the parameters as follow: n_neighbors at 10; min_dist at 0.25 for UMAP dimension reduction; minPts at 10 and eps at 0.5 for DBSCAN clustering. The pipeline distributed most of the variants into five clusters, with a few outliers, likely representing noise, that were not clustered into any of the five clusters.

### AlphaFold3 modeling

We modelled hTpoR, hEpoR and hJAK2 with the AlphaFold server powered by AlphaFold3 (https://alphafoldserver.com/)^33^. The following modeling jobs (detailed below) were submitted: 1) Two copies of hTpoR (Uniprot: P40238) spanning residues 389 – 569 (numbering includes signal sequence, protein sequence can be found in Supplementary Materials) covering the final fibronectin domain, TM domain, Box 1 and Box 2 motifs, with two copies of hJAK2 (Uniprot: O60674) from residues 36 - 841 spanning the FERM, SH2 and PK domains. 2) Two copies of the hTpoR extracellular domain, TM domain and juxtamembrane helix spanning residues 26 – 530 with one copy of Tpo, residues 22 - 174 of the immature protein sequence. 3) Two copies of the hTpoR extracellular domain, TM domain and juxtamembrane helix spanning residues 26 – 530 in the absence of Tpo. 4) Two copies each of hEpoR (Uniprot: P19235) and hJAK2 with one copy of hEpo (Uniprot: P01588). hEpoR residues from 7 – 307 (32-332 of full length EpoR) and residues 28 – 193 (numbering includes signal sequence) of hEpo were used in the modeling.

Modeling was performed four times with independent seeds. Five models were output from each seed resulting in 20 structures.

Job 1 (Seeds: 119010, 130167, 412147, 1751667495) (**Supplemental Figure 5**)

TpoR FN-TM-JM-Box1-Box2 (residues 389:569) 2 copies

VRLPTPNLHWREISSGHLELEWQHPSSWAAQETCYQLRYTGEGHQDWKVLEPPLGARGGTLELRPRSRYRLQLRARLNGPTYQGPWSSWSDPTRVETATETAWISLVTALHLVLGLS AVLGLLLLRWQFPAHYRRLRHALWPSLPDLHRVLGQYLRDTAALSPPKATVSDTCEEV EPSLLE

JAK2 FERM-SH2-PK (residues 36:841) 2 copies

DPVLQVYLYHSLGKSEADYLTFPSGEYVAEEICIAASKACGITPVYHNMFALMSETERIWYPPNHVFHIDESTRHNVLYRIRFYFPRWYCSGSNRAYRHGISRGAEAPLLDDFVMSYLFAQWRHDFVHGWIKVPVTHETQEECLGMAVLDMMRIAKENDQTPLAIYNSISYKTFLPKCIRAKIQDYHILTRKRIRYRFRRFIQQFSQCKATARNLKLKYLINLETLQSAFYTEKFEVKEPGSGPSGEEIFATIIITGNGGIQWSRGKHKESETLTEQDLQLYCDFPNIIDVSIKQANQEGSNESRVVTIHKQDGKNLEIELSSLREALSFVSLIDGYYRLTADAHHYLCKEVAPPAVLENIQSNCHGPISMDFAISKLKKAGNQTGLYVLRCSPKDFNKYFLTFAVERENVIEYKHCLITKNENEEYNLSGTKKNFSSLKDLLNCYQMETVRSDNIIFQFTKCCPPKPKDKSNLLVFRTNGVSDVPTSPTLQRPTHMNQMVFHKIRNEDLIFNESLGQGTFTKIFKGVRREVGDYGQLHETEVLLKVLDKAHRNYSESFFEAASMMSKLSHKHLVLNYGVCVCGDENILVQEFVKFGSLDTYLKKNKNCINILWKLEVAKQLAWAMHFLEENTLIHGNVCAKNILLIREEDRKTGNPPFIKLSDPGISITVLPKDILQERIPWVPPECIENPKNLNLATDKWSFGTTLWEICSGGDKPLSALDSQRKLQFYEDRHQLPAPKWAELANLINNCMDYEPDFRPSFRAIIRDLNSLFTPDYELLTENDMLPNMRIGALGFSGAFEDRDP

Job 2 (Seeds: 1061974485, 1880406353, 1977975823, 1648165899) (**Supplemental Figure 6**)

TpoR EC-TM-JM (residues 26:530) 2 copies

QDVSLLASDSEPLKCFSRTFEDLTCFWDEEEAAPSGTYQLLYAYPREKPRACPLSSQSMPHFGTRYVCQFPDQEEVRLFFPLHLWVKNVFLNQTRTQRVLFVDSVGLPAPPSIIKAMGGSQPGELQISWEEPAPEISDFLRYELRYGPRDPKNSTGPTVIQLIATETCCPALQRPHSASALDQSPCAQPTMPWQDGPKQTSPSREASALTAEGGSCLISGLQPGNSYWLQLRSEPDGISLGGSWGSWSLPVTVDLPGDAVALGLQCFTLDLKNVTCQWQQQDHASSQGFFYHSRARCCPRDRYPIWENCEEEEKTNPGLQTPQFSRCHFKSRNDSIIHILVEVTTAPGTVHSYLGSPFWIHQAVRLPTPNLHWREISSGHLELEWQHPSSWAAQETCYQLRYTGEGHQDWKVLEPPLARGGTLELRPRSRYRLQLRARLNGPTYQGPWSSWSDPTRVETATETAWISLVTALHLVLGLSAVLGLLLLRWQFPAHYRRLRHALWP

Tpo (residues 22-174) 1 copy

SPAPPACDLRVLSKLLRDSHVLHSRLSQCPEVHPLPTPVLLPAVDFSLGEWKTQMEETKAQDILGAVTLLLEGVMAARGQLGPTCLSSLLGQLSGQVRLLLGALQSLLGTQLPPQGRTTAHKDPNAIFLSFQHLLRGKVRFLMLVGGSTLCVR

Job 3 (Seeds: 189583173, 40579582, 531222035, 1400440743) (**Supplemental Figure 7**)

TpoR EC-TM-JM (residues 26:530) 2 copies

QDVSLLASDSEPLKCFSRTFEDLTCFWDEEEAAPSGTYQLLYAYPREKPRACPLSSQSMPHFGTRYVCQFPDQEEVRLFFPLHLWVKNVFLNQTRTQRVLFVDSVGLPAPPSIIKAMGGSQPGELQISWEEPAPEISDFLRYELRYGPRDPKNSTGPTVIQLIATETCCPALQRPHSASALDQSPCAQPTMPWQDGPKQTSPSREASALTAEGGSCLISGLQPGNSYWLQLRSEPDGISLGGSWGSWSLPVTVDLPGDAVALGLQCFTLDLKNVTCQWQQQDHASSQGFFYHSRARCCPRDRYPIWENCEEEEKTNPGLQTPQFSRCHFKSRNDSIIHILVEVTTAPGTVHSYLGSPFWIHQAVRLPTPNLHWREISSGHLELEWQHPSSWAAQETCYQLRYTGEGHQDWKVLEPPLGARGGTLELRPRSRYRLQLRARLNGPTYQGPWSSWSDPTRVETATETAWISLVTALHLVLGLSAVLGLLLLRWQFPAHYRRLRHALWP

Job 4 (Seeds: 1483843195, 732948, 372395, 993239) (**Supplemental Figure 9**)

EpoR EC-TM-JM-Box1-Box2 (residues 7:307) 2 copies

DPKFESKAALLAARGPEELLCFTERLEDLVCFWEEAASAGVGPGNYSFSYQLEDEPWKLCRLHQAPTARGAVRFWCSLPTADTSSFVPLELRVTAASGAPRYHRVIHINEVVLLDAPVGLVARLADESGHVVLRWLPPPETPMTSHIRYEVDVSAGNGAGSVQRVEILEGRTECVLSLRGRTRYTFAVRARMAEPSFGGFWSAWSEPVSLLTPSDLDPLILTLSLILVVILVLLTVLALLSHRRALKQKIWPGIPSPESEFEGLFTTHKGNFQLWLYQNDGCLWWSPCTPFTEDPP

ASLE JAK2 FERM-SH2-PK (residues 36:841) 2 copies

DPVLQVYLYHSLGKSEADYLTFPSGEYVAEEICIAASKACGITPVYHNMFALMSETERIWYPPNHVFHIDESTRHNVLYRIRFYFPRWYCSGSNRAYRHGISRGAEAPLLDDFVMSYLFAQWRHDFVHGWIKVPVTHETQEECLGMAVLDMMRIAKENDQTPLAIYNSISYKTFLPKCIRAKIQDYHILTRKRIRYRFRRFIQQFSQCKATARNLKLKYLINLETLQSAFYTEKFEVKEPGSGPSGEEIFATIIITGNGGIQWSRGKHKESETLTEQDLQLYCDFPNIIDVSIKQANQEGSNESRVVTIHKQDGKNLEIELSSLREALSFVSLIDGYYRLTADAHHYLCKEVAPPAVLENIQSNCHGPISMDFAISKLKKAGNQTGLYVLRCSPKDFNKYFLTFAVERENVIEYKHCLITKNENEEYNLSGTKKNFSSLKDLLNCYQMETVRSDNIIFQFTKCCPPKPKDKSNLLVFRTNGVSDVPTSPTLQRPTHMNQMVFHKIRNEDLIFNESLGQGTFTKIFKGVRREVGDYGQLHETEVLLKVLDKAHRNYSESFFEAASMMSKLSHKHLVLNYGVCVCGDENILVQEFVKFGSLDTYLKKNKNCINILWKLEVAKQLAWAMHFLEENTLIHGNVCAKNILLIREEDRKTGNPPFIKLSDPGISITVLPKDILQERIPWVPPECIENPKNLNLATDKWSFGTTLWEICSGGDKPLSALDSQRKLQFYEDRHQLPAPKWAELANLINNCMDYEPDFRPSFRAIIRDLNSLFTPDYELLTENDMLPNMRIGALGFSGAFEDRDP

Epo (residues 28:193) 1 copy

APPRLICDSRVLERYLLEAKEAENITTGCAEHCSLNENITVPDTKVNFYAWKRMEVGQQAVEVWQGLALLSEAVLRGQALLVNSSQPWEPLQLHVDKAVSGLRSLTTLLRALGAQKEAISPPDAASAAPLRTITADTFRKLFRVYSNFLRGKLKLYTGEACRTGDR

### Molecular Dynamics

To perform molecular dynamics simulations (**Figure 7**), a PDB file was created by aligning the lowest ranking models from Jobs 1 & 2 in Pymol and pair-fitting the W515 CA atoms of both TpoR chains. The distance between Job 1 W515 CA to that in Job 2 was 0.8 Å. TpoR residues 26-515 with Tpo from Job 2 and TpoR residues 516-569 with JAK2 from Job 1 were selected and output to a new PDB file. The PDB was edited to ensure consistent numbering of residues in each chain, and Coot was used to first alter the rotamer of R514 to remove a steric clash with Y521 introduced at the splice site. Subsequently, the Coot tool “Regularize Zone” and Phenix “Geometry Minimization” were used to correct minor issues around the join site.

Molecular dynamics simulations of the membrane-embedded TpoR were conducted using the GROMACS 2024 molecular dynamics package, employing the GROMOS 54a7 force field^53^ and using heavy-hydrogen atoms^54^. The AF3-derived model was incorporated into a lipid bilayer composed of 80% POPC and 20% cholesterol, which was generated using the MemGen webserver^55^. The system was solvated with simple point charge (SPC) water and 150 mM NaCl, with counterions added to maintain charge neutrality. All simulations were performed under periodic boundary conditions. The system underwent energy minimization using the steepest descent algorithm, followed by equilibrations with progressively decreasing harmonic restraints on the protein. This was achieved through four consecutive 2 ns simulations with a 2 fs timestep, applying harmonic restraints of 1000, 500, 100, and 10 kJ/mol/nm², respectively. A final equilibration without harmonic restraints (0 kJ/mol/nm²) was conducted to equilibrate the protein over a 2 ns simulation with a 2 fs timestep. Random initial velocities were assigned at the beginning of each equilibration, based on a Boltzmann distribution. Temperature was maintained at 310 K using the Bussi-Donadio Parrinello velocity rescale thermostat with a coupling constant of 0.1 ps^56^. Pressure was regulated at 1 bar using semi-isotropic pressure coupling with the Berendsen barostat (*https://doi.org/10.1063/1.448118), using an isothermal compressibility of 4.5 x 10^-5^ bar^-1^. The SETTLE algorithm (https://doi.org/10.1002/jcc.540130805) was utilized to constrain the geometry of water molecules, while the LINCS algorithm^57^ was used to constrain the covalent bond lengths of the solute. Electrostatic interactions were computed using the Particle Mesh Ewald method, and non-covalent interactions were evaluated using the Verlet scheme with a 1.4 nm cut-off. Simulations were visualized and RMSD analysis was performed using VMD v1.9.4^58^.

## ACKNOWLEDGEMENTS

This work was supported by NHMRC Project Grant 1157348 to MJC and MEC and funding from the Wilson Centre for Blood Cancer Genomics. AFR was supported by the National Institutes of Health grants UM1HG011969 and R01HG013025. The DMS libraries were created with the help of the WEHI Multiplexed Assay Technology Hub (MATH), which was founded with WEHI New Medicines and Advanced Technology Theme funding. We thank Stephen Wilcox, Sarah MacRaild, WEHI Flow Cytometry and Advanced Genomics Facilities for technical resources. Work in the laboratories of the authors was made possible through Victorian State Government Operational Infrastructure Support (OIS) and Australian Government NHMRC Independent Research Institute Infrastructure Support (IRIIS) Scheme. This project received grant funding from the Australian Government.

## AUTHORSHIP

MJC and MEC conceived and supervised the project. The manuscript was prepared by XW, MJC and MEC, and edited by all co-authors. HM, AM, SR, MJC and XW performed DMS data collection. MG, AM, and XW processed samples for NGS sequencing. XW processed DMS data, under the supervision of MJC and MEC and with advice from AFR. XW performed cellular experiments on individual variants, with the assistance from JVN. WACB performed molecular dynamics simulations. SH and PB provided clinical sequencing data. PB serves on the Advisory Boards of Adaptive Biotechnologies, Roche Diagnostics Corporation, and AbbVie and Speakers’ Bureaus: AstraZeneca, Janssen Cilag, and the European Medicines Evaluation Agency.

