## Supplementary Information for "Shared and distinct sequence-function signatures define different modes of human TpoR activation"

### Supplemental Figures

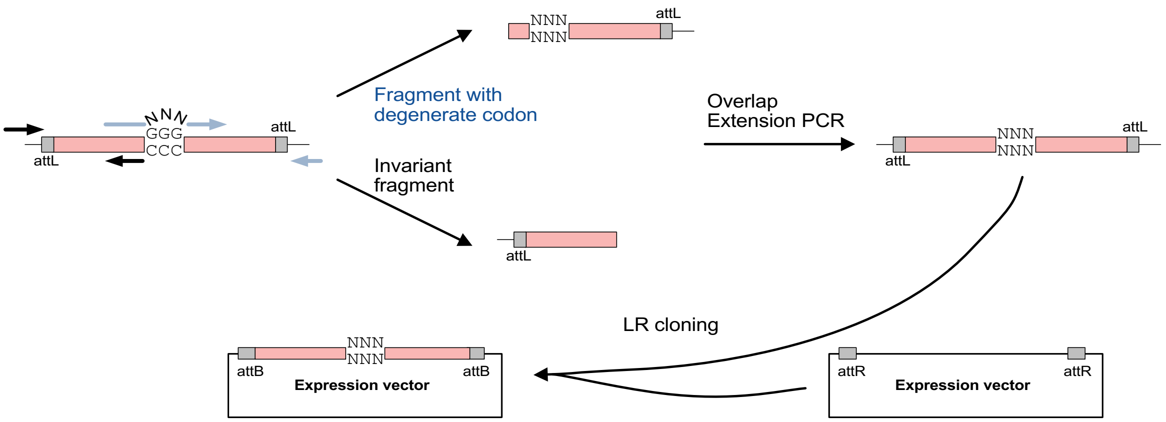


**Supplemental Figure 1.** Overlap extension PCR for library construction. An invariant 5' fragment is combined with a 3' fragment containing degenerate codons for overlap extension PCR to generate the complete open reading frame. The resulting amplicon is then inserted to expression vectors by LR cloning.

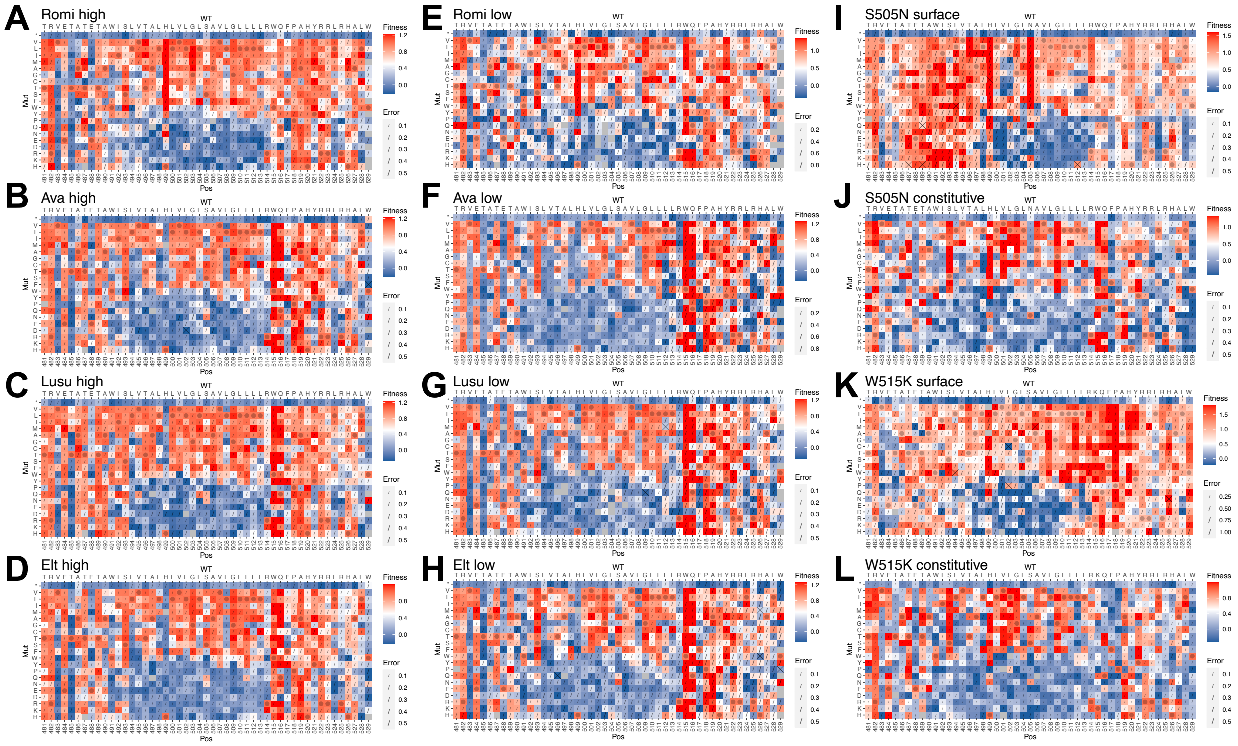


**Supplemental Figure 2. Heatmaps of all synthetic agonist and constitutively active mutant DMS datasets.** (A-L) Sequence-function heat maps are presented as in main **Figure 2** and labeled at the top left with the selection condition. All screens are based on growth/survival over two days under the indicated treatment, with the exception of panels (I) and (K), in which the selection was flow-sorting into high and low surface TpoR populations.

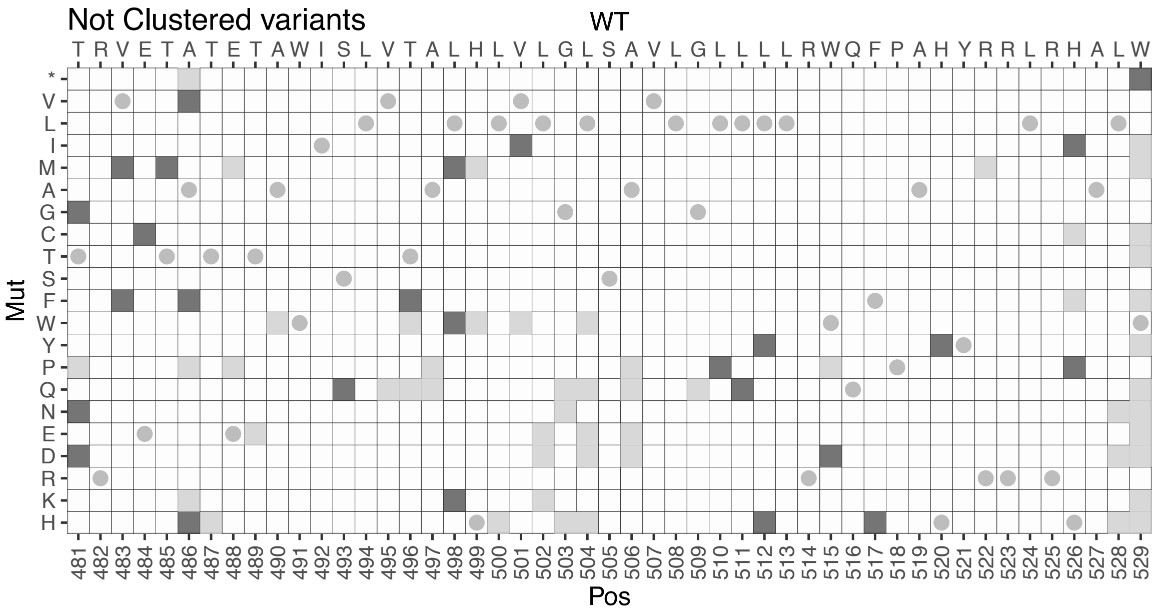


**Supplemental Figure 3.** Variants not clustered in main **Figure 3A** indicated by dark grey squares. Dots indicate the WT positions and light grey squares indicate variants that were not included in UMAP clustering because data were missing in one or more conditions.

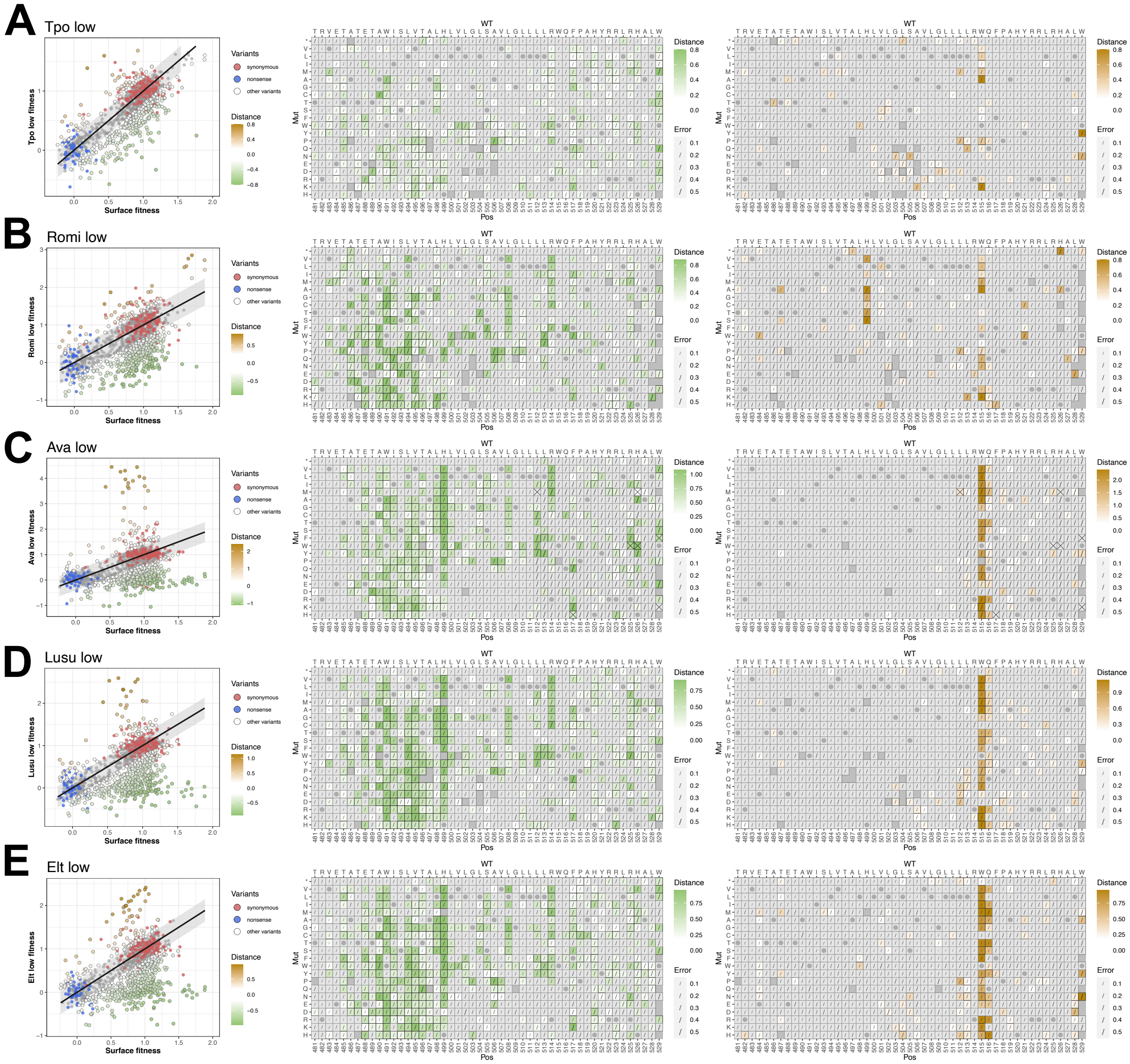


**Supplemental Figure 4. Residual analysis of low-dose agonist treatment datasets.** The residual analysis compares response fitness scores (y-axis) with the surface fitness scores (x-axis). (**A-E**) The shaded ribbon on the scatter plot was calculated from the spread of synonymous wildtype and nonsense variants, encompassing 2 SD. Outlier variants below the ribbon are displayed in a gradient of white to green, scaled by their distance from the straight line, and are functionally deficient. Variants above are highlighted in gold indicating the degree of enhanced activity. Scores for defective (middle panel) or enhanced (right panel) variants are separately mapped back onto the heat map structure for visual comparison to one another. Errors indicated by slashes were calculated as the square root of the sum of squared surface expression and activity errors, and variant scores with an error over 0.5 are marked with a cross.

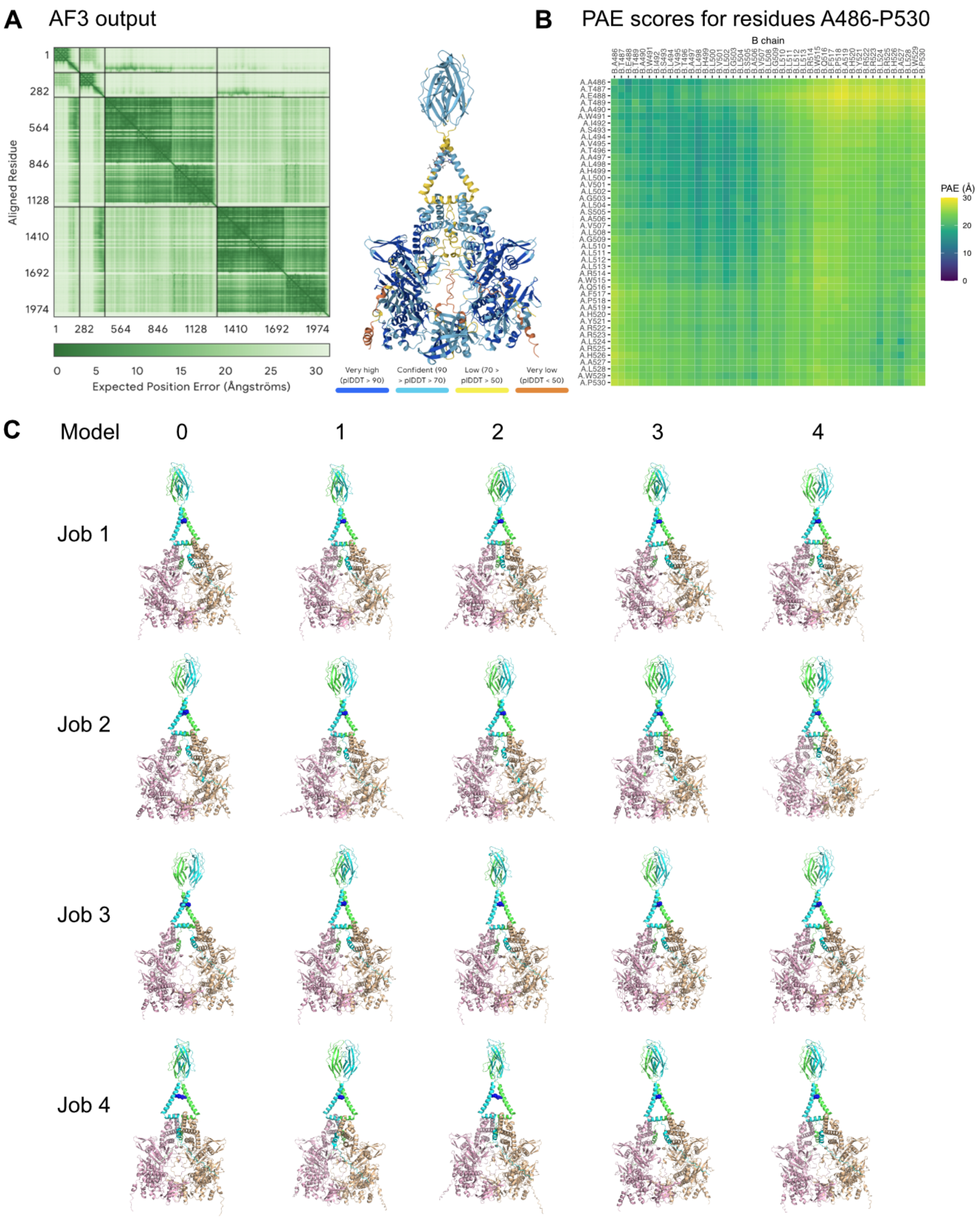


**Supplemental Figure 5. Summary of AF3 runs for 2x hTpoR FN-TM-JM-Box1-Box2, 2x hJAK2 FERM-SH2-PK**. A) Output from AF3 of representative structure showing whole model PAE matrix and ribbon diagram colored by pLDDT scores. B) PAE scores relating to the TM-JM domain residues 486-530 of hTpoR. C) All models output from each of the 4 AF3 Jobs. hTpoR chains are in green and cyan, hJAK2 is in wheat and light pink. The position of H499 of hTpoR is shown as blue spheres in each model.

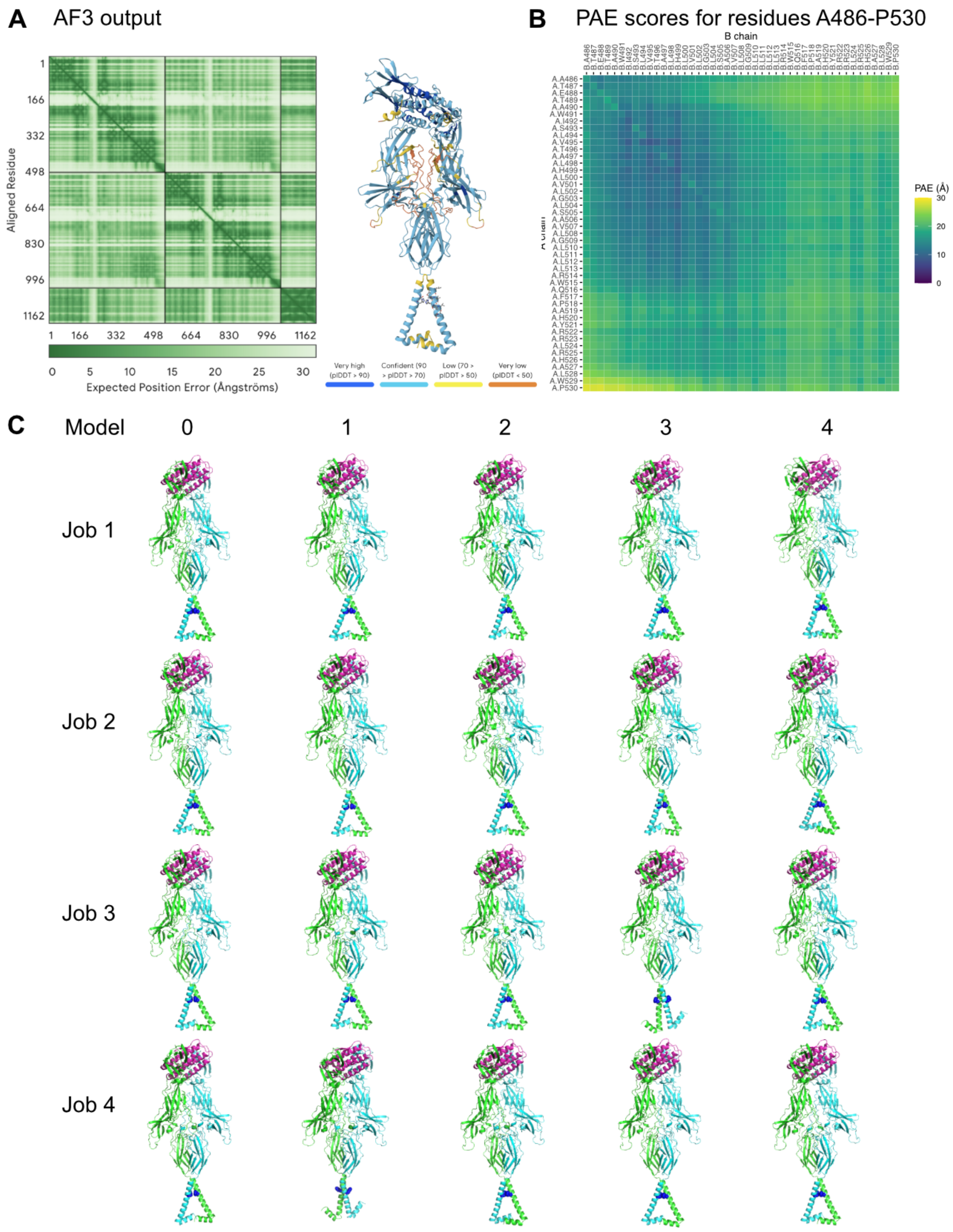


**Supplemental Figure 6. Summary of AF3 runs for 2x hTpoR EC-TM-JM, 1x hTpo**. A) Output from AF3 of representative structure showing whole model PAE matrix and ribbon diagram colored by pLDDT scores. B) PAE scores relating to the TM-JM domain residues 486-530 of hTpoR. C) All models output from each of the 4 AF3 Jobs. hTpoR chains are in green and cyan, hTpo is in magenta. The position of H499 of hTpoR is shown as blue spheres in each model.

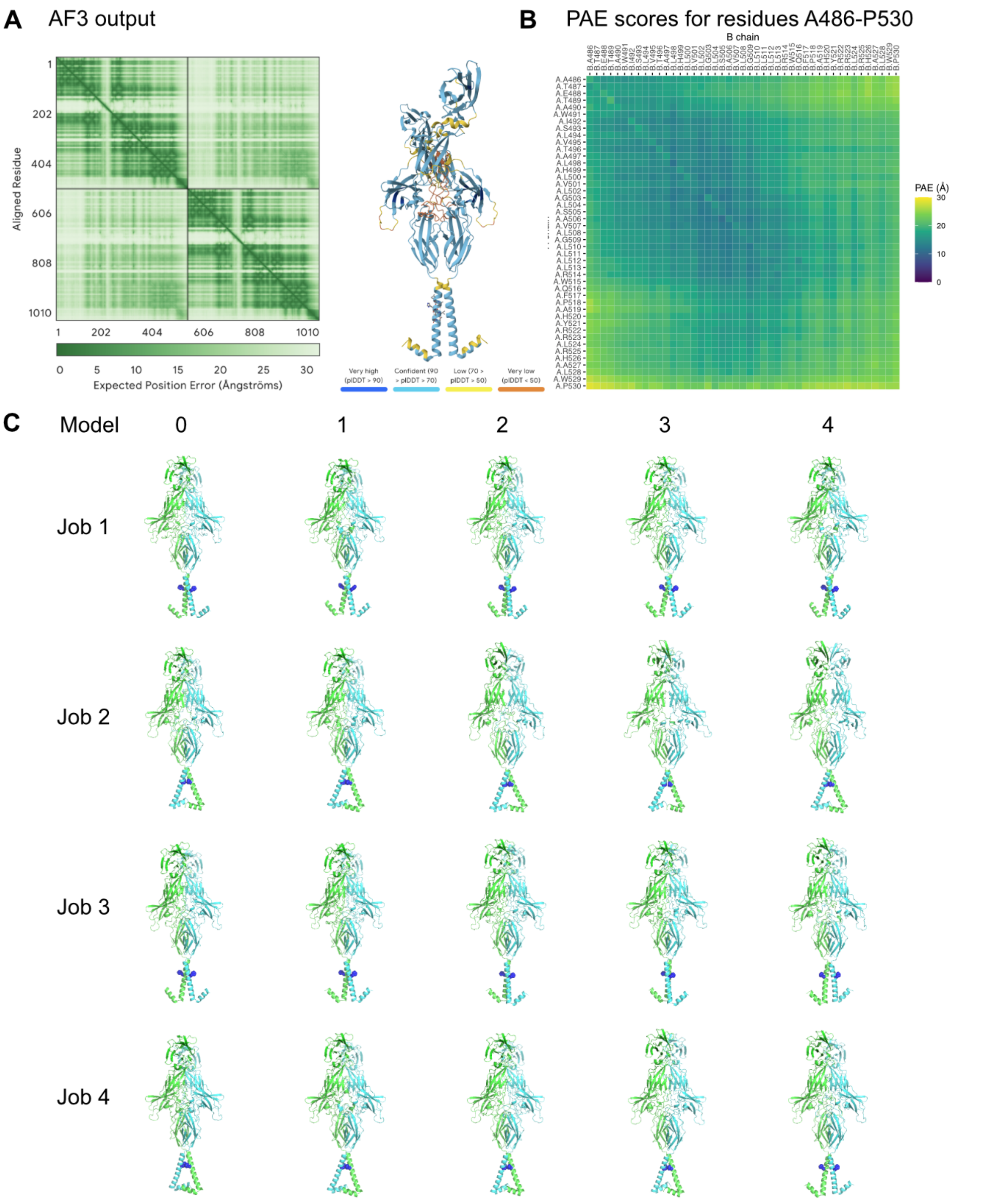


**Supplemental Figure 7. Summary of AF3 runs for 2x hTpoR EC-TM-JM**. A) Output from AF3 of representative structure showing whole model PAE matrix and ribbon diagram colored by pLDDT scores. B) PAE scores relating to the TM-JM domain residues 486-530 of hTpoR. C) All models output from each of the 4 AF3 Jobs. hTpoR chains are in green and cyan. The position of H499 of hTpoR is shown as blue spheres in each model.

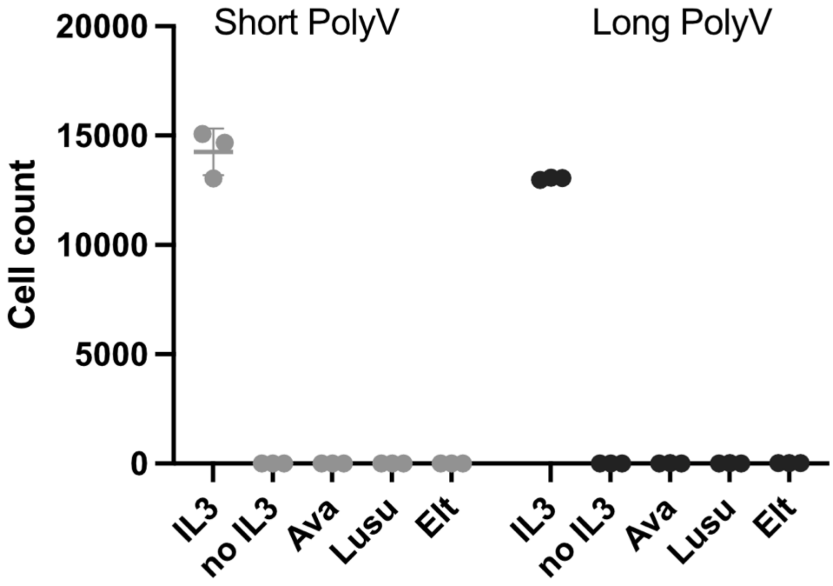


**Supplemental Figure 8. TpoR poly-valine transmembrane variants do not respond to avatrombopag (Ava), lusutrombopag (Lusu) or eltrombopag (Elt).** Labels on the x-axis indicate treatment group, Ava, Lusu and Elt were tested at EC95. Cell count is depicted on the y-axis. Short poly-valine is shown as light gray, while long poly-valine in dark grey. Representative experiment is shown with technical triplicates and error bars indicating mean ± SD. Three biological replicates were performed with similar results.

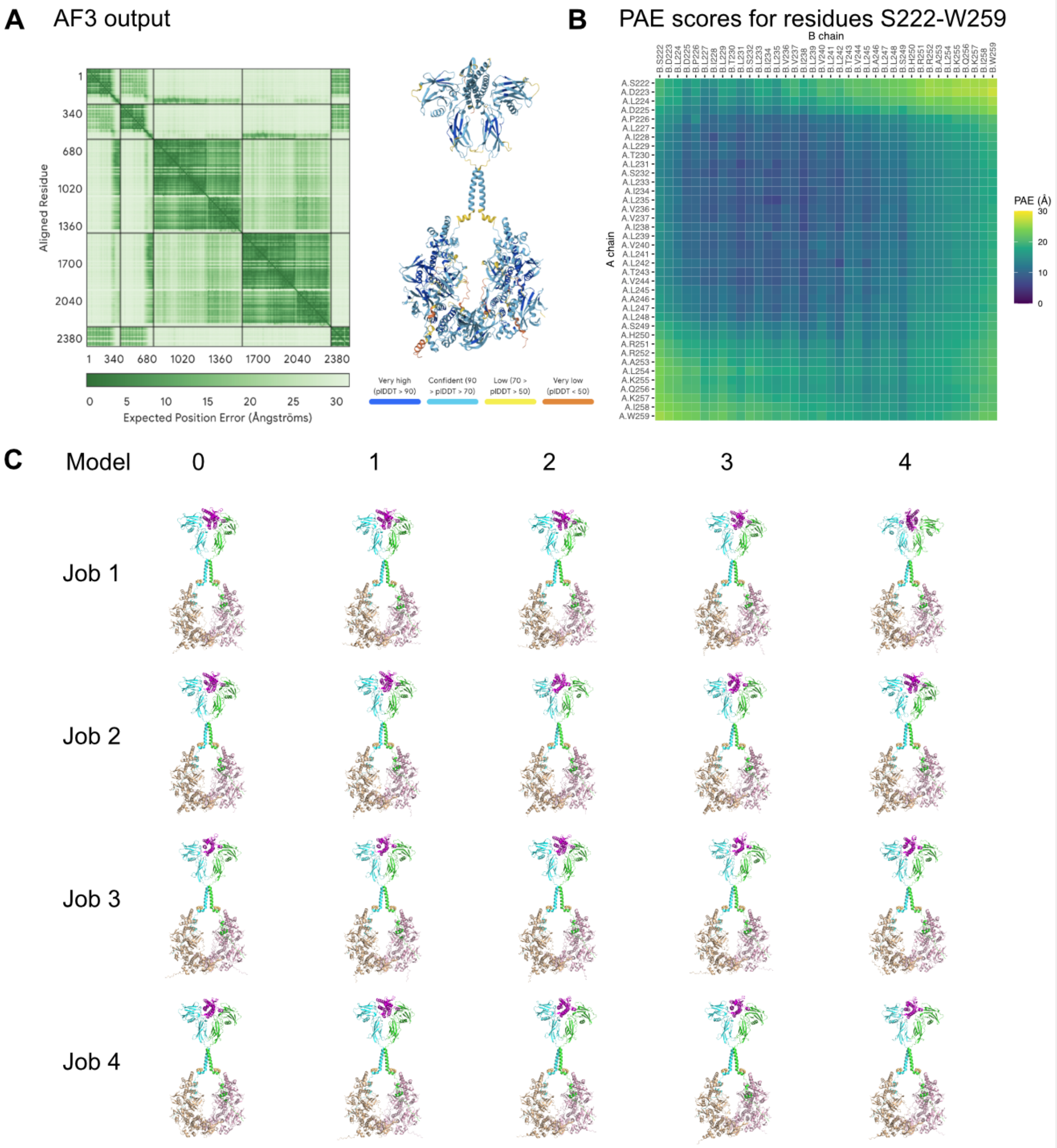


**Supplemental Figure 9. Summary of AF3 runs for 2x hEpoR, 2x hJAK2 FERM-SH2-PK, 1x hEpo**. A) Output from AF3 of representative structure showing whole model PAE matrix and ribbon diagram colored by pLDDT scores. B) PAE scores relating to the TM-JM domain residues 222-259 of hEpoR. C) All models output from each of the 4 AF3 Jobs. hEpoR chains are in green and cyan, hEpo is in faded red, hJAK2 is in light pink and wheat. The position of hydrophobic residues (L247, I251) in the JM of hEpoR are shown as orange spheres in each model.

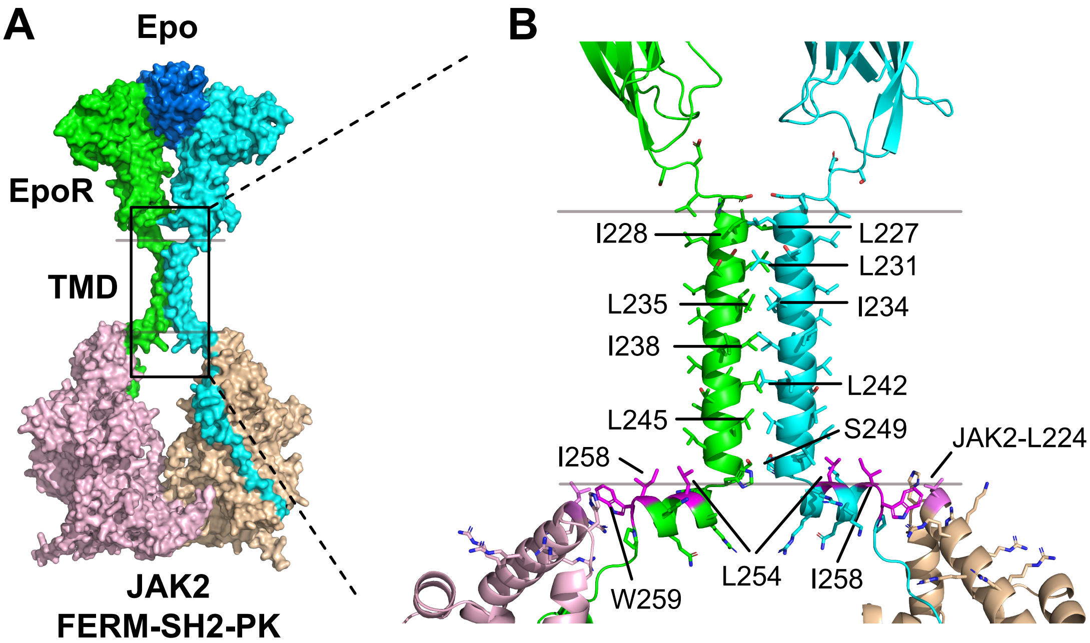


**Supplemental Figure 10. AlphaFold 3 model of activated hEpo-hEpoR-JAK2 shows membrane-bounds JM amphipathic helices similar to hTpoR. (A)** AlphaFold3-generated model of human Epo-EpoR-JAK2 (FERM-SH2-PK). Gray lines indicate the approximate boundaries of the cell membrane. Boxed region is enlarged in **(B)** to show TM dimer interface and intracellular JM helix in position to bind the inner leaflet of the lipid bilayer. TM interface residues are labelled to the left and right. Hydrophobic residues in the intracellular JM amphipathic helix are highlighted in magenta. JAK2 L224 is highlighted in violet.

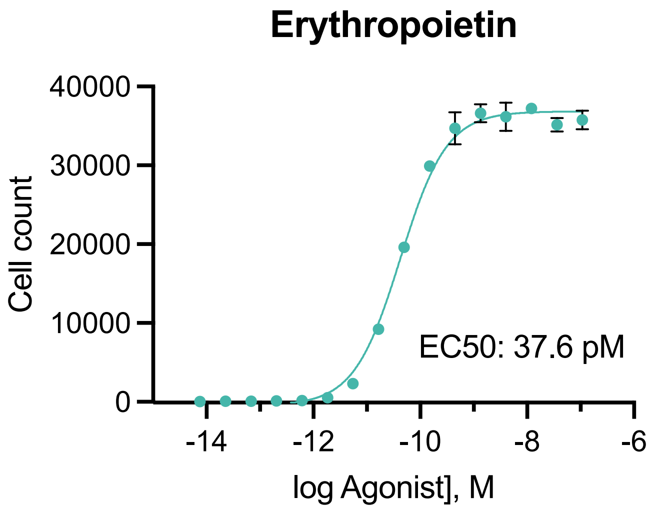


**Supplemental Figure 11. Human Epo dose response on WT hEpoR-expressing Ba/F3 cells.** The EC50 was calculated as the mean of two independent replicates.

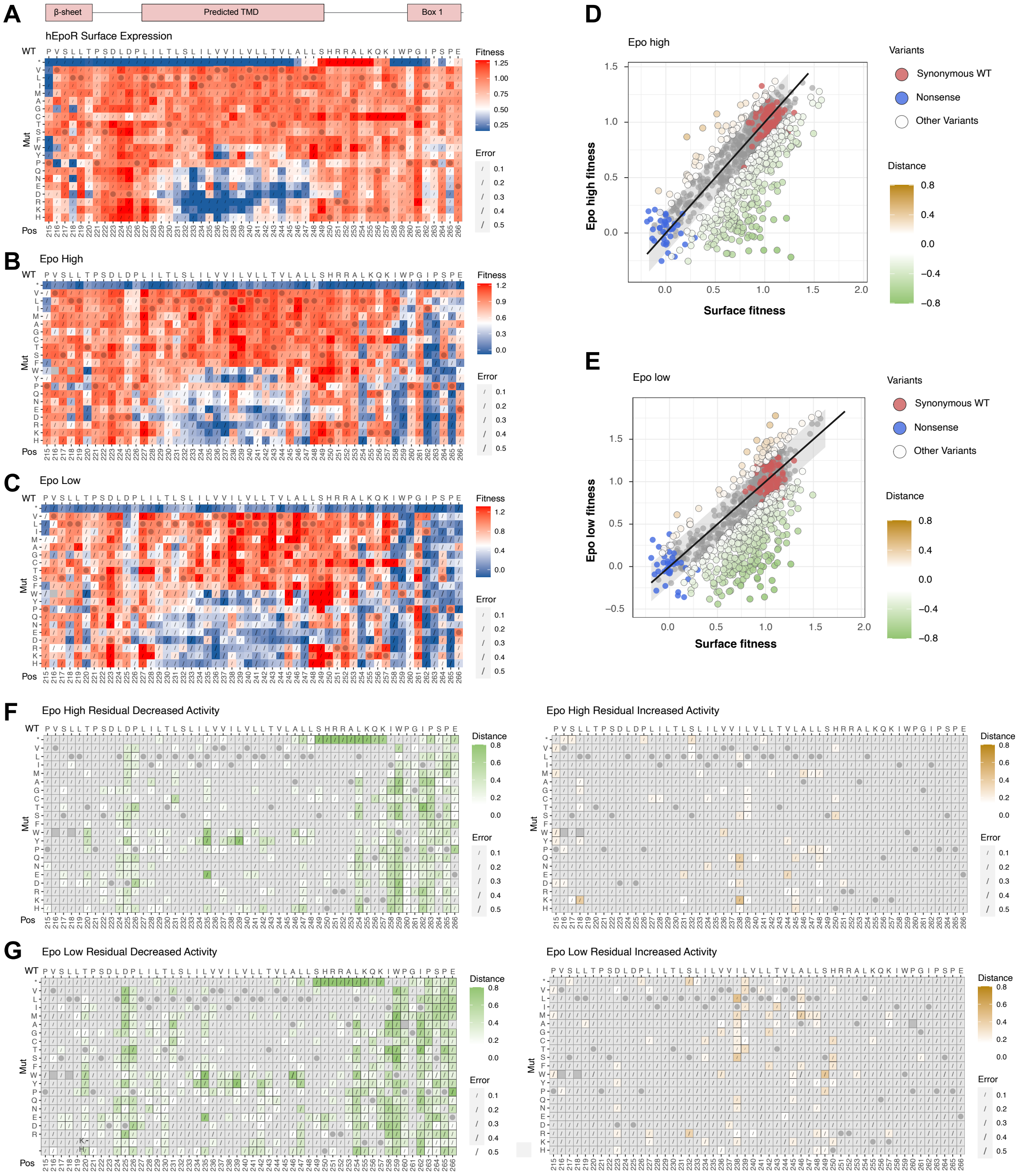


**Supplemental Figure 12. DMS and residual analysis of hEpoR surface expression and Epo response in Ba/F3 cells.** DMS sequence-function heatmaps showing **(A)** hEpoR variant surface expression and response to Epo at high concentration (EC95; **B**) and low concentration (EC30; **C**). Predicted secondary structure is illustrated above the heatmaps. Columns show consecutive amino acid residues in the hEpoR protein sequence from N-terminal (left) to C-terminal (right) direction for the region scanned in this study, numbered from the first residue of the mature sequence (without signal peptide). Native sequence is indicated at the top and position number at the bottom. Rows indicate the substituting amino acids (or premature stop codon *) and each box depicts the normalized fitness score in a three-color gradient: blue indicates impaired EpoR expression or Epo response, red indicates WT-like expression or Epo response, white indicates intermediate expression or Epo response. WT amino acid at each position is indicated with a dot and error (derived from "Sigma" calculated by DiMSum) is indicated by slashes with the length proportional to magnitude. Missing variants are shown in grey. **(D-E)** Scatter plots of Epo response fitness scores (y-axes) at high (EC95, panel D) and low (EC30, panel E) concentrations against fitness scores reflecting surface expression levels (x-axes), with each variant in the library represented by a dot. A straight line connects the clusters corresponding to synonymous-WT (red; centered at 1,1) and nonsense variants (blue; centered at 0,0), while most other variants are shown in gray. Variants whose activity is higher or lower than expected for their surface expression levels are indicated in shades of gold and green, respectively, and plotted in residual heat maps (**F-G**).

**Supplemental Results and Discussion: AlphaFold3 model of activated hEpoR and alignment with DMS data**

Our AlphaFold3 model of human EpoR (**Supplemental Figure 10A**) contains a TM helix dimer interface (**Supplemental Figure 10B**) consistent with other recent models of mouse and human EpoR[1] and with cysteine cross-linking studies performed on the mouse receptor[2]. The Epo-bound ectodomain dimer in our model aligns well with the corresponding crystal structure 1EER[3] (backbone RMSD of 2.21 Å over 386 Cα atoms) and with the corresponding region of the mouse JAK1-IFNGR dimer cryoEM structure 8EWY[4] (backbone RMSD of 5.92 Å over 1272 Cα atoms; this reduces to 3.501 Å over 500 Cα atoms when aligning only the PK domains). These high-quality alignments demonstrate that our model containing a potentially membrane-bound JM amphipathic helix is compatible with an active hEpoR structure.

Our hEpoR DMS screen encompassed a region beginning at P215 in the membrane proximal D2 domain and extending through the stalk, TM and intracellular tail, ending at E266 just beyond the Box 1 motif. Numbering follows the convention from prior EpoR literature of beginning from the first position after the 24-residue signal peptide. As expected, surface expression was abrogated by premature termination up to the last two hydrophobic TM residues L247-L248 (**Supplemental Figure 12A**, top row). However, unlike hTpoR, hEpoR variants that terminated after the TM domain and before box 1 (corresponding to premature stop codons at positions 249-257 inclusive) were very well expressed, reflecting an ability to traffic to the cell surface without JAK2 association. Strong polar substitutions were poorly tolerated in a core hydrophobic region of the TM domain at positions 231-244 (**Supplemental Figure 12A-C**). Various substitutions causing defects in Epo response that were not accounted for by reduced expression (**Supplemental Figure 12F-G**, left panels) were detected at the extracellular stalk D225-P226 and at TM dimer interface positions L231, I234, L235, I238, L239 and L242. In the cytoplasmic JM region, most substitutions at L254, I258, W259 and throughout the box 1 motif exhibited defective Epo responses, as did premature terminations that were nonetheless well expressed.

Few variants registered as enhancers of Epo responsiveness, but consistent signals at I238 (**Supplemental Figure 12F-G**, right panels) suggested that reinforcing polar contacts (Epo low) or potentially improved aliphatic packing (Epo high) at this central interface position could stabilize the active state. Individual variants have not been tested as traditional single-amino-acid mutants for constitutive or enhanced activity. As highlighted in the main text, cysteine variants at positions 253-259 registered increased expression (**Supplemental Figure 12A**) with corresponding increases in Epo response (**Supplemental Figure 12B-C**). This aligns precisely with the helical region positioned to bind the membrane in our AlphaFold3 model, and we speculate that this reflects the creation of S-acylation sites that could further stabilize the membrane-bound conformation of the amphipathic helix. This hypothesis has not been experimentally tested.

| Dataset | L1 | L2 | L3 | L4 |
|---|---|---|---|---|
| Tpo High | | 3 | 3 | |
| Tpo Low | | 3 | 3 | |
| Tpo Surface | | 3 | 3 | |
| Romi High | | | | 3 |
| Romi Low | | | | 3 |
| Ava High | | 3 | 3 | 6 |
| Ava Low | | 3 | 3 | |
| Lusu High | | 3 | 3 | |
| Lusu Low | | 3 | 2 | |
| Elt High | 3 | 3 | 3 | |
| Elt Low | 2 | 3 | 2 | |
| Constitutive | | 3 | | |
| S505N surface | | 3 | 3 | |
| S505N constitutive | 3* | | | 3 |
| W515K surface | | | 3 | 3 |
| W515K constitutive | | | | 3 |
| Epo High | 3 | 3 | | |
| Epo Low | 3 | 3 | | |
| EpoR surface | | 3 | 3 | |

**Supplemental Table 1**. Replicate transduction numbers contributing to each dataset. *The S505N constitutive dataset has been reported previously[5].

| Variant | Treatment | Log fold change | | | | p-values | significant levels |
|---|---|---|---|---|---|---|---|
| A486M | Day0 | -0.129 | -0.078 | 0.051 | 0.019 | NA | |
| | Day2-noIL3 | -0.664 | -1.151 | -0.078 | 1.079 | 0.734 | ns |
| | Day2-IL3 | -0.356 | -0.338 | 0.181 | 0.037 | 0.437 | ns |
| T487A | Day0 | -0.063 | -0.049 | 0.012 | 0.014 | NA | |
| | Day2-noIL3 | 1.373 | 1.711 | 1.334 | 0.797 | 0.007 | ** |
| | Day2-IL3 | -0.225 | -0.178 | -0.143 | 0.008 | 0.053 | ns |
| T487W | Day0 | -0.031 | -0.076 | 0.068 | 0.069 | NA | |
| | Day2-noIL3 | 0.555 | 0.782 | 1.132 | 1.251 | 0.006 | ** |
| | Day2-IL3 | -0.254 | -0.262 | 0.143 | -0.173 | 0.147 | ns |
| W491M | Day0 | -0.131 | -0.138 | 0.100 | 0.006 | NA | |
| | Day2-noIL3 | -1.045 | -0.887 | 0.057 | 1.277 | 0.841 | ns |
| | Day2-IL3 | -0.409 | -0.391 | 0.099 | 0.043 | 0.229 | ns |
| S493V | Day0 | -0.094 | -0.156 | 0.003 | 0.023 | NA | |
| | Day2-noIL3 | 0.976 | 0.906 | 1.195 | 0.867 | 0.001 | *** |
| | Day2-IL3 | -0.339 | -0.379 | -0.057 | -0.031 | 0.066 | ns |
| S493I | Day0 | -0.160 | -0.084 | 0.156 | 0.018 | NA | |
| | Day2-noIL3 | 0.658 | 0.654 | 1.345 | 1.271 | 0.005 | ** |
| | Day2-IL3 | -0.388 | -0.309 | 0.108 | -0.013 | 0.090 | ns |
| S493M | Day0 | -0.111 | -0.067 | 0.013 | 0.011 | NA | |
| | Day2-noIL3 | 0.657 | 0.908 | 1.861 | 1.429 | 0.014 | * |
| | Day2-IL3 | -0.352 | -0.282 | -0.090 | 0.005 | 0.080 | ns |
| S493A | Day0 | -0.136 | -0.086 | 0.047 | -0.022 | NA | |
| | Day2-noIL3 | 0.528 | 1.095 | 2.181 | 1.876 | 0.022 | * |
| | Day2-IL3 | -0.377 | -0.300 | -0.056 | -0.085 | 0.037 | * |
| S493F | Day0 | -0.082 | -0.188 | 0.005 | 0.036 | NA | |
| | Day2-noIL3 | 0.033 | 0.078 | 1.428 | 0.843 | 0.115 | ns |
| | Day2-IL3 | -0.318 | -0.409 | 0.036 | 0.164 | 0.473 | ns |
| L502T | Day0 | -0.109 | -0.154 | 0.053 | -0.006 | NA | |
| | Day2-noIL3 | 1.306 | 0.880 | 1.911 | 2.790 | 0.018 | * |
| | Day2-IL3 | -0.239 | -0.318 | -0.154 | -0.136 | 0.003 | ** |
| L504K | Day0 | -0.082 | -0.140 | 0.023 | -0.036 | NA | |
| | Day2-noIL3 | -0.931 | -1.202 | 0.471 | 0.247 | 0.499 | ns |
| | Day2-IL3 | -0.319 | -0.400 | 0.031 | 0.035 | 0.304 | ns |
| S505M | Day0 | -0.126 | -0.067 | 0.078 | -0.014 | NA | |
| | Day2-noIL3 | -1.073 | -1.201 | 0.353 | -0.018 | 0.283 | ns |
| | Day2-IL3 | -0.414 | -0.364 | 0.146 | 0.080 | 0.399 | ns |
| S505N | Day0 | -0.113 | -0.103 | 0.021 | -0.060 | NA | |
| | Day2-noIL3 | 2.228 | 2.031 | 2.518 | 2.260 | 0.000 | *** |
| | Day2-IL3 | -0.118 | -0.128 | -0.225 | -0.146 | 0.195 | ns |
| L510C | Day0 | -0.060 | -0.073 | 0.025 | 0.047 | NA | |
| | Day2-noIL3 | -0.516 | -0.827 | 0.363 | 0.542 | 0.776 | ns |
| | Day2-IL3 | -0.348 | -0.327 | 0.109 | 0.233 | 0.608 | ns |
| W515K | Day0 | -0.116 | -0.078 | 0.051 | -0.014 | NA | |
| | Day2-noIL3 | 2.632 | 2.584 | 2.799 | 3.395 | 0.000 | *** |
| | Day2-IL3 | -0.156 | -0.036 | 0.001 | 0.212 | 0.538 | ns |

**Supplemental Table 2**. **Statistical analysis of potential TpoR constitutively active variants examined in Figure 9B.** Log fold change scores were calculated using the formula: $\log((Count_{Var} + 1)/(Count_{WT} + 1))$. Scores in each cell represent the mean of three technical replicates, with data from four biological replicates shown in the same row. Paired t-tests were performed to compare Day2 noIL3 / IL3 treated cell with Day0 data. Significance was denoted: p-value ≥ 0.05, ns; 0.01 ≤ p-value ≤ 0.05, *; 0.001 ≤ p-value ≤ 0.01, **; p-value ≤ 0.001, ***.

### Supplemental Materials

### DNA sequences:

TpoR:

ATGCCCTCCTGGGCCCTCTTCATGGTCACCTCCTGCCTCCTCCTGGCCCCTCAAAACCTGGCCCAAGTCAGCAGCCAAGATTATCCA
TATGATGTTCCAGATTACGCAGTCTCCTTGCTGGCATCAGACTCAGAGCCCCTGAAGTGTTTCTCCCGAACATTTGAGGACCTCACT
TGCTTCTGGGATGAGGAAGAGGCAGCGCCCAGTGGGACATACCAGCTGCTGTATGCCTACCCGCGGGAGAAGCCCCGTGCTTGCCC
CCTGAGTTCCCAGAGCATGCCCCACTTTGGAACCCGATACGTGTGCCAGTTTCCAGACCAGGAGGAAGTGCGTCTCTTCTTTCCGC
TGCACCTCTGGGTGAAGAATGTGTTCCTAAACCAGACTCGGACTCAGCGAGTCCTCTTTGTGGACAGTGTAGGCCTGCCGGCTCCC
CCCAGTATCATCAAGGCCATGGGTGGGAGCCAGCCAGGGGAACTTCAGATCAGCTGGGAGGAGCCAGCTCCAGAAATCAGTGATT
TCCTGAGGTACGAACTCCGCTATGGCCCCAGAGATCCCAAGAACTCCACTGGTCCCACGGTCATACAGCTGATTGCCACAGAAACC
TGCTGCCCTGCTCTGCAGAGGCCTCACTCAGCCTCTGCTCTGGACCAGTCTCCATGTGCTCAGCCCACAATGCCCTGGCAAGATGG
ACCAAAGCAGACCTCCCCAAGTAGAGAAGCTTCAGCTCTGACAGCAGAGGGTGGAAGCTGCCTCATCTCAGGACTCCAGCCTGGC
AACTCCTACTGGCTGCAGCTGCGCAGCGAACCTGATGGGATCTCCCTCGGTGGCTCCTGGGGATCCTGGTCCCTCCCTGTGACTGTG
GACCTGCCTGGAGATGCAGTGGCACTTGGACTGCAATGCTTTACCTTGGACCTGAAGAATGTTACCTGTCAATGGCAGCAACAGGA
CCATGCTAGCTCCCAAGGCTTCTTCTACCACAGCAGGGCACGGTGCTGCCCCAGAGACAGGTACCCCATCTGGGAGAACTGCGAAG
AGGAAGAGAAAACAAATCCAGGACTACAGACCCCACAGTTCTCTCGCTGCCACTTCAAGTCACGAAATGACAGCATTATTCACATC
CTTGTGGAGGTGACCACAGCCCCGGGTACTGTTCACAGCTACCTGGGCTCCCCTTTCTGGATCCACCAGGCTGTGCGCCTCCCCAC
CCCAAACTTGCACTGGAGGGAGATCTCCAGTGGGCATCTGGAATTGGAGTGGCAGCACCCATCGTCCTGGGCAGCCCAAGAGACC
TGTTATCAACTCCGATACACAGGAGAAGGCCATCAGGACTGGAAGGTGCTGGAGCCGCCTCTCGGGGCCCGAGGAGGGACCCTGG
AGCTGCGCCCGCGATCTCGCTACCGTTTACAGCTGCGCGCCAGGCTCAACGGCCCCACCTACCAAGGTCCCTGGAGCTCGTGGTCG
GACCCAACTAGGGTGGAGACCGCCACCGAGACCGCCTGGATCTCCTTGGTGACCGCTCTGCATCTAGTGCTGGGCCTCAGCGCCGT
CCTGGGCCTGCTGCTGCTGAGGTGGCAGTTTCCTGCACACTACAGGAGACTGAGGCATGCCCTGTGGCCCTCACTTCCAGACCTGC
ACCGGGTCCTAGGCCAGTACCTTAGGGACACTGCAGCCCTGAGCCCGCCCAAGGCCACAGTCTCAGATACCTGTGAAGAAGTGGA
ACCCAGCCTCCTTGAAATCCTCCCCAAGTCCTCAGAGAGGACTCCTTTGCCCCTGTGTTCCTCCCAGGCCCAGATGGACTACCGAA
GATTGCAGCCTTCTTGCCTGGGGACCATGCCCCTGTCTGTGTGCCCACCCATGGCTGAGTCAGGGTCCTGCTGTACCACCCACATTG
CCAACCATTCCTACCTACCACTAAGCTATTGGCAGCAGCCTTGA

S505N TpoR codon change: AGC505AAC

W515K TpoR codon change: TGG515AAG

EpoR:

ATGGACCACCTCGGGGCGTCCCTCTGGCCCCAGGTCGGCTCCCTTTGTCTCCTGCTCGCTGGGGCCGCCTGGGCCTACCCTTACGAT
GTGCCTGATTACGCCGCGCCCCCGCCTAACCTCCCGGACCCCAAGTTCGAGAGCAAAGCGGCCTTGCTGGCGGCCCGGGGGCCCG
AAGAGCTTCTGTGCTTCACCGAGCGGTTGGAGGACTTGGTGTGTTTCTGGGAGGAAGCGGCGAGCGCTGGGGTGGGCCCGGGCAA
CTACAGCTTCTCCTACCAGCTCGAGGATGAGCCATGGAAGCTGTGTCGCCTGCACCAGGCTCCCACGGCTCGTGGTGCGGTGCGCT
TCTGGTGTTCGCTGCCTACAGCCGACACGTCGAGCTTCGTGCCCCTAGAGTTGCGCGTCACAGCAGCCTCCGGCGCTCCGCGATATC
ACCGTGTCATCCACATCAATGAAGTAGTGCTCCTAGACGCCCCCGTGGGGCTGGTGGCGCGGTTGGCTGACGAGAGCGGCCACGTA
GTGTTGCGCTGGCTCCCGCCGCCTGAGACACCCATGACGTCTCACATCCGCTACGAGGTGGACGTCTCGGCCGGCAACGGCGCAG
GGAGCGTACAGAGGGTGGAGATCCTGGAGGGCCGCACCGAGTGTGTGCTGAGCAACCTGCGGGGCCGGACGCGCTACACCTTCGC
CGTCCGCGCGCGTATGGCTGAGCCGAGCTTCGGCGGCTTCTGGAGCGCCTGGTCGGAGCCTGTGTCGCTGCTGACGCCTAGCGACC
TGGACCCCCTCATCCTGACGCTCTCCCTCATCCTCGTGGTCATCCTGGTGCTGCTGACCGTGCTCGCGCTGCTCTCCCACCGCCGGG
CTCTGAAGCAGAAGATCTGGCCTGGCATCCCGAGCCCAGAGAGCGAGTTTGAAGGCCTCTTCACCACCCACAAGGGTAACTTCCA
GCTGTGGCTGTACCAGAATGATGGCTGCCTGTGGTGGAGCCCCTGCACCCCCTTCACGGAGGACCCACCTGCTTCCCTGGAAGTCC
TCTCAGAGCGCTGCTGGGGGACGATGCAGGCAGTGGAGCCGGGGACAGATGATGAGGGCCCCCTGCTGGAGCCAGTGGGCAGTG
AGCATGCCCAGGATACCTATCTGGTGCTGGACAAATGGTTGCTGCCCCGGAACCCGCCCAGTGAGGACCTCCCAGGGCCTGGTGGC
AGTGTGGACATAGTGGCCATGGATGAAGGCTCAGAAGCATCCTCCTGCTCATCTGCTTTGGCCTCGAAGCCCAGCCCAGAGGGAGC
CTCTGCTGCCAGCTTTGAGTACACTATCCTGGACCCCAGCTCCCAGCTCTTGCGTCCATGGACACTGTGCCCTGAGCTGCCCCCTAC
CCCACCCCACCTAAAGTACCTGTACCTTGTGGTATCTGACTCTGGCATCTCAACTGACTACAGCTCAGGGGACTCCCAGGGAGCCCA
AGGGGGCTTATCCGATGGCCCCTACTCCAACCCTTATGAGAACAGCCTTATCCCAGCCGCTGAGCCTCTGCCCCCCAGCTATGTGGC
TTGCTCTTGA

### Vector information:

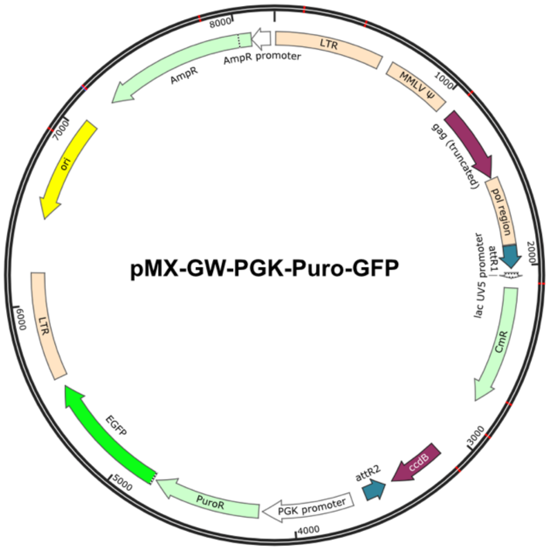


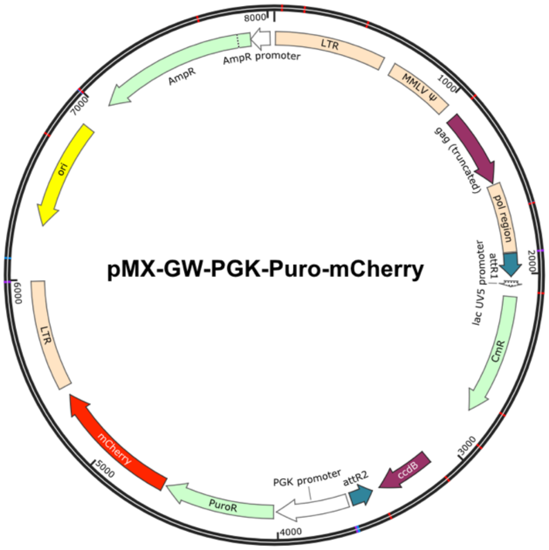


**Protein sequence used for AF3 models**:

### hTpoR:

VRLPTPNLHWREISSGHLELEWQHPSSWAAQETCYQLRYTGEGHQDWKVLEPPLGARGGTLELRPRSRYRLQLRARLNGPTYQGPWS
SWSDPTRVETATETAWISLVTALHLVLGLSAVLGLLLLRWQFPAHYRRLRHALWPSLPDLHRVLGQYLRDTAALSPPKATVSDTCEEV
EPSLLE

### hJAK2:

DPVLQVYLYHSLGKSEADYLTFPSGEYVAEEICIAASKACGITPVYHNMFALMSETERIWYPPNHVFHIDESTRHNVLYRIRFYFPRWYC
SGSNRAYRHGISRGAEAPLLDDFVMSYLFAQWRHDFVHGWIKVPVTHETQEECLGMAVLDMMRIAKENDQTPLAIYNSISYKTFLPKC
IRAKIQDYHILTRKRIRYRFRRFIQQFSQCKATARNLKLKYLINLETLQSAFYTEKFEVKEPGSGPSGEEIFATIIITGNGGIQWSRGKHKES
ETLTEQDLQLYCDFPNIIDVSIKQANQEGSNESRVVTIHKQDGKNLEIELSSLREALSFVSLIDGYYRLTADAHHYLCKEVAPPAVLENIQ
SNCHGPISMDFAISKLKKAGNQTGLYVLRCSPKDFNKYFLTFAVERENVIEYKHCLITKNENEEYNLSGTKKNFSSLKDLLNCYQMETV
RSDNIIFQFTKCCPPKPKDKSNLLVFRTNGVSDVPTSPTLQRPTHMNQMVFHKIRNEDLIFNESLGQGTFTKIFKGVRREVGDYGQLHET
EVLLKVLDKAHRNYSESFFEAASMMSKLSHKHLVLNYGVCVCGDENILVQEFVKFGSLDTYLKKNKNCINILWKLEVAKQLAWAMHF
LEENTLIHGNVCAKNILLIREEDRKTGNPPFIKLSDPGISITVLPKDILQERIPWVPPECIENPKNLNLATDKWSFGTTLWEICSGGDKPLSA
LDSQRKLQFYEDRHQLPAPKWAELANLINNCMDYEPDFRPSFRAIIRDLNSLFTPDYELLTENDMLPNMRIGALGFSGAFEDRDP

### hEpoR:

DPKFESKAALLAARGPEELLCFTERLEDLVCFWEEAASAGVGPGNYSFSYQLEDEPWKLCRLHQAPTARGAVRFWCSLPTADTSSFVPL
ELRVTAASGAPRYHRVIHINEVVLLDAPVGLVARLADESGHVVLRWLPPPETPMTSHIRYEVDVSAGNGAGSVQRVEILEGRTECVLSN
LRGRTRYTFAVRARMAEPSFGGFWSAWSEPVSLLTPSDLDPLILTLSLILVVILVLLTVLALLSHRRALKQKIWPGIPSPESEFEGLFTTHK
GNFQLWLYQNDGCLWWSPCTPFTEDPPASLE

### hEpo:

APPRLICDSRVLERYLLEAKEAENITTGCAEHCSLNENITVPDTKVNFYAWKRMEVGQQAVEVWQGLALLSEAVLRGQALLVNSSQPWE
PLQLHVDKAVSGLRSLTTLLRALGAQKEAISPPDAASAAPLRTITADTFRKLFRVYSNFLRGKLKLYTGEACRTGDR

**Oligos for constructing DMS libraries:**
**1. wildtype TpoR as base sequence**

| | Invariant fragment | | Fragment with degenerate codon | |
|---|---|---|---|---|
| | Forward oligo | Reverse oligo | Forward oligo | Reverse oligo |
| T481X | TGTAAAACGACGGCCAGTCTTAAG | GGGTCCGACCACGAGCTC | GAGCTCGTGGTCGGACCCANNNAGGGTGGAGACCGCCACC | GCTATGACCATGTAATACGACTCACTATAGGGG |
| R482X | TGTAAAACGACGGCCAGTCTTAAG | GGGTCCGACCACGAGCTC | GAGCTCGTGGTCGGACCCAACTNNNGTGGAGACCGCCACCGAG | GCTATGACCATGTAATACGACTCACTATAGGGG |
| V483X | TGTAAAACGACGGCCAGTCTTAAG | GGGTCCGACCACGAGCTC | GAGCTCGTGGTCGGACCCAACTAGGNNNGAGACCGCCACCGAGACC | GCTATGACCATGTAATACGACTCACTATAGGGG |
| E484X | TGTAAAACGACGGCCAGTCTTAAG | GGGTCCGACCACGAGCTC | GAGCTCGTGGTCGGACCCAACTAGGGTGNNNACCGCCACCGAGACCGCC | GCTATGACCATGTAATACGACTCACTATAGGGG |
| T485X | TGTAAAACGACGGCCAGTCTTAAG | GGGTCCGACCACGAGCTC | GAGCTCGTGGTCGGACCCAACTAGGGTGGAGNNNGCCACCGAGACCGCCTGG | GCTATGACCATGTAATACGACTCACTATAGGGG |
| A486X | TGTAAAACGACGGCCAGTCTTAAG | GGGTCCGACCACGAGCTC | GAGCTCGTGGTCGGACCCAACTAGGGTGGAGACCNNNACCGAGACCGCCTGGATC | GCTATGACCATGTAATACGACTCACTATAGGGG |
| T487X | TGTAAAACGACGGCCAGTCTTAAG | GGGTCCGACCACGAGCTC | GAGCTCGTGGTCGGACCCAACTAGGGTGGAGACCGCCNNNGAGACCGCCTGGATCTCC | GCTATGACCATGTAATACGACTCACTATAGGGG |
| E488X | TGTAAAACGACGGCCAGTCTTAAG | GGTGGCGGTCTCCACCCTAG | CTAGGGTGGAGACCGCCACCNNNACCGCCTGGATCTCCTTG | GCTATGACCATGTAATACGACTCACTATAGGGG |
| T489X | TGTAAAACGACGGCCAGTCTTAAG | GGTGGCGGTCTCCACCCTAG | CTAGGGTGGAGACCGCCACCGAGNNNGCCTGGATCTCCTTGGTG | GCTATGACCATGTAATACGACTCACTATAGGGG |
| A490X | TGTAAAACGACGGCCAGTCTTAAG | GGTGGCGGTCTCCACCCTAG | CTAGGGTGGAGACCGCCACCGAGACCNNNTGGATCTCCTTGGTGACC | GCTATGACCATGTAATACGACTCACTATAGGGG |
| W491X | TGTAAAACGACGGCCAGTCTTAAG | GGTGGCGGTCTCCACCCTAG | CTAGGGTGGAGACCGCCACCGAGACCGCCNNNATCTCCTTGGTGACCGCTC | GCTATGACCATGTAATACGACTCACTATAGGGG |
| I492X | TGTAAAACGACGGCCAGTCTTAAG | GGTGGCGGTCTCCACCCTAG | CTAGGGTGGAGACCGCCACCGAGACCGCCTGGNNNTCCTTGGTGACCGCTCTG | GCTATGACCATGTAATACGACTCACTATAGGGG |
| S493X | TGTAAAACGACGGCCAGTCTTAAG | GGTGGCGGTCTCCACCCTAG | CTAGGGTGGAGACCGCCACCGAGACCGCCTGGATCNNNTTGGTGACCGCTCTGCATC | GCTATGACCATGTAATACGACTCACTATAGGGG |
| L494X | TGTAAAACGACGGCCAGTCTTAAG | GGTGGCGGTCTCCACCCTAG | CTAGGGTGGAGACCGCCACCGAGACCGCCTGGATCTCCNNNGTGACCGCTCTGCATCTAG | GCTATGACCATGTAATACGACTCACTATAGGGG |

| | | | | |
|---|---|---|---|---|
| V495X | TGTAAAACGACGGCCAGTCTTAAG | CAAGGAGATCCAGGCGGTC | GACCGCCTGGATCTCCTTGNNNACCGCTCTGCATCTAGTG | GCTATGACCATGTAATACGACTCACTATAGGGG |
| T496X | TGTAAAACGACGGCCAGTCTTAAG | CAAGGAGATCCAGGCGGTC | GACCGCCTGGATCTCCTTGGTGNNNGCTCTGCATCTAGTGCTG | GCTATGACCATGTAATACGACTCACTATAGGGG |
| A497X | TGTAAAACGACGGCCAGTCTTAAG | CAAGGAGATCCAGGCGGTC | GACCGCCTGGATCTCCTTGGTGACCNNNCTGCATCTAGTGCTGGGC | GCTATGACCATGTAATACGACTCACTATAGGGG |
| L498X | TGTAAAACGACGGCCAGTCTTAAG | CAAGGAGATCCAGGCGGTC | GACCGCCTGGATCTCCTTGGTGACCGCTNNNCATCTAGTGCTGGGCCTC | GCTATGACCATGTAATACGACTCACTATAGGGG |
| H499X | TGTAAAACGACGGCCAGTCTTAAG | CAAGGAGATCCAGGCGGTC | GACCGCCTGGATCTCCTTGGTGACCGCTCTGNNNCTAGTGCTGGGCCTCAGC | GCTATGACCATGTAATACGACTCACTATAGGGG |
| L500X | TGTAAAACGACGGCCAGTCTTAAG | CAAGGAGATCCAGGCGGTC | GACCGCCTGGATCTCCTTGGTGACCGCTCTGCATNNNGTGCTGGGCCTCAGCGCC | GCTATGACCATGTAATACGACTCACTATAGGGG |
| V501X | TGTAAAACGACGGCCAGTCTTAAG | CAAGGAGATCCAGGCGGTC | GACCGCCTGGATCTCCTTGGTGACCGCTCTGCATCTANNNCTGGGCCTCAGCGCCGTC | GCTATGACCATGTAATACGACTCACTATAGGGG |
| L502X | TGTAAAACGACGGCCAGTCTTAAG | CACTAGATGCAGAGCGGTC | GACCGCTCTGCATCTAGTGNNNGGCCTCAGCGCCGTCCTG | GCTATGACCATGTAATACGACTCACTATAGGGG |
| G503X | TGTAAAACGACGGCCAGTCTTAAG | CACTAGATGCAGAGCGGTC | GACCGCTCTGCATCTAGTGCTGNNNCTCAGCGCCGTCCTGGGC | GCTATGACCATGTAATACGACTCACTATAGGGG |
| L504X | TGTAAAACGACGGCCAGTCTTAAG | CACTAGATGCAGAGCGGTC | GACCGCTCTGCATCTAGTGCTGGGCNNNAGCGCCGTCCTGGGCCTG | GCTATGACCATGTAATACGACTCACTATAGGGG |
| S505X | TGTAAAACGACGGCCAGTCTTAAG | CACTAGATGCAGAGCGGTC | GACCGCTCTGCATCTAGTGCTGGGCCTCNNNGCCGTCCTGGGCCTGCTG | GCTATGACCATGTAATACGACTCACTATAGGGG |
| A506X | TGTAAAACGACGGCCAGTCTTAAG | CACTAGATGCAGAGCGGTC | GACCGCTCTGCATCTAGTGCTGGGCCTCAGCNNNGTCCTGGGCCTGCTGCTG | GCTATGACCATGTAATACGACTCACTATAGGGG |
| V507X | TGTAAAACGACGGCCAGTCTTAAG | CACTAGATGCAGAGCGGTC | GACCGCTCTGCATCTAGTGCTGGGCCTCAGCGCCNNNCTGGGCCTGCTGCTGCTG | GCTATGACCATGTAATACGACTCACTATAGGGG |
| L508X | TGTAAAACGACGGCCAGTCTTAAG | CACTAGATGCAGAGCGGTC | GACCGCTCTGCATCTAGTGCTGGGCCTCAGCGCCGTCNNNGGCCTGCTGCTGCTGAGG | GCTATGACCATGTAATACGACTCACTATAGGGG |
| G509X | TGTAAAACGACGGCCAGTCTTAAG | CAGGACGGCGCTGAGGCC | GGCCTCAGCGCCGTCCTGNNNCTGCTGCTGCTGAGGTGG | GCTATGACCATGTAATACGACTCACTATAGGGG |
| L510X | TGTAAAACGACGGCCAGTCTTAAG | CAGGACGGCGCTGAGGCC | GGCCTCAGCGCCGTCCTGGGCNNNCTGCTGCTGAGGTGGCAG | GCTATGACCATGTAATACGACTCACTATAGGGG |

| | | | | |
|---|---|---|---|---|
| L511X | TGTAAAACGACGGCCAGTCTTAAG | CAGGACGGCGCTGAGGCC | GGCCTCAGCGCCGTCCTGGGCCTGNNNCTGCTGAGGTGGCAGTTTC | GCTATGACCATGTAATACGACTCACTATAGGGG |
| L512X | TGTAAAACGACGGCCAGTCTTAAG | CAGGACGGCGCTGAGGCC | GGCCTCAGCGCCGTCCTGGGCCTGCTGNNNCTGAGGTGGCAGTTTCCTG | GCTATGACCATGTAATACGACTCACTATAGGGG |
| L513X | TGTAAAACGACGGCCAGTCTTAAG | CAGGACGGCGCTGAGGCC | GGCCTCAGCGCCGTCCTGGGCCTGCTGCTGNNNAGGTGGCAGTTTCCTGCAC | GCTATGACCATGTAATACGACTCACTATAGGGG |
| R514X | TGTAAAACGACGGCCAGTCTTAAG | CAGGACGGCGCTGAGGCC | GGCCTCAGCGCCGTCCTGGGCCTGCTGCTGCTGNNNTGGCAGTTTCCTGCACAC | GCTATGACCATGTAATACGACTCACTATAGGGG |
| W515X | TGTAAAACGACGGCCAGTCTTAAG | CAGGACGGCGCTGAGGCC | GGCCTCAGCGCCGTCCTGGGCCTGCTGCTGCTGAGGNNNCAGTTTCCTGCACACTAC | GCTATGACCATGTAATACGACTCACTATAGGGG |
| Q516X | TGTAAAACGACGGCCAGTCTTAAG | CAGGACGGCGCTGAGGCC | GGCCTCAGCGCCGTCCTGGGCCTGCTGCTGCTGAGGTGGNNNTTTCCTGCACACTACAGG | GCTATGACCATGTAATACGACTCACTATAGGGG |
| F517X | TGTAAAACGACGGCCAGTCTTAAG | CTGCCACCTCAGCAGCAG | CTGCTGCTGAGGTGGCAGNNNCCTGCACACTACAGGAGAC | GCTATGACCATGTAATACGACTCACTATAGGGG |
| P518X | TGTAAAACGACGGCCAGTCTTAAG | CTGCCACCTCAGCAGCAG | CTGCTGCTGAGGTGGCAGTTTNNNGCACACTACAGGAGACTG | GCTATGACCATGTAATACGACTCACTATAGGGG |
| A519X | TGTAAAACGACGGCCAGTCTTAAG | CTGCCACCTCAGCAGCAG | CTGCTGCTGAGGTGGCAGTTTCCTNNNCACTACAGGAGACTGAGG | GCTATGACCATGTAATACGACTCACTATAGGGG |
| H520X | TGTAAAACGACGGCCAGTCTTAAG | CTGCCACCTCAGCAGCAG | CTGCTGCTGAGGTGGCAGTTTCCTGCANNNTACAGGAGACTGAGGCATG | GCTATGACCATGTAATACGACTCACTATAGGGG |
| Y521X | TGTAAAACGACGGCCAGTCTTAAG | CTGCCACCTCAGCAGCAG | CTGCTGCTGAGGTGGCAGTTTCCTGCACACNNNAGGAGACTGAGGCATGCC | GCTATGACCATGTAATACGACTCACTATAGGGG |
| R522X | TGTAAAACGACGGCCAGTCTTAAG | CTGCCACCTCAGCAGCAG | CTGCTGCTGAGGTGGCAGTTTCCTGCACACTACNNNAGACTGAGGCATGCCCTG | GCTATGACCATGTAATACGACTCACTATAGGGG |
| R523X | TGTAAAACGACGGCCAGTCTTAAG | CTGCCACCTCAGCAGCAG | CTGCTGCTGAGGTGGCAGTTTCCTGCACACTACAGGNNNCTGAGGCATGCCCTGTGG | GCTATGACCATGTAATACGACTCACTATAGGGG |
| L524X | TGTAAAACGACGGCCAGTCTTAAG | CTCCTGTAGTGTGCAGG | CCTGCACACTACAGGAGANNNAGGCATGCCCTGTGGCCC | GCTATGACCATGTAATACGACTCACTATAGGGG |
| R525X | TGTAAAACGACGGCCAGTCTTAAG | CTCCTGTAGTGTGCAGG | CCTGCACACTACAGGAGACTGNNNCATGCCCTGTGGCCCTCAC | GCTATGACCATGTAATACGACTCACTATAGGGG |
| H526X | TGTAAAACGACGGCCAGTCTTAAG | CTCCTGTAGTGTGCAGG | CCTGCACACTACAGGAGACTGAGGNNNGCCCTGTGGCCCTCACTTC | GCTATGACCATGTAATACGACTCACTATAGGGG |

| | | | | |
|---|---|---|---|---|
| A527X | TGTAAAACGACGGCCAGTCTTAAG | CTCCTGTAGTGTGCAGG | CCTGCACACTACAGGAGACTGAGGCATNNNCTGTGGCCCTCACTTCCAG | GCTATGACCATGTAATACGACTCACTATAGGGG |
| L528X | TGTAAAACGACGGCCAGTCTTAAG | CTCCTGTAGTGTGCAGG | CCTGCACACTACAGGAGACTGAGGCATGCCNNNTGGCCCTCACTTCCAGAC | GCTATGACCATGTAATACGACTCACTATAGGGG |
| W529X | TGTAAAACGACGGCCAGTCTTAAG | CTCCTGTAGTGTGCAGG | CCTGCACACTACAGGAGACTGAGGCATGCCCTGNNNCCCTCACTTCCAGACCTG | GCTATGACCATGTAATACGACTCACTATAGGGG |

### 2. S505N TpoR as base sequence

| | backbone | | Oligo pools |
|---|---|---|---|
| Library | Forward oligo | Reverse oligo | |
| L2 | CTGCTGAGGTGGCAGTTTCCTGCAC | TGGGTCCGACCACGAGCTCCAGGGA | TCCCTGGAGCTCGTGGTCGGACCCANNNAGGGTGGAGACCGCCACCG<br>AGACCGCCTGGATCTCCTTGGTGACCGCTCTGCATCTAGTGCTGGGCC<br>TCAACGCCGTCCTGGGCCTGCTGCTGCTGAGGTGGCAGTTTCCTGCAC<br>TCCCTGGAGCTCGTGGTCGGACCCAACTNNNGTGGAGACCGCCACCG<br>AGACCGCCTGGATCTCCTTGGTGACCGCTCTGCATCTAGTGCTGGGCC<br>TCAACGCCGTCCTGGGCCTGCTGCTGCTGAGGTGGCAGTTTCCTGCAC<br>TCCCTGGAGCTCGTGGTCGGACCCAACTAGGNNNGAGACCGCCACCG<br>AGACCGCCTGGATCTCCTTGGTGACCGCTCTGCATCTAGTGCTGGGCC<br>TCAACGCCGTCCTGGGCCTGCTGCTGCTGAGGTGGCAGTTTCCTGCAC<br>TCCCTGGAGCTCGTGGTCGGACCCAACTAGGGTGNNNACCGCCACCG<br>AGACCGCCTGGATCTCCTTGGTGACCGCTCTGCATCTAGTGCTGGGCC<br>TCAACGCCGTCCTGGGCCTGCTGCTGCTGAGGTGGCAGTTTCCTGCAC<br>TCCCTGGAGCTCGTGGTCGGACCCAACTAGGGTGGAGNNNGCCACCG<br>AGACCGCCTGGATCTCCTTGGTGACCGCTCTGCATCTAGTGCTGGGCC<br>TCAACGCCGTCCTGGGCCTGCTGCTGCTGAGGTGGCAGTTTCCTGCAC<br>TCCCTGGAGCTCGTGGTCGGACCCAACTAGGGTGGAGACCNNNACCG<br>AGACCGCCTGGATCTCCTTGGTGACCGCTCTGCATCTAGTGCTGGGCC<br>TCAACGCCGTCCTGGGCCTGCTGCTGCTGAGGTGGCAGTTTCCTGCAC<br>TCCCTGGAGCTCGTGGTCGGACCCAACTAGGGTGGAGACCGCCNNNG<br>AGACCGCCTGGATCTCCTTGGTGACCGCTCTGCATCTAGTGCTGGGCC<br>TCAACGCCGTCCTGGGCCTGCTGCTGCTGAGGTGGCAGTTTCCTGCAC<br>TCCCTGGAGCTCGTGGTCGGACCCAACTAGGGTGGAGACCGCCACCN<br>NNACCGCCTGGATCTCCTTGGTGACCGCTCTGCATCTAGTGCTGGGCC<br>TCAACGCCGTCCTGGGCCTGCTGCTGCTGAGGTGGCAGTTTCCTGCAC<br>TCCCTGGAGCTCGTGGTCGGACCCAACTAGGGTGGAGACCGCCACCG<br>AGNNNGCCTGGATCTCCTTGGTGACCGCTCTGCATCTAGTGCTGGGCC<br>TCAACGCCGTCCTGGGCCTGCTGCTGCTGAGGTGGCAGTTTCCTGCAC<br>TCCCTGGAGCTCGTGGTCGGACCCAACTAGGGTGGAGACCGCCACCG<br>AGACCNNNTGGATCTCCTTGGTGACCGCTCTGCATCTAGTGCTGGGCC<br>TCAACGCCGTCCTGGGCCTGCTGCTGCTGAGGTGGCAGTTTCCTGCAC<br>TCCCTGGAGCTCGTGGTCGGACCCAACTAGGGTGGAGACCGCCACCG<br>AGACCGCCNNNATCTCCTTGGTGACCGCTCTGCATCTAGTGCTGGGCC<br>TCAACGCCGTCCTGGGCCTGCTGCTGCTGAGGTGGCAGTTTCCTGCAC<br>TCCCTGGAGCTCGTGGTCGGACCCAACTAGGGTGGAGACCGCCACCG<br>AGACCGCCTGGNNNTCCTTGGTGACCGCTCTGCATCTAGTGCTGGGCC<br>TCAACGCCGTCCTGGGCCTGCTGCTGCTGAGGTGGCAGTTTCCTGCAC<br>TCCCTGGAGCTCGTGGTCGGACCCAACTAGGGTGGAGACCGCCACCG<br>AGACCGCCTGGATCNNNTTGGTGACCGCTCTGCATCTAGTGCTGGGCC<br>TCAACGCCGTCCTGGGCCTGCTGCTGCTGAGGTGGCAGTTTCCTGCAC<br>TCCCTGGAGCTCGTGGTCGGACCCAACTAGGGTGGAGACCGCCACCG<br>AGACCGCCTGGATCTCCNNNGTGACCGCTCTGCATCTAGTGCTGGGCC<br>TCAACGCCGTCCTGGGCCTGCTGCTGCTGAGGTGGCAGTTTCCTGCAC<br>TCCCTGGAGCTCGTGGTCGGACCCAACTAGGGTGGAGACCGCCACCG<br>AGACCGCCTGGATCTCCTTGNNNACCGCTCTGCATCTAGTGCTGGGCC<br>TCAACGCCGTCCTGGGCCTGCTGCTGCTGAGGTGGCAGTTTCCTGCAC<br>TCCCTGGAGCTCGTGGTCGGACCCAACTAGGGTGGAGACCGCCACCG<br>AGACCGCCTGGATCTCCTTGGTGNNNGCTCTGCATCTAGTGCTGGGCC<br>TCAACGCCGTCCTGGGCCTGCTGCTGCTGAGGTGGCAGTTTCCTGCAC<br>TCCCTGGAGCTCGTGGTCGGACCCAACTAGGGTGGAGACCGCCACCG<br>AGACCGCCTGGATCTCCTTGGTGACCNNNCTGCATCTAGTGCTGGGCC<br>TCAACGCCGTCCTGGGCCTGCTGCTGCTGAGGTGGCAGTTTCCTGCAC<br>TCCCTGGAGCTCGTGGTCGGACCCAACTAGGGTGGAGACCGCCACCG<br>AGACCGCCTGGATCTCCTTGGTGACCGCTNNNCATCTAGTGCTGGGCC<br>TCAACGCCGTCCTGGGCCTGCTGCTGCTGAGGTGGCAGTTTCCTGCAC |

| | | | |
|---|---|---|---|
| | | | TCCCTGGAGCTCGTGGTCGGACCCAACTAGGGTGGAGACCGCCACCG<br>AGACCGCCTGGATCTCCTTGGTGACCGCTCTGNNNCTAGTGCTGGGCC<br>TCAACGCCGTCCTGGGCCTGCTGCTGCTGAGGTGGCAGTTTCCTGCAC<br>TCCCTGGAGCTCGTGGTCGGACCCAACTAGGGTGGAGACCGCCACCG<br>AGACCGCCTGGATCTCCTTGGTGACCGCTCTGCATNNNGTGCTGGGCC<br>TCAACGCCGTCCTGGGCCTGCTGCTGCTGAGGTGGCAGTTTCCTGCAC<br>TCCCTGGAGCTCGTGGTCGGACCCAACTAGGGTGGAGACCGCCACCG<br>AGACCGCCTGGATCTCCTTGGTGACCGCTCTGCATCTANNNCTGGGCC<br>TCAACGCCGTCCTGGGCCTGCTGCTGCTGAGGTGGCAGTTTCCTGCAC<br>TCCCTGGAGCTCGTGGTCGGACCCAACTAGGGTGGAGACCGCCACCG<br>AGACCGCCTGGATCTCCTTGGTGACCGCTCTGCATCTAGTGNNNGGCC<br>TCAACGCCGTCCTGGGCCTGCTGCTGCTGAGGTGGCAGTTTCCTGCAC<br>TCCCTGGAGCTCGTGGTCGGACCCAACTAGGGTGGAGACCGCCACCG<br>AGACCGCCTGGATCTCCTTGGTGACCGCTCTGCATCTAGTGCTGNNNC<br>TCAACGCCGTCCTGGGCCTGCTGCTGCTGAGGTGGCAGTTTCCTGCAC<br>TCCCTGGAGCTCGTGGTCGGACCCAACTAGGGTGGAGACCGCCACCG<br>AGACCGCCTGGATCTCCTTGGTGACCGCTCTGCATCTAGTGCTGGGCN<br>NNAACGCCGTCCTGGGCCTGCTGCTGCTGAGGTGGCAGTTTCCTGCAC<br>TCCCTGGAGCTCGTGGTCGGACCCAACTAGGGTGGAGACCGCCACCG<br>AGACCGCCTGGATCTCCTTGGTGACCGCTCTGCATCTAGTGCTGGGCC<br>TCNNNGCCGTCCTGGGCCTGCTGCTGCTGAGGTGGCAGTTTCCTGCAC<br>TCCCTGGAGCTCGTGGTCGGACCCAACTAGGGTGGAGACCGCCACCG<br>AGACCGCCTGGATCTCCTTGGTGACCGCTCTGCATCTAGTGCTGGGCC<br>TCAACNNNGTCCTGGGCCTGCTGCTGCTGAGGTGGCAGTTTCCTGCAC<br>TCCCTGGAGCTCGTGGTCGGACCCAACTAGGGTGGAGACCGCCACCG<br>AGACCGCCTGGATCTCCTTGGTGACCGCTCTGCATCTAGTGCTGGGCC<br>TCAACGCCNNNCTGGGCCTGCTGCTGCTGAGGTGGCAGTTTCCTGCAC<br>TCCCTGGAGCTCGTGGTCGGACCCAACTAGGGTGGAGACCGCCACCG<br>AGACCGCCTGGATCTCCTTGGTGACCGCTCTGCATCTAGTGCTGGGCC<br>TCAACGCCGTCNNNGGCCTGCTGCTGCTGAGGTGGCAGTTTCCTGCAC<br>TCCCTGGAGCTCGTGGTCGGACCCAACTAGGGTGGAGACCGCCACCG<br>AGACCGCCTGGATCTCCTTGGTGACCGCTCTGCATCTAGTGCTGGGCC<br>TCAACGCCGTCCTGNNNCTGCTGCTGCTGAGGTGGCAGTTTCCTGCAC<br>TCCCTGGAGCTCGTGGTCGGACCCAACTAGGGTGGAGACCGCCACCG<br>AGACCGCCTGGATCTCCTTGGTGACCGCTCTGCATCTAGTGCTGGGCC<br>TCAACGCCGTCCTGGGCNNNCTGCTGCTGAGGTGGCAGTTTCCTGCAC<br>TCCCTGGAGCTCGTGGTCGGACCCAACTAGGGTGGAGACCGCCACCG<br>AGACCGCCTGGATCTCCTTGGTGACCGCTCTGCATCTAGTGCTGGGCC<br>TCAACGCCGTCCTGGGCCTGNNNCTGCTGAGGTGGCAGTTTCCTGCAC |
| L3 | CCCTCACT<br>TCCAGACC<br>TGCACCGG<br>G | GAGGCCCA<br>GCACTAGA<br>TGCAGAGC<br>G | CGCTCTGCATCTAGTGCTGGGCCTCAACGCCNNNCTGGGCCTGCTGCT<br>GCTGAGGTGGCAGTTTCCTGCACACTACAGGAGACTGAGGCATGCCCT<br>GTGGCCCTCACTTCCAGACCTGCACCGGG<br>CGCTCTGCATCTAGTGCTGGGCCTCAACGCCGTCNNNGGCCTGCTGCT<br>GCTGAGGTGGCAGTTTCCTGCACACTACAGGAGACTGAGGCATGCCCT<br>GTGGCCCTCACTTCCAGACCTGCACCGGG<br>CGCTCTGCATCTAGTGCTGGGCCTCAACGCCGTCCTGNNNCTGCTGCT<br>GCTGAGGTGGCAGTTTCCTGCACACTACAGGAGACTGAGGCATGCCCT<br>GTGGCCCTCACTTCCAGACCTGCACCGGG<br>CGCTCTGCATCTAGTGCTGGGCCTCAACGCCGTCCTGGGCNNNCTGCT<br>GCTGAGGTGGCAGTTTCCTGCACACTACAGGAGACTGAGGCATGCCCT<br>GTGGCCCTCACTTCCAGACCTGCACCGGG<br>CGCTCTGCATCTAGTGCTGGGCCTCAACGCCGTCCTGGGCCTGNNNCT<br>GCTGAGGTGGCAGTTTCCTGCACACTACAGGAGACTGAGGCATGCCCT<br>GTGGCCCTCACTTCCAGACCTGCACCGGG<br>CGCTCTGCATCTAGTGCTGGGCCTCAACGCCGTCCTGGGCCTGCTGNN<br>NCTGAGGTGGCAGTTTCCTGCACACTACAGGAGACTGAGGCATGCCCT<br>GTGGCCCTCACTTCCAGACCTGCACCGGG<br>CGCTCTGCATCTAGTGCTGGGCCTCAACGCCGTCCTGGGCCTGCTGCT<br>GNNNAGGTGGCAGTTTCCTGCACACTACAGGAGACTGAGGCATGCCCT<br>GTGGCCCTCACTTCCAGACCTGCACCGGG |

| | | | |
|---|---|---|---|
| | | | CGCTCTGCATCTAGTGCTGGGCCTCAACGCCGTCCTGGGCCTGCTGCT<br>GCTGNNNTGGCAGTTTCCTGCACACTACAGGAGACTGAGGCATGCCCT<br>GTGGCCCTCACTTCCAGACCTGCACCGGG<br>CGCTCTGCATCTAGTGCTGGGCCTCAACGCCGTCCTGGGCCTGCTGCT<br>GCTGAGGNNNCAGTTTCCTGCACACTACAGGAGACTGAGGCATGCCCT<br>GTGGCCCTCACTTCCAGACCTGCACCGGG<br>CGCTCTGCATCTAGTGCTGGGCCTCAACGCCGTCCTGGGCCTGCTGCT<br>GCTGAGGTGGNNNTTTCCTGCACACTACAGGAGACTGAGGCATGCCCT<br>GTGGCCCTCACTTCCAGACCTGCACCGGG<br>CGCTCTGCATCTAGTGCTGGGCCTCAACGCCGTCCTGGGCCTGCTGCT<br>GCTGAGGTGGCAGNNNCCTGCACACTACAGGAGACTGAGGCATGCCCT<br>GTGGCCCTCACTTCCAGACCTGCACCGGG<br>CGCTCTGCATCTAGTGCTGGGCCTCAACGCCGTCCTGGGCCTGCTGCT<br>GCTGAGGTGGCAGTTTNNNGCACACTACAGGAGACTGAGGCATGCCCT<br>GTGGCCCTCACTTCCAGACCTGCACCGGG<br>CGCTCTGCATCTAGTGCTGGGCCTCAACGCCGTCCTGGGCCTGCTGCT<br>GCTGAGGTGGCAGTTTCCTNNNCACTACAGGAGACTGAGGCATGCCCT<br>GTGGCCCTCACTTCCAGACCTGCACCGGG<br>CGCTCTGCATCTAGTGCTGGGCCTCAACGCCGTCCTGGGCCTGCTGCT<br>GCTGAGGTGGCAGTTTCCTGCANNNTACAGGAGACTGAGGCATGCCCT<br>GTGGCCCTCACTTCCAGACCTGCACCGGG<br>CGCTCTGCATCTAGTGCTGGGCCTCAACGCCGTCCTGGGCCTGCTGCT<br>GCTGAGGTGGCAGTTTCCTGCACACNNNAGGAGACTGAGGCATGCCCT<br>GTGGCCCTCACTTCCAGACCTGCACCGGG<br>CGCTCTGCATCTAGTGCTGGGCCTCAACGCCGTCCTGGGCCTGCTGCT<br>GCTGAGGTGGCAGTTTCCTGCACACTACNNNAGACTGAGGCATGCCCT<br>GTGGCCCTCACTTCCAGACCTGCACCGGG<br>CGCTCTGCATCTAGTGCTGGGCCTCAACGCCGTCCTGGGCCTGCTGCT<br>GCTGAGGTGGCAGTTTCCTGCACACTACAGGNNNCTGAGGCATGCCCT<br>GTGGCCCTCACTTCCAGACCTGCACCGGG<br>CGCTCTGCATCTAGTGCTGGGCCTCAACGCCGTCCTGGGCCTGCTGCT<br>GCTGAGGTGGCAGTTTCCTGCACACTACAGGAGANNNAGGCATGCCCT<br>GTGGCCCTCACTTCCAGACCTGCACCGGG<br>CGCTCTGCATCTAGTGCTGGGCCTCAACGCCGTCCTGGGCCTGCTGCT<br>GCTGAGGTGGCAGTTTCCTGCACACTACAGGAGACTGNNNCATGCCCT<br>GTGGCCCTCACTTCCAGACCTGCACCGGG<br>CGCTCTGCATCTAGTGCTGGGCCTCAACGCCGTCCTGGGCCTGCTGCT<br>GCTGAGGTGGCAGTTTCCTGCACACTACAGGAGACTGAGGNNNGCCCT<br>GTGGCCCTCACTTCCAGACCTGCACCGGG<br>CGCTCTGCATCTAGTGCTGGGCCTCAACGCCGTCCTGGGCCTGCTGCT<br>GCTGAGGTGGCAGTTTCCTGCACACTACAGGAGACTGAGGCATNNNCT<br>GTGGCCCTCACTTCCAGACCTGCACCGGG<br>CGCTCTGCATCTAGTGCTGGGCCTCAACGCCGTCCTGGGCCTGCTGCT<br>GCTGAGGTGGCAGTTTCCTGCACACTACAGGAGACTGAGGCATGCCNN<br>NTGGCCCTCACTTCCAGACCTGCACCGGG<br>CGCTCTGCATCTAGTGCTGGGCCTCAACGCCGTCCTGGGCCTGCTGCT<br>GCTGAGGTGGCAGTTTCCTGCACACTACAGGAGACTGAGGCATGCCCT<br>GNNNCCCTCACTTCCAGACCTGCACCGGG |

#### 3. W515K TpoR as base sequence

| | backbone | | Oligo pools |
|---|---|---|---|
| Library | Forward oligo | Reverse oligo | |
| L2 | CAGTTTCCTGCACACTACAGGAGAC | TGGGTCCGACCACGAGCTCCAGGGA | TCCCTGGAGCTCGTGGTCGGACCCANNNAGGGTGGAGACCGCCACCGAGACCGCCTGGATCTCCTTGGTGACCGCTCTGCATCTAGTGCTGGGCCTCAGCGCCGTCCTGGGCCTGCTGCTGCTGAGGAAGCAGTTTCCTGCACACTACAGGAGAC<br>TCCCTGGAGCTCGTGGTCGGACCCAACTNNNGTGGAGACCGCCACCGAGACCGCCTGGATCTCCTTGGTGACCGCTCTGCATCTAGTGCTGGGCCTCAGCGCCGTCCTGGGCCTGCTGCTGCTGAGGAAGCAGTTTCCTGCACACTACAGGAGAC<br>TCCCTGGAGCTCGTGGTCGGACCCAACTAGGNNNGAGACCGCCACCGAGACCGCCTGGATCTCCTTGGTGACCGCTCTGCATCTAGTGCTGGGCCTCAGCGCCGTCCTGGGCCTGCTGCTGCTGAGGAAGCAGTTTCCTGCACACTACAGGAGAC<br>TCCCTGGAGCTCGTGGTCGGACCCAACTAGGGTGNNNACCGCCACCGAGACCGCCTGGATCTCCTTGGTGACCGCTCTGCATCTAGTGCTGGGCCTCAGCGCCGTCCTGGGCCTGCTGCTGCTGAGGAAGCAGTTTCCTGCACACTACAGGAGAC<br>TCCCTGGAGCTCGTGGTCGGACCCAACTAGGGTGGAGNNNGCCACCGAGACCGCCTGGATCTCCTTGGTGACCGCTCTGCATCTAGTGCTGGGCCTCAGCGCCGTCCTGGGCCTGCTGCTGCTGAGGAAGCAGTTTCCTGCACACTACAGGAGAC<br>TCCCTGGAGCTCGTGGTCGGACCCAACTAGGGTGGAGACCNNNACCGAGACCGCCTGGATCTCCTTGGTGACCGCTCTGCATCTAGTGCTGGGCCTCAGCGCCGTCCTGGGCCTGCTGCTGCTGAGGAAGCAGTTTCCTGCACACTACAGGAGAC<br>TCCCTGGAGCTCGTGGTCGGACCCAACTAGGGTGGAGACCGCCNNNGAGACCGCCTGGATCTCCTTGGTGACCGCTCTGCATCTAGTGCTGGGCCTCAGCGCCGTCCTGGGCCTGCTGCTGCTGAGGAAGCAGTTTCCTGCACACTACAGGAGAC<br>TCCCTGGAGCTCGTGGTCGGACCCAACTAGGGTGGAGACCGCCACCNNNACCGCCTGGATCTCCTTGGTGACCGCTCTGCATCTAGTGCTGGGCCTCAGCGCCGTCCTGGGCCTGCTGCTGCTGAGGAAGCAGTTTCCTGCACACTACAGGAGAC<br>TCCCTGGAGCTCGTGGTCGGACCCAACTAGGGTGGAGACCGCCACCGAGNNNGCCTGGATCTCCTTGGTGACCGCTCTGCATCTAGTGCTGGGCCTCAGCGCCGTCCTGGGCCTGCTGCTGCTGAGGAAGCAGTTTCCTGCACACTACAGGAGAC<br>TCCCTGGAGCTCGTGGTCGGACCCAACTAGGGTGGAGACCGCCACCGAGACCNNNTGGATCTCCTTGGTGACCGCTCTGCATCTAGTGCTGGGCCTCAGCGCCGTCCTGGGCCTGCTGCTGCTGAGGAAGCAGTTTCCTGCACACTACAGGAGAC<br>TCCCTGGAGCTCGTGGTCGGACCCAACTAGGGTGGAGACCGCCACCGAGACCGCCNNNATCTCCTTGGTGACCGCTCTGCATCTAGTGCTGGGCCTCAGCGCCGTCCTGGGCCTGCTGCTGCTGAGGAAGCAGTTTCCTGCACACTACAGGAGAC<br>TCCCTGGAGCTCGTGGTCGGACCCAACTAGGGTGGAGACCGCCACCGAGACCGCCTGGNNNTCCTTGGTGACCGCTCTGCATCTAGTGCTGGGCCTCAGCGCCGTCCTGGGCCTGCTGCTGCTGAGGAAGCAGTTTCCTGCACACTACAGGAGAC<br>TCCCTGGAGCTCGTGGTCGGACCCAACTAGGGTGGAGACCGCCACCGAGACCGCCTGGATCNNNTTGGTGACCGCTCTGCATCTAGTGCTGGGCCTCAGCGCCGTCCTGGGCCTGCTGCTGCTGAGGAAGCAGTTTCCTGCACACTACAGGAGAC<br>TCCCTGGAGCTCGTGGTCGGACCCAACTAGGGTGGAGACCGCCACCGAGACCGCCTGGATCTCCNNNGTGACCGCTCTGCATCTAGTGCTGGGCCTC |

| | | | |
|---|---|---|---|
| | | | AGCGCCGTCCTGGGCCTGCTGCTGCTGAGGAAGCAGTTTCCTGCACACT<br>ACAGGAGAC<br>TCCCTGGAGCTCGTGGTCGGACCCAACTAGGGTGGAGACCGCCACCGA<br>GACCGCCTGGATCTCCTTGNNNACCGCTCTGCATCTAGTGCTGGGCCTC<br>AGCGCCGTCCTGGGCCTGCTGCTGCTGAGGAAGCAGTTTCCTGCACACT<br>ACAGGAGAC<br>TCCCTGGAGCTCGTGGTCGGACCCAACTAGGGTGGAGACCGCCACCGA<br>GACCGCCTGGATCTCCTTGGTGNNNGCTCTGCATCTAGTGCTGGGCCTC<br>AGCGCCGTCCTGGGCCTGCTGCTGCTGAGGAAGCAGTTTCCTGCACACT<br>ACAGGAGAC<br>TCCCTGGAGCTCGTGGTCGGACCCAACTAGGGTGGAGACCGCCACCGA<br>GACCGCCTGGATCTCCTTGGTGACCNNNCTGCATCTAGTGCTGGGCCTC<br>AGCGCCGTCCTGGGCCTGCTGCTGCTGAGGAAGCAGTTTCCTGCACACT<br>ACAGGAGAC<br>TCCCTGGAGCTCGTGGTCGGACCCAACTAGGGTGGAGACCGCCACCGA<br>GACCGCCTGGATCTCCTTGGTGACCGCTNNNCATCTAGTGCTGGGCCTC<br>AGCGCCGTCCTGGGCCTGCTGCTGCTGAGGAAGCAGTTTCCTGCACACT<br>ACAGGAGAC<br>TCCCTGGAGCTCGTGGTCGGACCCAACTAGGGTGGAGACCGCCACCGA<br>GACCGCCTGGATCTCCTTGGTGACCGCTCTGNNNCTAGTGCTGGGCCTC<br>AGCGCCGTCCTGGGCCTGCTGCTGCTGAGGAAGCAGTTTCCTGCACACT<br>ACAGGAGAC<br>TCCCTGGAGCTCGTGGTCGGACCCAACTAGGGTGGAGACCGCCACCGA<br>GACCGCCTGGATCTCCTTGGTGACCGCTCTGCATNNNGTGCTGGGCCTC<br>AGCGCCGTCCTGGGCCTGCTGCTGCTGAGGAAGCAGTTTCCTGCACACT<br>ACAGGAGAC<br>TCCCTGGAGCTCGTGGTCGGACCCAACTAGGGTGGAGACCGCCACCGA<br>GACCGCCTGGATCTCCTTGGTGACCGCTCTGCATCTANNNCTGGGCCTC<br>AGCGCCGTCCTGGGCCTGCTGCTGCTGAGGAAGCAGTTTCCTGCACACT<br>ACAGGAGAC<br>TCCCTGGAGCTCGTGGTCGGACCCAACTAGGGTGGAGACCGCCACCGA<br>GACCGCCTGGATCTCCTTGGTGACCGCTCTGCATCTAGTGNNNGGCCTC<br>AGCGCCGTCCTGGGCCTGCTGCTGCTGAGGAAGCAGTTTCCTGCACACT<br>ACAGGAGAC<br>TCCCTGGAGCTCGTGGTCGGACCCAACTAGGGTGGAGACCGCCACCGA<br>GACCGCCTGGATCTCCTTGGTGACCGCTCTGCATCTAGTGCTGNNNCTC<br>AGCGCCGTCCTGGGCCTGCTGCTGCTGAGGAAGCAGTTTCCTGCACACT<br>ACAGGAGAC<br>TCCCTGGAGCTCGTGGTCGGACCCAACTAGGGTGGAGACCGCCACCGA<br>GACCGCCTGGATCTCCTTGGTGACCGCTCTGCATCTAGTGCTGGGCNNN<br>AGCGCCGTCCTGGGCCTGCTGCTGCTGAGGAAGCAGTTTCCTGCACACT<br>ACAGGAGAC<br>TCCCTGGAGCTCGTGGTCGGACCCAACTAGGGTGGAGACCGCCACCGA<br>GACCGCCTGGATCTCCTTGGTGACCGCTCTGCATCTAGTGCTGGGCCTC<br>NNNGCCGTCCTGGGCCTGCTGCTGCTGAGGAAGCAGTTTCCTGCACACT<br>ACAGGAGAC<br>TCCCTGGAGCTCGTGGTCGGACCCAACTAGGGTGGAGACCGCCACCGA<br>GACCGCCTGGATCTCCTTGGTGACCGCTCTGCATCTAGTGCTGGGCCTC<br>AGCNNNGTCCTGGGCCTGCTGCTGCTGAGGAAGCAGTTTCCTGCACACT<br>ACAGGAGAC<br>TCCCTGGAGCTCGTGGTCGGACCCAACTAGGGTGGAGACCGCCACCGA<br>GACCGCCTGGATCTCCTTGGTGACCGCTCTGCATCTAGTGCTGGGCCTC<br>AGCGCCNNNCTGGGCCTGCTGCTGCTGAGGAAGCAGTTTCCTGCACACT<br>ACAGGAGAC<br>TCCCTGGAGCTCGTGGTCGGACCCAACTAGGGTGGAGACCGCCACCGA<br>GACCGCCTGGATCTCCTTGGTGACCGCTCTGCATCTAGTGCTGGGCCTC<br>AGCGCCGTCNNNGGCCTGCTGCTGCTGAGGAAGCAGTTTCCTGCACACT<br>ACAGGAGAC<br>TCCCTGGAGCTCGTGGTCGGACCCAACTAGGGTGGAGACCGCCACCGA<br>GACCGCCTGGATCTCCTTGGTGACCGCTCTGCATCTAGTGCTGGGCCTC |

| | | | |
|---|---|---|---|
| | | | AGCGCCGTCCTGNNNCTGCTGCTGCTGAGGAAGCAGTTTCCTGCACACT<br>ACAGGAGAC<br>TCCCTGGAGCTCGTGGTCGGACCCAACTAGGGTGGAGACCGCCACCGA<br>GACCGCCTGGATCTCCTTGGTGACCGCTCTGCATCTAGTGCTGGGCCTC<br>AGCGCCGTCCTGGGCNNNCTGCTGCTGAGGAAGCAGTTTCCTGCACACT<br>ACAGGAGAC<br>TCCCTGGAGCTCGTGGTCGGACCCAACTAGGGTGGAGACCGCCACCGA<br>GACCGCCTGGATCTCCTTGGTGACCGCTCTGCATCTAGTGCTGGGCCTC<br>AGCGCCGTCCTGGGCCTGNNNCTGCTGAGGAAGCAGTTTCCTGCACACT<br>ACAGGAGAC |
| L<br>3 | CCCTCACT<br>TCCAGAC<br>CTGCACC<br>GGG | GGCGCTG<br>AGGCCCA<br>GCACTAGA<br>TGC | GCATCTAGTGCTGGGCCTCAGCGCCNNNCTGGGCCTGCTGCTGCTGAG<br>GAAGCAGTTTCCTGCACACTACAGGAGACTGAGGCATGCCCTGTGGCCC<br>TCACTTCCAGACCTGCACCGGG<br>GCATCTAGTGCTGGGCCTCAGCGCCGTCNNNGGCCTGCTGCTGCTGAG<br>GAAGCAGTTTCCTGCACACTACAGGAGACTGAGGCATGCCCTGTGGCCC<br>TCACTTCCAGACCTGCACCGGG<br>GCATCTAGTGCTGGGCCTCAGCGCCGTCCTGNNNCTGCTGCTGCTGAGG<br>AAGCAGTTTCCTGCACACTACAGGAGACTGAGGCATGCCCTGTGGCCCT<br>CACTTCCAGACCTGCACCGGG<br>GCATCTAGTGCTGGGCCTCAGCGCCGTCCTGGGCNNNCTGCTGCTGAG<br>GAAGCAGTTTCCTGCACACTACAGGAGACTGAGGCATGCCCTGTGGCCC<br>TCACTTCCAGACCTGCACCGGG<br>GCATCTAGTGCTGGGCCTCAGCGCCGTCCTGGGCCTGNNNCTGCTGAG<br>GAAGCAGTTTCCTGCACACTACAGGAGACTGAGGCATGCCCTGTGGCCC<br>TCACTTCCAGACCTGCACCGGG<br>GCATCTAGTGCTGGGCCTCAGCGCCGTCCTGGGCCTGCTGNNNCTGAG<br>GAAGCAGTTTCCTGCACACTACAGGAGACTGAGGCATGCCCTGTGGCCC<br>TCACTTCCAGACCTGCACCGGG<br>GCATCTAGTGCTGGGCCTCAGCGCCGTCCTGGGCCTGCTGCTGNNNAG<br>GAAGCAGTTTCCTGCACACTACAGGAGACTGAGGCATGCCCTGTGGCCC<br>TCACTTCCAGACCTGCACCGGG<br>GCATCTAGTGCTGGGCCTCAGCGCCGTCCTGGGCCTGCTGCTGCTGNNN<br>AAGCAGTTTCCTGCACACTACAGGAGACTGAGGCATGCCCTGTGGCCCT<br>CACTTCCAGACCTGCACCGGG<br>GCATCTAGTGCTGGGCCTCAGCGCCGTCCTGGGCCTGCTGCTGCTGAG<br>GNNNCAGTTTCCTGCACACTACAGGAGACTGAGGCATGCCCTGTGGCCC<br>TCACTTCCAGACCTGCACCGGG<br>GCATCTAGTGCTGGGCCTCAGCGCCGTCCTGGGCCTGCTGCTGCTGAG<br>GAAGNNNTTTCCTGCACACTACAGGAGACTGAGGCATGCCCTGTGGCCC<br>TCACTTCCAGACCTGCACCGGG<br>GCATCTAGTGCTGGGCCTCAGCGCCGTCCTGGGCCTGCTGCTGCTGAG<br>GAAGCAGNNNCCTGCACACTACAGGAGACTGAGGCATGCCCTGTGGCCC<br>TCACTTCCAGACCTGCACCGGG<br>GCATCTAGTGCTGGGCCTCAGCGCCGTCCTGGGCCTGCTGCTGCTGAG<br>GAAGCAGTTTNNNGCACACTACAGGAGACTGAGGCATGCCCTGTGGCCC<br>TCACTTCCAGACCTGCACCGGG<br>GCATCTAGTGCTGGGCCTCAGCGCCGTCCTGGGCCTGCTGCTGCTGAG<br>GAAGCAGTTTCCTNNNCACTACAGGAGACTGAGGCATGCCCTGTGGCCC<br>TCACTTCCAGACCTGCACCGGG<br>GCATCTAGTGCTGGGCCTCAGCGCCGTCCTGGGCCTGCTGCTGCTGAG<br>GAAGCAGTTTCCTGCANNNTACAGGAGACTGAGGCATGCCCTGTGGCCC<br>TCACTTCCAGACCTGCACCGGG<br>GCATCTAGTGCTGGGCCTCAGCGCCGTCCTGGGCCTGCTGCTGCTGAG<br>GAAGCAGTTTCCTGCACACNNNAGGAGACTGAGGCATGCCCTGTGGCCC<br>TCACTTCCAGACCTGCACCGGG<br>GCATCTAGTGCTGGGCCTCAGCGCCGTCCTGGGCCTGCTGCTGCTGAG<br>GAAGCAGTTTCCTGCACACTACNNNAGACTGAGGCATGCCCTGTGGCCC<br>TCACTTCCAGACCTGCACCGGG<br>GCATCTAGTGCTGGGCCTCAGCGCCGTCCTGGGCCTGCTGCTGCTGAG<br>GAAGCAGTTTCCTGCACACTACAGGNNNCTGAGGCATGCCCTGTGGCCC<br>TCACTTCCAGACCTGCACCGGG |

| | | | |
|---|---|---|---|
| | | | GCATCTAGTGCTGGGCCTCAGCGCCGTCCTGGGCCTGCTGCTGCTGAG<br>GAAGCAGTTTCCTGCACACTACAGGAGANNNAGGCATGCCCTGTGGCCC<br>TCACTTCCAGACCTGCACCGGG<br>GCATCTAGTGCTGGGCCTCAGCGCCGTCCTGGGCCTGCTGCTGCTGAG<br>GAAGCAGTTTCCTGCACACTACAGGAGACTGNNNCATGCCCTGTGGCCC<br>TCACTTCCAGACCTGCACCGGG<br>GCATCTAGTGCTGGGCCTCAGCGCCGTCCTGGGCCTGCTGCTGCTGAG<br>GAAGCAGTTTCCTGCACACTACAGGAGACTGAGGNNNGCCCTGTGGCCC<br>TCACTTCCAGACCTGCACCGGG<br>GCATCTAGTGCTGGGCCTCAGCGCCGTCCTGGGCCTGCTGCTGCTGAG<br>GAAGCAGTTTCCTGCACACTACAGGAGACTGAGGCATNNNCTGTGGCCC<br>TCACTTCCAGACCTGCACCGGG<br>GCATCTAGTGCTGGGCCTCAGCGCCGTCCTGGGCCTGCTGCTGCTGAG<br>GAAGCAGTTTCCTGCACACTACAGGAGACTGAGGCATGCCNNNTGGCCC<br>TCACTTCCAGACCTGCACCGGG<br>GCATCTAGTGCTGGGCCTCAGCGCCGTCCTGGGCCTGCTGCTGCTGAG<br>GAAGCAGTTTCCTGCACACTACAGGAGACTGAGGCATGCCCTGNNNCCC<br>TCACTTCCAGACCTGCACCGGG |

**Oligos for constructing TpoR variants:**

**1. PolyValine construct:**

| | | | |
|---|---|---|---|
| Long PolyV | Backbone | For | TGGCAGTTTCCTGCACAC |
| | | Rev | GGTCTCGGTGGCGGTCTC |
| | Long PolyV oligo | | AGGGTGGAGACCGCCACCGAGACCGCCTGGGTTGTTGTC GTCGTAGTGGTGGTCGTTGTCGTTGTAGTCGTCGTTGTAG TTGTCGTAGTTGTTgtaGTGTGGCAGTTTCCTGCACACTAC AGG |
| Short PolyV | Backbone | For | TGGCAGTTTCCTGCACAC |
| | | Rev | GGTCACCAAGGAGATCCAGGC |
| | Short PolyV oligo | | ACCGCCTGGATCTCCTTGGTGACCGTGGTGGTCGTTGTC GTTGTAGTCGTCGTTGTAGTTGTCGTAGTTGTTgtaGTGTG GCAGTTTCCTGCACACTACAGG |

**2. TpoR single-residue substituted variants:**

| | | | |
|---|---|---|---|
| A486M | Fragment 1 | Forward oligo | TGTAAAACGACGGCCAGTCTTAAG |
| | | Reverse oligo | GGTCTCCACCCTAGTTGGGTC |
| | Fragment 2 | Forward oligo | GACCCAACTAGGGTGGAGACCATGACCGAGACCGCCTGGATC |
| | | Reverse oligo | GCTATGACCATGTAATACGACTCACTATAGGGG |
| T487A | gift from Andrews Brooks | | |
| T487W | Fragment 1 | Forward oligo | TGTAAAACGACGGCCAGTCTTAAG |
| | | Reverse oligo | GGTCTCCACCCTAGTTGGGTC |
| | Fragment 2 | Forward oligo | GACCCAACTAGGGTGGAGACCGCCTGGGAGACCGCCTGGATC TCC |
| | | Reverse oligo | GCTATGACCATGTAATACGACTCACTATAGGGG |
| E488N | Fragment 1 | Forward oligo | TGTAAAACGACGGCCAGTCTTAAG |
| | | Reverse oligo | GGTGGCGGTCTCCACCCTAG |
| | Fragment 2 | Forward oligo | AGGGTGGAGACCGCCACCAATACCGCCTGGATCTCCTTG |
| | | Reverse oligo | GCTATGACCATGTAATACGACTCACTATAGGGG |
| W491M | Fragment 1 | Forward oligo | TGTAAAACGACGGCCAGTCTTAAG |
| | | Reverse oligo | GGTCTCGGTGGCGGTCTC |
| | Fragment 2 | Forward oligo | ACCGCCACCGAGACCGCCATGATCTCCTTGGTGACCGCT |
| | | Reverse oligo | GCTATGACCATGTAATACGACTCACTATAGGGG |
| S493V | Fragment 1 | Forward oligo | TGTAAAACGACGGCCAGTCTTAAG |
| | | Reverse oligo | GATCCAGGCGGTCTCGGT |
| | Fragment 2 | Forward oligo | ACCGAGACCGCCTGGATCGTGTTGGTGACCGCTCTGCAT |
| | | Reverse oligo | GCTATGACCATGTAATACGACTCACTATAGGGG |
| S493I | Fragment 1 | Forward oligo | TGTAAAACGACGGCCAGTCTTAAG |
| | | Reverse oligo | GATCCAGGCGGTCTCGGT |
| | Fragment 2 | Forward oligo | ACCGAGACCGCCTGGATCATCTTGGTGACCGCTCTGCAT |
| | | Reverse oligo | GCTATGACCATGTAATACGACTCACTATAGGGG |
| S493M | Fragment 1 | Forward oligo | TGTAAAACGACGGCCAGTCTTAAG |

| | | | |
|---|---|---|---|
| | | Reverse oligo | GATCCAGGCGGTCTCGGT |
| | Fragment 2 | Forward oligo | ACCGAGACCGCCTGGATCATGTTGGTGACCGCTCTGCAT |
| | | Reverse oligo | GCTATGACCATGTAATACGACTCACTATAGGGG |
| S493A | Fragment 1 | Forward oligo | TGTAAAACGACGGCCAGTCTTAAG |
| | | Reverse oligo | GATCCAGGCGGTCTCGGT |
| | Fragment 2 | Forward oligo | ACCGAGACCGCCTGGATCGCCTTGGTGACCGCTCTGCAT |
| | | Reverse oligo | GCTATGACCATGTAATACGACTCACTATAGGGG |
| S493F | Fragment 1 | Forward oligo | TGTAAAACGACGGCCAGTCTTAAG |
| | | Reverse oligo | GATCCAGGCGGTCTCGGT |
| | Fragment 2 | Forward oligo | ACCGAGACCGCCTGGATCTTCTTGGTGACCGCTCTGCAT |
| | | Reverse oligo | GCTATGACCATGTAATACGACTCACTATAGGGG |
| L502T | Fragment 1 | Forward oligo | TGTAAAACGACGGCCAGTCTTAAG |
| | | Reverse oligo | CACTAGATGCAGAGCGGTC |
| | Fragment 2 | Forward oligo | GTGACCGCTCTGCATCTAGTGACCGGCCTCAGCGCCGTCCTG |
| | | Reverse oligo | GCTATGACCATGTAATACGACTCACTATAGGGG |
| L504K | Fragment 1 | Forward oligo | TGTAAAACGACGGCCAGTCTTAAG |
| | | Reverse oligo | CACTAGATGCAGAGCGGTCACC |
| | Fragment 2 | Forward oligo | GCTCTGCATCTAGTGCTGGGCAAGAGCGCCGTCCTGGGCCTG |
| | | Reverse oligo | GCTATGACCATGTAATACGACTCACTATAGGGG |
| S505M | Fragment 1 | Forward oligo | TGTAAAACGACGGCCAGTCTTAAG |
| | | Reverse oligo | GCCCAGCACTAGATGCAGAGC |
| | Fragment 2 | Forward oligo | CTGCATCTAGTGCTGGGCCTCATGGCCGTCCTGGGCCTGCTG |
| | | Reverse oligo | GCTATGACCATGTAATACGACTCACTATAGGGG |
| L508V | Fragment 1 | Forward oligo | TGTAAAACGACGGCCAGTCTTAAG |
| | | Reverse oligo | GACGGCGCTGAGGCCCAG |
| | Fragment 2 | Forward oligo | CTGGGCCTCAGCGCCGTCGTTGGCCTGCTGCTGCTGAGG |
| | | Reverse oligo | GCTATGACCATGTAATACGACTCACTATAGGGG |
| L508I | Fragment 1 | Forward oligo | TGTAAAACGACGGCCAGTCTTAAG |
| | | Reverse oligo | GACGGCGCTGAGGCCCAG |
| | Fragment 2 | Forward oligo | CTGGGCCTCAGCGCCGTCATTGGCCTGCTGCTGCTGAGG |
| | | Reverse oligo | GCTATGACCATGTAATACGACTCACTATAGGGG |
| L510C | Fragment 1 | Forward oligo | TGTAAAACGACGGCCAGTCTTAAG |
| | | Reverse oligo | GCCCAGGACGGCGCTGAG |
| | Fragment 2 | Forward oligo | CTCAGCGCCGTCCTGGGCTGCCTGCTGCTGAGGTGGCAG |
| | | Reverse oligo | GCTATGACCATGTAATACGACTCACTATAGGGG |

**Oligos for next-generation sequencing:**

**1. Oligos for reverse transcription oligos and cDNA amplification**

TpoR

| Library | Reverse Transcription oligo | Amplify cDNA | |
|---|---|---|---|
| | | Reverse oligo | Forward oligo |
| L1 | CTGAGACTTGCACATCGCAGCNNNNNNNNNNNNNNNNNTCTCCTGTAGTGTGCAGG | CTGAGACTTGCACATCGCAGC | GTGACCTATGAACTCAGGAGTCACTAGGGTGGAGACCGCC |
| L2 | CTGAGACTTGCACATCGCAGCNNNNNNNNNNNNNNNNNGGAAACTGCCACCTCAG | CTGAGACTTGCACATCGCAGC | GTGACCTATGAACTCAGGAGTCGAGCTCGTGGTCGGAC |
| L3 | CTGAGACTTGCACATCGCAGCNNNNNNNNNNNNNNNNNGTGCAGGTCTGGAAGTG | CTGAGACTTGCACATCGCAGC | GTGACCTATGAACTCAGGAGTCGTGCTGGGCCTCAGC |
| L4 | CTGAGACTTGCACATCGCAGCNNNNNNNNNNNNNNNNNGTGCAGGTCTGGAAGTG | CTGAGACTTGCACATCGCAGC | GTGACCTATGAACTCAGGAGTCGAGCTCGTGGTCGGAC |

EpoR

| Library | Reverse Transcription oligo | Amplify cDNA | |
|---|---|---|---|
| | | Reverse oligo | Forward oligo |
| L1 | CTGAGACTTGCACATCGCAGCNNNNNNNNNNNNNNNNGAGAGCAGCGCGAG | CTGAGACTTGCACATCGCAGC | GTGACCTATGAACTCAGGAGTCCTGGAGCGCCTGGTC |
| L2 | CTGAGACTTGCACATCGCAGCNNNNNNNNNNNNNNNNGGTGAAGAGGCCTTCAAACTC | CTGAGACTTGCACATCGCAGC | GTGACCTATGAACTCAGGAGTCGCTCTCCCTCATCCTCGTG |
| L3 | CTGAGACTTGCACATCGCAGCNNNNNNNNNNNNNNNNGGTGAAGAGGCCTTCAAACTC | CTGAGACTTGCACATCGCAGC | GTGACCTATGAACTCAGGAGTCCTGGAGCGCCTGGTC |

**2. Oligos structure for indexing amplicons**
**Read 1 end primer:**
(5′)AATGATACGGCGACCACCGAGATCTACACTCTTTCCCTACACGACGCTCTTCCGATCTZZZZZZZZGTGACCTATGAACTCAGGAGTC(3')

**Read 2 end primer:**
(5′)CAAGCAGAAGACGGCATACGAGATCGGTCTCGGCATTCCTGCTGAACCGCTCTTCCGATCTZZZZZZZZCTGAGACTTGCACATCGCAGC(3')
